# Genetic dissection of Mycobacteriophage D29 host lysis reveals two lysis regulators and a novel lipoprotein that regulate the lysis event and are localized to distinct regions of the genome

**DOI:** 10.64898/2026.08.27.747656

**Authors:** Richard S. Pollenz, Makenzie Davenport, Kira Ruiz-Houston

## Abstract

Phage D29 infects *Mycobacterium smegmatis* mc^2^ 155 and has a non-canonical lysis cassette that encodes two endolysin enzymes (Lysin A and Lysin B) and a single two transmembrane domain (TMD) protein, LysA2a similar to F1 cluster phage LysF1a. A 1TMD LysF1b homolog, LysA2b, is encoded by a gene found downstream of the *tape measure*. Exogenous expression of both LysA2 proteins in tandem is a cytotoxic to *M. smegmatis*. Deletion of *lysA2a* produces phages that are lysis competent with a 10-minute triggering delay and 30% plaque size reduction. Deletion of *lysA2b* results in severe lysis defects manifest by 70% reduced plaque size, delayed lysis timing and reduced burst size. Deletion of both *lysA2* genes results in phages that are viable and show lysis phenotypes like the *lysF1b* deletion. Genetic complementation of *lysA2b* deleted phage with the *lysF1b* gene fully complements the lysis phenotypes but alters the triggering time to that of an F1 cluster phage. Energy poisons trigger lysis prematurely in all phages with *lysA2* gene deletions. Lysis recovery mutants (LRM) isolated from phages lacking the *lysA2b* genes generate wild type plaque size and have point mutations that map to TMD1 or the C-terminal region of the *lysA2a* gene. LRMs isolated from phages lacking both *lysA2* genes show premature lysis and have mutations that all map to residue C31 of a novel lipoprotein (gene *64*). Deletion of gene *64* does not change wild type D29 lysis phenotypes or rescue the lysis defects of any of the *lysA2* mutants. A fitness/competition assay shows that loss of the *lysA2* genes imposes a substantial competitive fitness cost. These finding support a lysis regulatory network model where the 2TMD protein is maintained in an inactive state until activated by its cognate 1TMD lysis regulator and the lipoprotein has accessory function that may enhance lysis efficiency.

## INTRODUCTION

Bacteriophages are viruses that infect bacteria generally with high host specificity. This property makes them promising agents to combat the growing number of antibiotic resistant bacterial strains impacting human health [reviewed in: 1, 2]. In addition, other sectors such as the food industry, agriculture, and environment are evaluating the use of phages to combat specific bacterial strains [3–4]. Across all of these applications, the utility of the phage is based on its ability to ultimately lyse and destroy the bacteria. Thus, understanding how the phage lysis pathway is mediated across phages that infect diverse hosts may help in isolating and engineering phages for specific applications.

The most well studied phage host lysis pathways are from phages that infect Gram negative *E coli*. Numerous genetic and biochemical studies have established a general three-step model for the triggering of the lysis pathway that involves three distinct sets of proteins. The first are proteins with 1-4 transmembrane domains (TMDs). These proteins termed, holins or antiholins are encoded by the same gene is some phages and work together to, 1) control the timing of the lysis event following phage infection [5, 6], 2) destabilize the membrane proton motive force (PMF) [5, 6] and 3) in the canonical Type I holin model, generate large 1μm “pores” in the inner membrane (IM) to allow the exit of the endolysin enzymes to the periplasmic space where they can access and cleave the PG layer [7, 8]. The second are the endolysin/s that have defined catalytic activity to lyse the peptidoglycan (PG) layer [9]. Finally, there are secreted membrane proteins and lipoproteins termed spanins that are associated with the IM and outer membrane (OM). There proteins interact after the PG layer is cleaved to generates the lesion across the IM and OM that allows for the efficient release of the phage progeny [10–12]. What is fascinating from the rigorous body of work on phages lambda, P1, T4, T7 and Mu is that each appear to use a similar set of proteins (endolysins, holins, antiholins and spanins) to complete the lysis pathway through distinct molecular mechanisms [reviewed in: 13-15]. With a predicted 10^31^ phages in the biosphere [16] and 968,661 holins currently archived in GenBank, it is likely that additional variations on these pathways are yet to be discovered.

Rigorous genetic and biochemical studies of host lysis by phages that infect Gram positive hosts are much less plentiful, but several recent studies have provided insight into the lysis pathway. The first evaluated several F1 cluster phages that infect *Mycobacterium smegmatis* [17]. These phages have a defined lysis cassette that encodes Lysin A, Lysin B and two holin-like TMD lysis regulatory proteins termed LysF1a (2TMD) and LysF1b (1TMD). Genetic deletion of the *lysF1* genes either individually or in tandem was not lethal to phage propagation but produced significant lysis phenotypes including reduced plaque size, reduced lysis in liquid media, reduced lysis timing and reduced burst size. The lysis phenotypes were most pronounced in phages with the 1TMD *lysF1b* deletion and the LysF1b protein represents a new class of lysis regulator that does not fit the Type III holin model and appears to be found predominantly in phages that infect Gram positive hosts. Screening of phages with *lysF1b* deletions identified lysis recovery mutants (LRMs) that had various point mutations in the lysF1a open reading frame that generated a LysF1a protein that was lysis competent in the absence of the LysF1b. The lysF1a and LysF1b proteins are not like the holin/antiholin system in phage lambda and phage 21 where the holin proteins are encoded by the same gene and triggering converts the antiholin to an active holin [10–12]. In the F1 phages, both lysis regulatory proteins are distinct proteins, and triggering does not convert the 2TMD LysF1a protein to an active regulator. These results suggest a novel model in which the LysF1b is the main lysis regulator and its expression activates the LysF1a.

A second study evaluated phages that infect the Gram-positive *Corynebacterium glutamicum* [18]. Like the F1 cluster phages, *Corynebacterium* phages Cog and CL31 have a defined lysis cassette, but in these phages, there is a single endolysin and three different TMD proteins. One termed LysZ is a 177 amino acid protein with a single TMD. Like the F1 cluster phages, deletion of *lysZ* from phages CL31 or Cog results in a small plaque phenotype. However, the phenotype can be rescued if phages are plated on a *Corynebacterium glutamicum* strain that is defective in the synthesis of lipoarabinomannan (LAM). Thus, it is proposed that the LysZ is topologically localized with its extensive C-terminus outside the *Corynebacterium* IM, and functions more like a spanin during the host lysis event. The function of the other TMD proteins in the lysis of *C. glutamicum* has not been resolved, but they are predicted to provide the “holin” function to theses phages. Although the LysF1b and LysZ both are 1TMD proteins, the LysF1b is 50 aa smaller, shares <20% amino acid identity and is topologically predicted to have the opposite membrane orientation. Thus, the results support these proteins having distinct functions in the lysis pathway in their respective bacterial hosts.

This current study expands on the F1 cluster phage work and aims to genetically evaluate the lysis pathway of A2 cluster phage D29 that infects *Mycobacterium smegmatis* mc^2^ 155. D29 is a key Actinobacteriophage and has relevance to phage therapy since it has host range to *Mycobacterium tuberculosis* [19–21]. Phage D29 encodes predicted homologs to the LysF1a and LysF1b proteins, but does not encode them from a canonical lysis cassette but from distinct regions of the genome. Experiments utilizing recombineered D29 phages and multiple lysis assays under physiological growth conditions will be used to explore the hypothesis that these proteins function in host lysis in a similar manner to those from F1 cluster phages.

## MATERIALS AND METHODS

### Bioinformatic analysis of transmembrane domains and AlphaFold3 analysis

The identification of transmembrane domains (TMD) and protein topologies was carried out essentially as described by Pollenz et al., 2022 [22] utilizing Deep TMHMM (v1.0.57) [23, TOPCONS (v2.0) [24], SignalP-6.0 [25, 26], and HHpred (databases: PDB_mmCIF70_20_Feb; Pfam-A_v38.2; NCBI_Conservd_Domains(CD)_v3.19) [27]. All membrane proteins described in this report had 100% consensus predictions of TMD domain location, number and protein topology in all programs. Structural predictions of proteins and protein-protein interactions were determined using AlphFold3 [28]. For single proteins, pTM scores >50% indicate high confidence in the model predictions. For protein-protein interactions, ipTM scores <90% are considered low confidence and should be used with caution [28]. Pairwise Sequence Alignments were performed using Emboss Needle [29] and Multiple Sequence Alignments were performed using Clustal Omega [29].

**Table 1.**
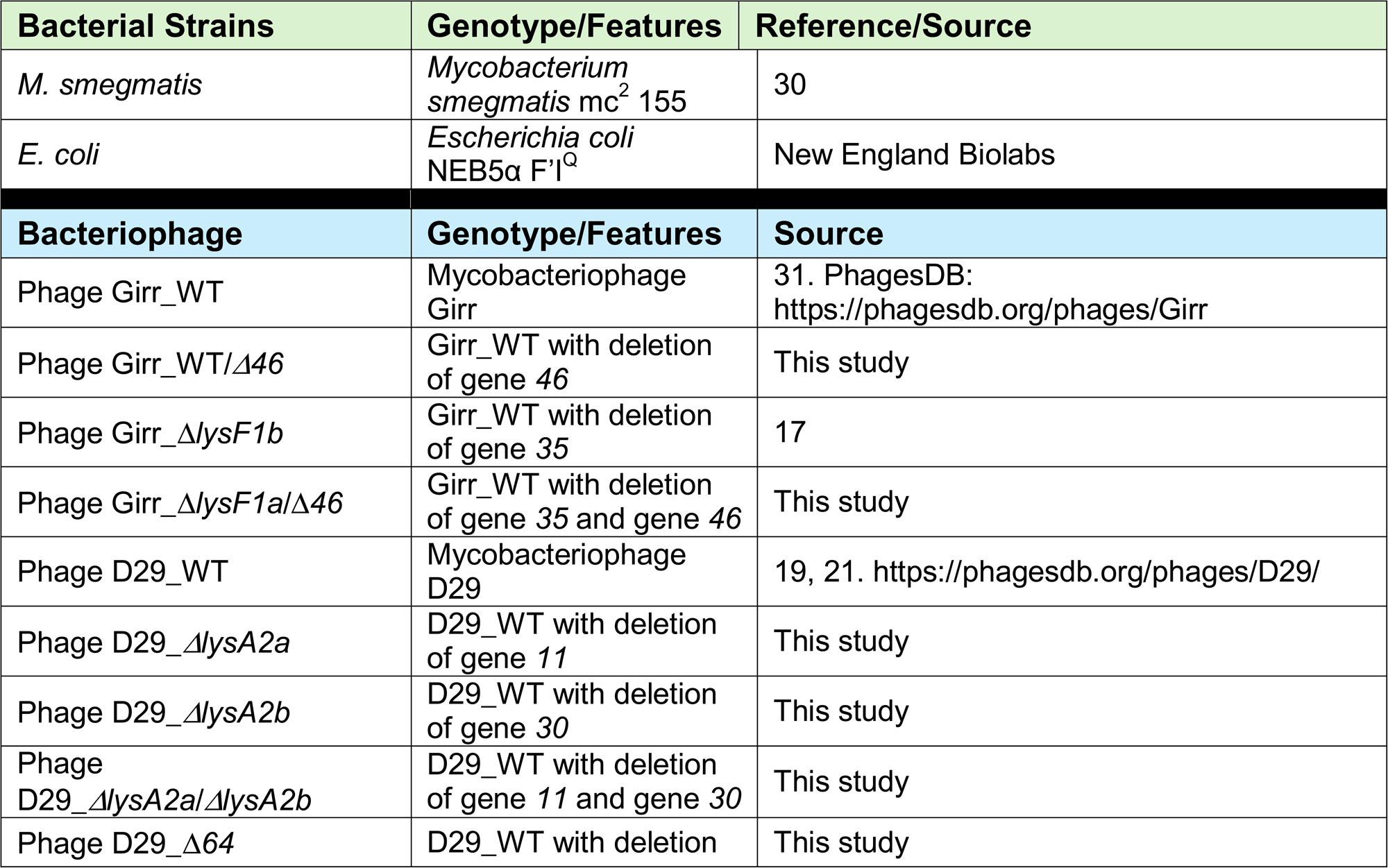
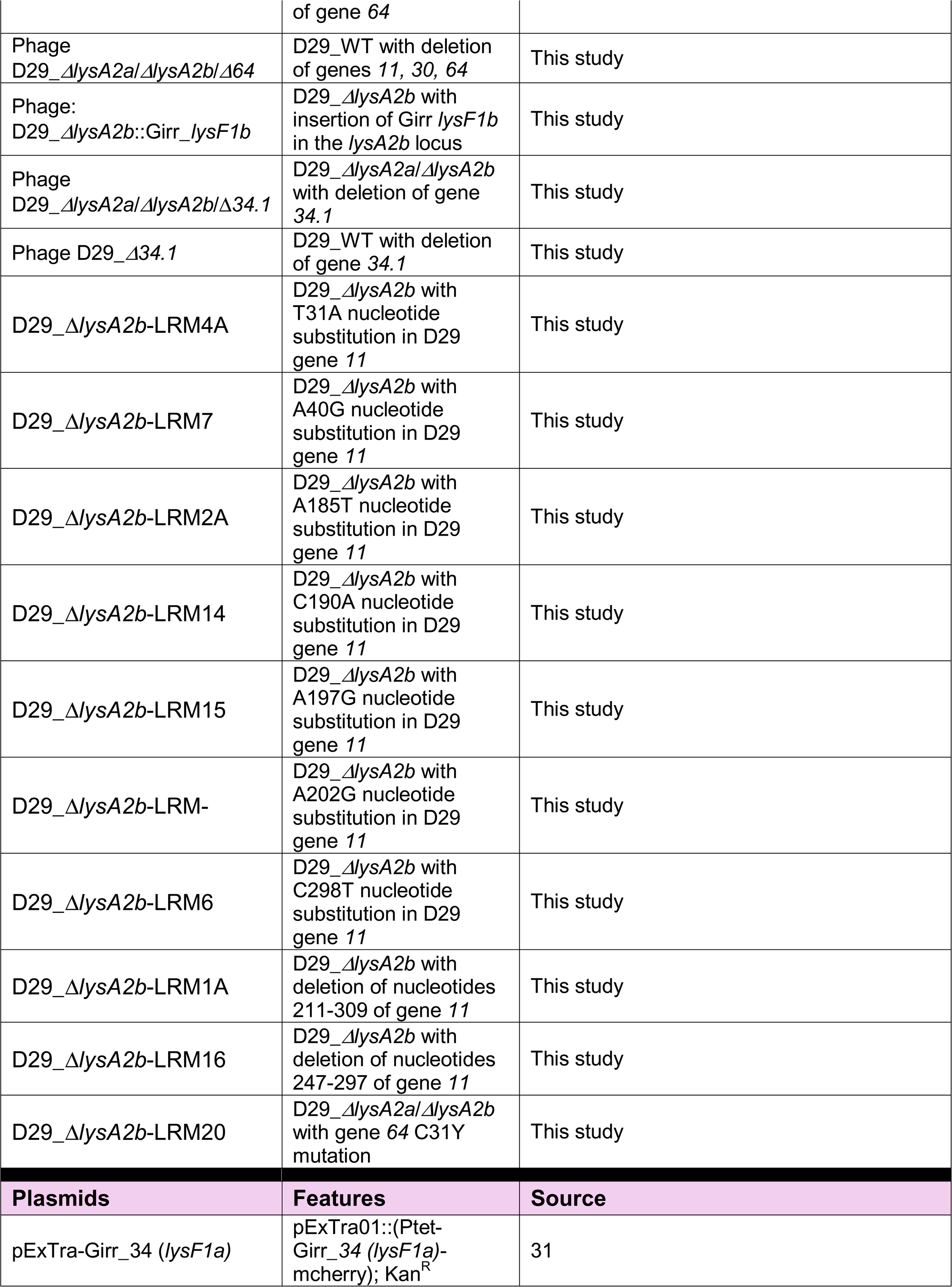
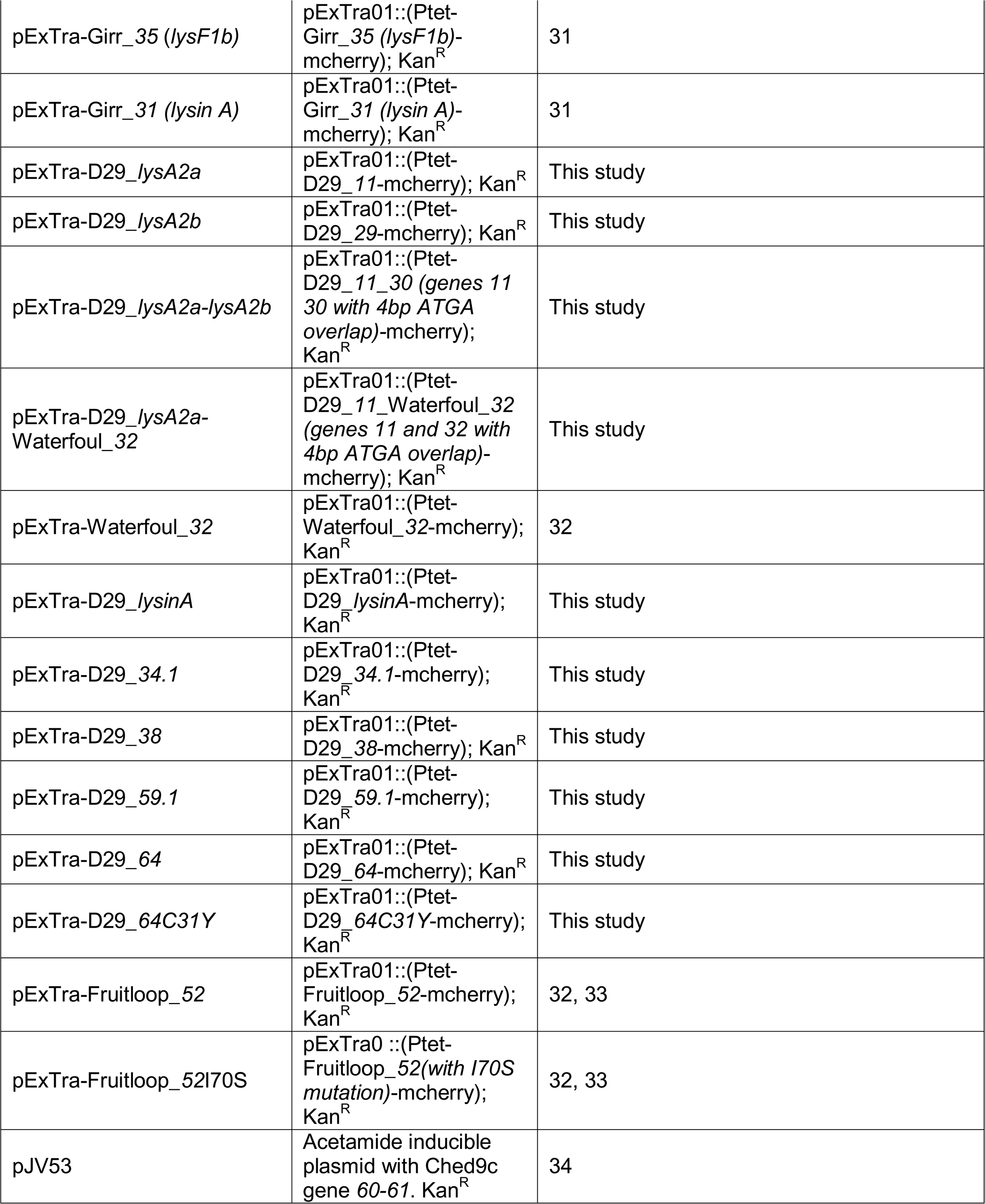
Bacterial Strains, Phages and Plasmids.

### Bacterial strains, growth, plaque assay and lysogeny

The list of bacterial strains, phage and plasmids utilized in this study is presented in Table 1. *Mycobacterium smegmatis* mc^2^155 [30] was used as the wild type host for all phage related experiments and propagated on 7H10 agar as described previously [17]. To generate saturated stock cultures for daily use, a single colony of *M. smegmatis* was propagated in 5ml of Middlebrook 7H9 media supplemented with 10% Albumen Dextrose Complex (ADC; 5% albumen, 2% dextrose, 154mM NaCl), glycerol (0.5%) Tween 80 (0.05%) with shaking (225rpm) at 37°C for 18 hrs. To generate the cultures for liquid lysis experiments, 500ul of saturated stock was centrifuged at 500rpm for 1 minute and 100ul of the supernatant added to 250ml baffled flasks containing 55ml of supplemented 7H9 without Tween. Cultures were shaken at 225rpm for ∼18 hrs at 37°C to an OD_600_ <0.8. Prior to use in liquid lysis studies, the culture was centrifuged at 500rpm for 1 minute to pellet any aggregated cells and the cells in the supernatant diluted to in supplemented 7H9 without Tween to an OD_600_ of ∼0.25 prior to phage infection. Cultures were infected at a multiplicity of infection (MOI) of 10 and changes in OD_600_ evaluated as detailed previously [17]. Plaque assays were completed as detailed previously [17]. Competent *M. smegmatis* cells for recombineering or for plasmid transfection and complementation studies were prepared as described previously [32, 35, 36]. *Escherichia coli* NEB5α F′I^Q^ (New England Biolabs) was used for plasmid amplification and purification and grown in LB broth or on LB agar supplemented with 50ug/ml kanamycin sulfate. To generate lysogens, 100ul of phage stock at 1 x 10^9^ PFU/ml was spread on 7H10 agar plates and then 100ul of saturated *M. smegmatis* dilutions from 1 x10^-2^ to 1 x 10^-7^ were spread on the plates. Plates were incubated at 37°C for 3-5 days. Plates with no phage served as controls. Surviving colonies (lysogens) were evaluated as described in Wise *et al*., 2026 [36] and the efficiency of lysogeny (EOL) determined by comparing the number of lysogens to the number of colonies on the plates without phage.

### PCR screening of plaques and generation of pure phage filtrates

Individual plaques were picked from plaque assay plates using a p1000 pipette tip to “core-out” the center of the plaque. The core was suspended in 120µl of phage buffer (10 mM Tris-HCl, pH 7.5; 10 mM MgSO_4_, 68.5 mM NaCl; 1mM CaCl_2_), vortexed, and incubated at 22°C for 2-8 hrs. 1ul of the sample was utilized as a template to evaluate the presence of phage using the primer sets indicated. To generate purified phage filtrates, the PCR verified plaque pick sample was serial diluted and plaque assays performed. Plates with ∼50 plaques were tested by picking all plaques and using PCR to verify the purity of the population. Webbed plates from the same dilution series were flooded with 5ml of phage buffer, incubated on a rocker platform at 22°C for 5 hrs. and then filtered through sterile 0.2μ filter units to generate the purified high titer filtrate. Phage titer was determined by serial diluting phage filtrates and counting plaques.

### Phage Fitness/Competition assay

A 1ml sample of D29_WT and D29_Δ*lysA2a*/Δ*lysA2b*/Δ*64* phage, each at ∼5 x10^9^ PFU/ml was prepared. The 1:1 stock was serial diluted and plaque assays performed. Plates with ∼50 plaques were evaluated by PCR as described above to validate that the phage population was ∼1:1. Plates from the 1:1 stock with ∼200 total plaques were then flooded and filtrates prepared as detailed above. Filtrates were serail diluted, individual plaques evaluated by PCR and webbed plates flooded to prepare filtrates. This plating/harvesting strategy was repeated a total of 4 times. The ratio of D29_WT to D29_Δ*lysA2a*/Δ*lysA2b*/Δ*64* was determined for each filtrate by PCR analysis of individual plaques on plates with 200-350 plaques. Filtrates were also evaluated by DADA PCR using the D29-DADA4_Δ*lysA2b* primer as detailed previously [17] and in S2 Figure 1 in the S2 File to detect the presence of the D29_Δ*lysA2a*/Δ*lysA2b*/Δ*64* phage in the D29_WT background.

**Figure 1.**
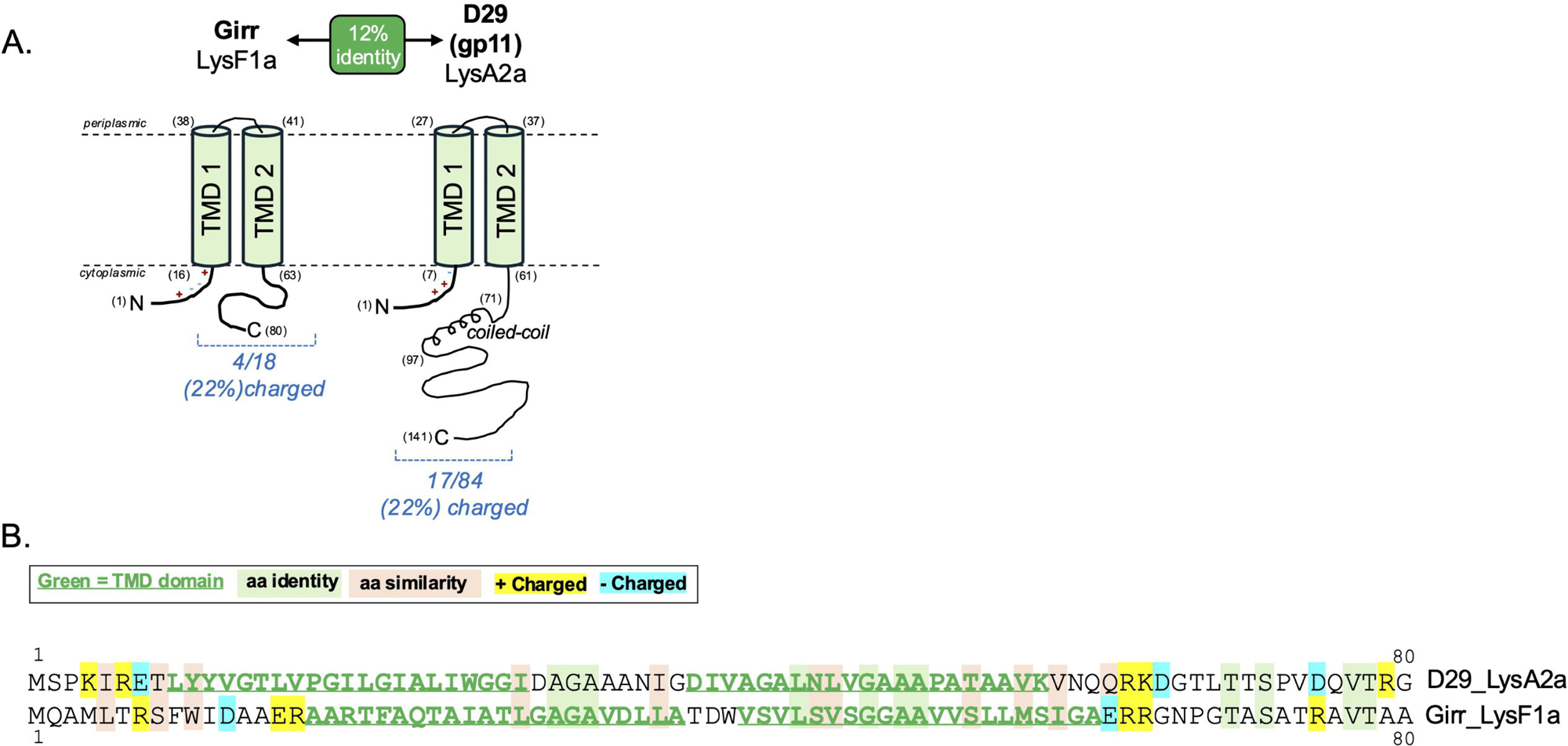
Amino acid sequence and topology of LyA2a and LysF1a. **A.** The amino acid sequences of full length Girr LysF1a and the first 80 amino acids of D29 LysA2a showing the location of the two TMDs. The key to the highlighted residues is shown. **B.** Predicted membrane topology of Girr LysF1a and D29 LysA2a showing the extended C-terminal domain of LysA2a and the location of the coiled-coil.

### Plasmid constructs and *M. smegmatis* transformation

All PCR primers utilized in this study are shown in S1 Table 1 in the S1 File. To generate the inducible expression plasmids the indicated genes were PCR amplified from phage stocks with Q5 DNA polymerase (New England Biolabs) using gene specific primers containing pExTra homology regions (Table S1 in S1 File) as detailed previously (32). Purified PCR products (were ligated into linearized pExTra01 using isothermal assembly (NEB HiFi 2x Master Mix) as described [31, 32, 36]. Recombinant plasmids were recovered by transformation of *E. coli* NEB5α F′I^Q^ and colonies selected on LB agar supplemented with 50μg/µl kanamycin sulfate. Plasmid DNA was purified using the GeneJet kit (Fisher) and all plasmids were sequence-verified by Sanger sequencing using the pExTra Seq primer (Azenta).

### Cytotoxicity assay

Cytotoxicity assays were completed as describe previously [31, 32, 36]. Briefly, colonies of *M. smegmatis* transformed with pExTra constructs were suspended in 7H9 media, serial diluted and spotted onto 7H10 agar supplemented with kanamycin (10ug/ml) with and without 100ng/ml anhydrotetracycline (aTc). Plates were propagated at 37°C for 4-5 days. Cytotoxicity phenotypes were evaluated by comparing the bacterial growth on the control and aTc plates and comparing to the positive and negative controls. To visualize the mCherry expression, plates were inverted on a UV box and photographed. Fruitloop gene *52* and the Fruitloop *52* I70S mutant served as positive and negative controls as described previously [31–33, 36].

### Bacteriophage recombineering and genome sequencing

The deletion and insertion of genes in phage D29 and Girr were carried out using the Bacteriophage Recombineering of Electroporated DNA (BRED) procedure as originally described [34, 37], with the modifications detailed in Pollenz *et al*., 2026 [17]. All gene deletions were validated by PCR using DNA oligonucleotide primers that flanked the deletion (Table S1 in S1 File). Gene insertions were validated by PCR using DNA oligonucleotide primers that flanked the insertion and also primers that were internal to the added gene (Table S1 in S1 File). Sanger sequencing of the amplified PCR products was used to validate the insertion and deletion regions of the upstream and downstream genes (Azenta). Pure populations of mutant phage were evaluated by full genome sequencing as described [38, 39], to validate that there were no point mutations outside of the deleted or inserted genes.

### Plaque size analysis

Plaques were generated using the plaque assay procedure described above. For experiments where plaque size would be evaluated between several different phages, identical bacterial samples, incubation times, phage concentrations and top agar amounts were utilized. Plaque size was evaluated using ImageJ software [40] and by manual measurement using photos embedded in PowerPoint and custom rulers. In most experiments, results are presented for an average of at least 20 plaques and the statistical significance of measurements determined by unpaired t-test (https://www.graphpad.com/quickcalcs/ttest1/). Plaque volume was calculated by the formula πr^2^h where r is the radius of the plaque and h was set to 1mm for the top agar thickness.

### Liquid lysis assay and cell viability assay from liquid cultures

Liquid lysis assays were carried out essentially as described by Payne et al., 2009 [41] with the modifications described in Pollenz *et. al.* 2026 [17]. All liquid lysis experiments utilized multiplicity of infection (MOI) of 10. In some experiments, KCN (1M stock in water), was added directly to the shaking culture to produce a final concentration of 10mM. All liquid lysis experiments were repeated at least three separate times and representative experiments are presented. To assess if there were viable cells surviving in the phage infected cultures, an aliquot was removed from the infected cultures at the times specified in the results and 5ul spotted directly without dilution on supplemented 7H10 agar. Bacterial growth was evaluated after incubation at 37°C for 36-48 hours. All phage infected samples were compared to the non-infected *M. smegmatis* control.

### One-step growth curves and burst size determination

One-step growth curves and burst size were determined using the technique outlined in detail in Pollenz *et. al.* 2026 [17]. Briefly, fresh overnight cultures of *M. smegmatis* were centrifuged and set to an OD_600_ of 4.0. 250ul of the culture (an effective OD_600_ of 1.0 = ∼7 x 10^7^ CFU) was infected with 7 x 10^7^ phages (MOI = 1.0) for 30 minutes at 37°C, mixed with 750ul 7H9 and then centrifuged at 6,200 rpm for 4 minutes to pellet the cells. Infected cells were then suspended in 1.0ml 7H9, 7ul added to 70ml supplemented 7H9 and shaken at 225rpm at 37°C. To assess infective centers, aliquots of the diluted culture were immediately used for plaque assay. The liberation of phage in the culture was determined by removing an aliquot of media from the shaking culture at the specified times, immediately passing it through a 0.2µ syringe filter to remove cells and then completing plaque assays. The number of plaques on the plates were counted and burst size at all time points could be directly determined by dividing the total phage in the 70ml culture by the number of infectious centers at the start of the experiment. All phages were evaluated through a minimum of three separate assays.

## RESULTS AND DISCUSSION

### Cluster A phages have a non-canonical genomic organization of genes encoding putative lysis enzymes and transmembrane lysis regulators

Mycobacteriophages in the Actinobacteriophage Database (PhagesDB.org) are assigned to clusters primarily based on gene content similarity [42–44]. Of the approximately 2,800 annotated phages that infect *Mycobacterium smegmatis*, members are distributed among 36 clusters ranging in size from two to more than 800 phages. Previous genetic and bioinformatic analyses of putative lysis genes demonstrated that approximately 81% of these clusters encode a canonical lysis cassette located downstream of the tape measure and minor tail genes. These cassettes typically contain genes encoding Lysin A, Lysin B, and homologs of the 1-transmembrane domain (1TMD) and 2-transmembrane domain (2TMD) lysis regulators LysF1b and LysF1a, respectively, first characterized in the F1 cluster phages Girr and NormanBulbieJr (NBJ) [17].

A notable exception to phages with canonical lysis cassettes are the Cluster A mycobacteriophages. Cluster A is the largest group of mycobacteriophages in PhagesDB, comprising more than 800 phages (approximately 29% of all annotated *M. smegmatis* phages) and is subdivided into 22 subclusters (A1–A22). Although all Cluster A phages encode a Lysin A homolog, the gene occupies a distinctly non-canonical position that is upstream of the tape measure gene within the structural gene region in the 5′ arm of the genome (S3 Figure 2A, S4 Figure 2B and S5 Figure 2C in S2 File). The *lysin A* gene is often, but not always adjacent to a gene encoding Lysin B. Furthermore, the *lysin* genes are either adjacent to a gene encoding a 2TMD LysF1a homolog or is not associated with any predicted transmembrane lysis regulator. This genomic organization (and lack of multiple TMD genes) contrasts sharply with the defined lysis cassettes observed in most clusters of actinobacteriophages [17, 22, 31, 45-47].

**Figure 2.**
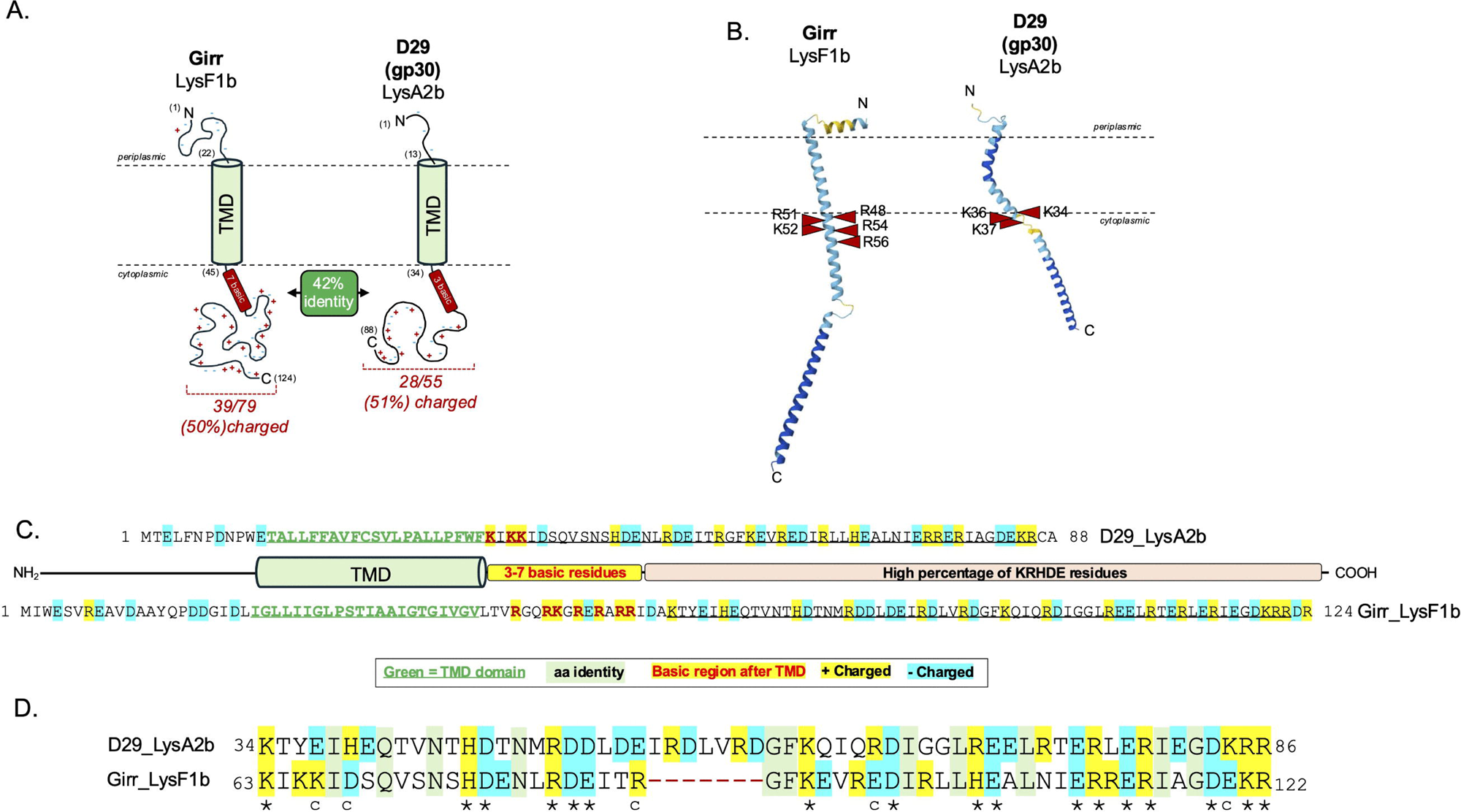
Amino acid sequence and topology of LyA2b and LysF1b. **A.** The amino acid sequences of Girr LysF1b and D29 LysA2b LysF1b showing the location of the TMD. The key to the highlighted residues is shown. **B.** Pairwise sequence alignment of the conserved C-terminal region. Identical charged residues are indicated with an asterisk and conserved charged residues indicated with a c. **C.** Schematic showing the predicted membrane topology and highly charged C-terminal domain. **D.** AlphaFold3 structural prediction of LysF1b (pTM 0.40) and LysA2bp (pTM 0.42). The location of the basic residues after the TMD are shown.

To determine whether additional lysis regulators were encoded elsewhere, all predicted proteins from representative phages in each A Cluster were analyzed for transmembrane domains and topological similarity to the experimentally characterized LysF1a and LysF1b proteins as detailed in Methods. S3 Figure 2A, S4 Figure 2B and S5 Figure 2C in the S2 File, shows the predicted lysis gene organizations across the A cluster phages. What is noteworthy, is that all cluster A phage encode a 1TMD LysF1b homolog; however, unlike canonical lysis cassettes, these genes are not adjacent to the *lysin A* gene but instead are located downstream of the *tape measure* gene embedded within the various *minor tail* genes. In the A cluster phages that do not have a *lysF1a-like* gene adjacent to the *lysin A*, the gene is found adjacent to the *lysF1b-like* gene. This general location (downstream of the *tape measure* and *minor tails*) corresponds to the genomic region where canonical lysis cassettes are typically found in other actinobacteriophages that have genomes of <100,000bp [17, 22, 31, 45-47]. Thus, all Cluster A phages encode a 2TMD LysF1a homolog that is located either adjacent to the 1TMD gene or adjacent to *lysin A*. Phages belonging to subclusters A2, A4, A7, and A10 also encode an additional predicted 4TMD protein immediately upstream of the *lysF1b-like* gene (S3 Figure 2A, S4 Figure 2B and S5 Figure 2C in S2 File).

Comparative analysis of the encoded LysF1a and LysF1b homologs further revealed substantial diversity among the predicted lysis regulators in the A cluster phages. The 2TMD LysF1a homologs are distributed among five distinct phams, none of which include the Girr LysF1a protein. In contrast, the 1TMD LysF1b homologs are grouped into four phams, with the majority of A cluster phages encoding a 1TMD protein grouped to the same pham as the experimentally characterized Girr LysF1b (Pham 297967; PhagesDB release June 2026).

Collectively, these analyses demonstrate that, despite their compact genomes of < 60,000bp, Cluster A phages have evolved a non-canonical genomic organization in which the lysis enzymes and their putative transmembrane regulators are encoded in distinct genomic regions rather than within a single conserved lysis cassette. This atypical genomic organization led us to hypothesize that the spatial separation of the lysis enzymes and transmembrane regulators reflects a fundamentally different mechanism of lysis control in Cluster A phages. To test this hypothesis, we used the A2 cluster phage D29 as a genetically tractable model to define the functions of its predicted transmembrane lysis regulators and to identify additional factors contributing to lysis timing.

### Membrane topology of putative lysis regulator (Lys) proteins encoded by A2 cluster phage D29

To investigate the functions of the predicted transmembrane lysis regulators encoded by Cluster A phages, we selected mycobacteriophage D29 as a genetically tractable model. D29 is an extensively studied A2 cluster phage that infects both *Mycobacterium smegmatis* and *Mycobacterium tuberculosis*, making it an important experimental model and a promising candidate for phage therapy [19–21]. In addition, the two D29 lytic enzymes, Lysin A (gp10) and Lysin B (gp12), have been characterized extensively and are being evaluated as potential antimicrobial agents [48–52].

As shown in the previous section, bioinformatic analysis identified two D29 proteins with strong structural and topological similarity to the experimentally characterized F1 cluster lysis regulators LysF1a and LysF1b. Although these proteins are encoded in distinct regions of the D29 genome rather than within a canonical lysis cassette, their predicted membrane topologies suggest that they may function as lysis regulators.

D29 gene *11* encodes a 141-amino-acid protein containing two predicted transmembrane domains with an N-in/C-in membrane topology, closely resembling the topology of the 2TMD lysis regulator LysF1a from the F1 cluster phages Girr and NBJ (Figure 1A). Unlike the 80-amino-acid LysF1a proteins, gp11 possesses an extended C-terminal cytoplasmic domain containing a predicted coiled-coil region spanning residues 71–97 that may participate in protein-protein interactions important for lysis regulation [53, 54]. Despite their similar membrane architecture, gp11 and LysF1a exhibit limited primary sequence conservation, sharing less than 15% amino acid identity and less than 28% similarity, with no strongly conserved subdomains identified by pairwise alignment (Figure 1B). Consistent with this divergence, the proteins belong to different phams within the Actinobacteriophage Database, however, they share similar features. Like *lysF1a*, gene *11* lacks the dual translational start motif characteristic of type II pinholins such as phage 21 S^21^68 [6, 13–15]. In addition, the N-terminal cytoplasmic domain of gp11 contains three charged residues together with several uncharged polar residues. This feature closely resembles LysF1a and suggest that the TMD1 of gp11 is unlikely to function like the TMD1 of the classical type II pinholin [6, 13–15]. Based on its predicted topology, genomic context, and our previously established nomenclature for actinobacteriophage lysis regulators [17], we designate gp11 as LysA2a.

Gene *30* encodes an 88-amino-acid protein with a single N-terminal transmembrane domain and a predicted N-out/C-in membrane topology, matching the 1TMD lysis regulator LysF1b (Figure 2A). Both proteins have C-terminal domains with > 50% charged KRHDE residues. AlphaFold3 predicts that gp30 adopts an elongated α-helical structure that closely resembles the predicted structure of LysF1b (Figure 2B). Consistent with the positive-inside rule [55], multiple positively charged residues are located immediately downstream of the transmembrane domain, supporting the predicted N-out/C-in orientation of the various TMD prediction programs (Figure 2BCD). Although gp30 shares only 25% overall amino acid identity with LysF1b, pairwise sequence alignment revealed a conserved 59-amino-acid region within the C-terminal cytoplasmic domain exhibiting 42% identity and 60% similarity, including 16 conserved charged residues (Figure 3D). The conservation of membrane topology, predicted secondary structure, and a highly conserved charged cytoplasmic domain strongly suggests that gp30 is a functional homolog of LysF1b even though this gene is not adjacent to another *lys* regulator or to the various *lysin* genes in D29. We therefore hypothesize that gp30 functions as a lysis regulator in D29 and designate this protein LysA2b.

**Figure 3.**
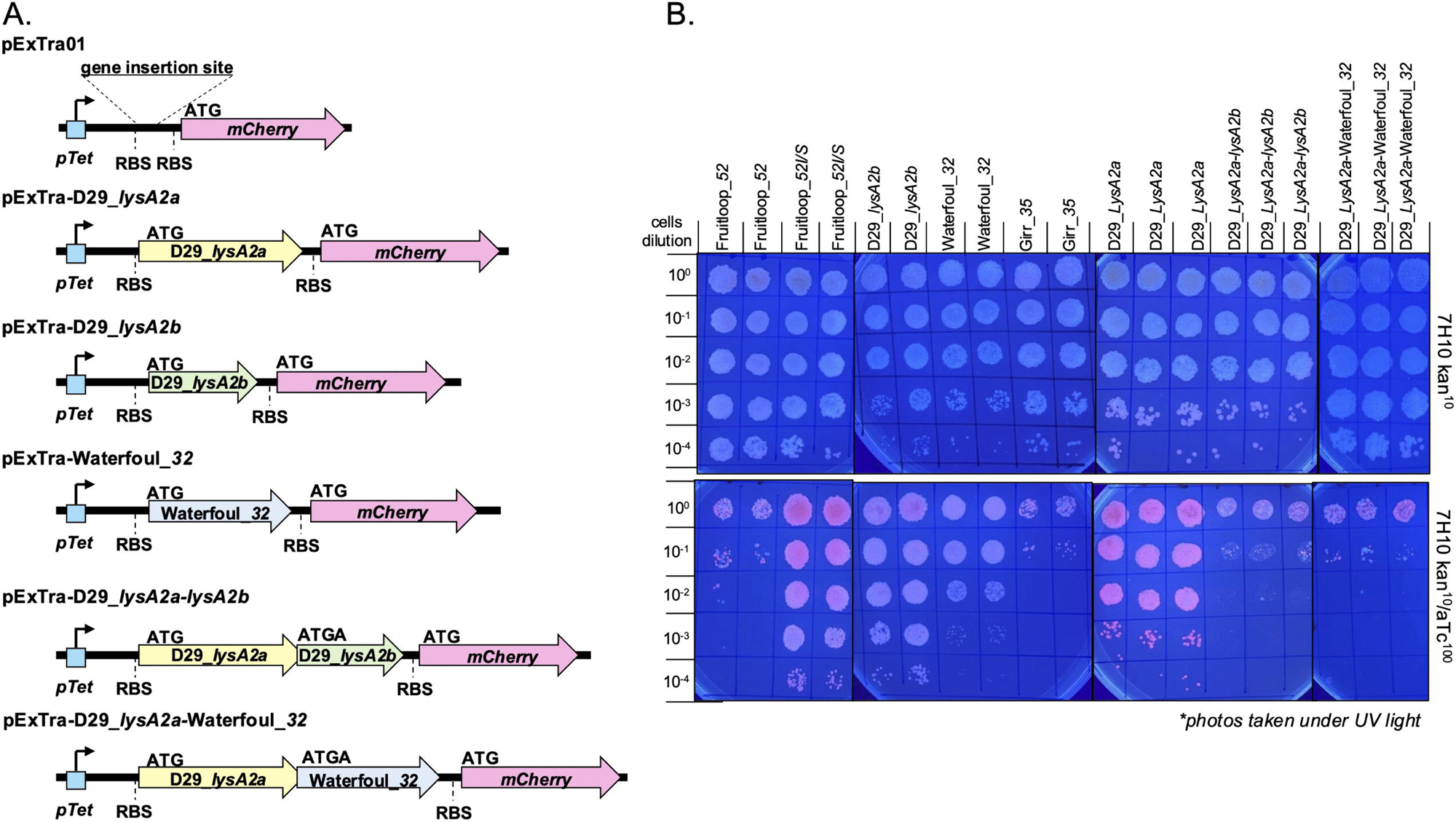
Cytotoxicity assay of Lys proteins. **A.** Schematic showing gene context of Lys genes inserted into pExtra01 between the pTet promoter and *mcherry* gene. RBS is the ribosomal binding site. ATG indicates start codon. ATGA is the 4bp overlap of the tandem genes indicating the TGA stop codon of the 1^st^ gene and ATG start codon of the 2^nd^ gene. **B.** In each assay, an individual colony of *M. smegmatis* transformed with the specified pExTra plasmid was resuspended in 7H9, serially diluted, and spotted on 7H10 Kan agar containing 0 or 100 ng/mL aTc and incubated at 37°C as detailed in Methods. Plates were inverted and photographed under UV light to visualize the mCherry expression. Fruitloop_*52* serves as the positive cytotoxicity control and the Fruitloop_*52I/S* serves as the negative control [31, 32].

### Exogenous expression of LysA2a or LysA2b in *M. smegmatis* is not cytotoxic, while co-expression of both LysA2a and LysA2b is highly cytotoxic

Previous studies demonstrated that exogenous expression of the 1TMD lysis regulator LysF1b is highly cytotoxic to *M. smegmatis*, whereas exogenous expression of the 2TMD protein LysF1a has little effect on bacterial viability [17, 22, 36]. To determine whether the D29 proteins LysA2a and LysA2b exhibit similar activities, each gene was cloned individually into the pExTra01 expression vector and evaluated for cytotoxicity as described in Methods. The resulting expression constructs are shown in Figure 3A.

Like LysF1a, induction of LysA2a did not produce a toxic phenotype (Figure 3B). Surprisingly, induction of LysA2b also had little effect on bacterial growth (Figure 3B). This result contrasts with the strong toxicity observed following expression of LysF1b and the Fruitloop_52 positive-control protein, both of which produced the expected growth inhibition phenotype (Figure 3B). Detection of mCherry fluorescence in the induced cultures confirms successful transcription and translation from the pExTra constructs, indicating that the absence of toxicity is likely not due to a lack of protein expression.

The lack of toxicity associated with LysA2b expression is unexpected as several previously characterized 1TMD LysF1b homologs, including Adephagia gp33 [56], Amelie gp29 [57], Hammy gp32 [58], and LeBron gp29 [59], exhibits significant cytotoxicity when expressed in *M. smegmatis* while Waterfoul gp32 has a moderate cytotoxicity (Figure 3B [32]). LysA2b is one of the smaller 1TMD homologs at 88-amino acids (Figure 2), but its apparent inactivity suggests that its function might depend on the presence of an additional phage-encoded factor. To test this, the lysA2a and lysA2b open reading frames were cloned in tandem into pExTra01 using a four-base-pair ATGA stop/start overlap as shown in Figure 4A. This translational arrangement closely resembles the organization of transmembrane *lysis regulator* genes typically found adjacent to each other in canonical actinobacteriophage lysis cassettes [17, 31].

**Figure 4.**
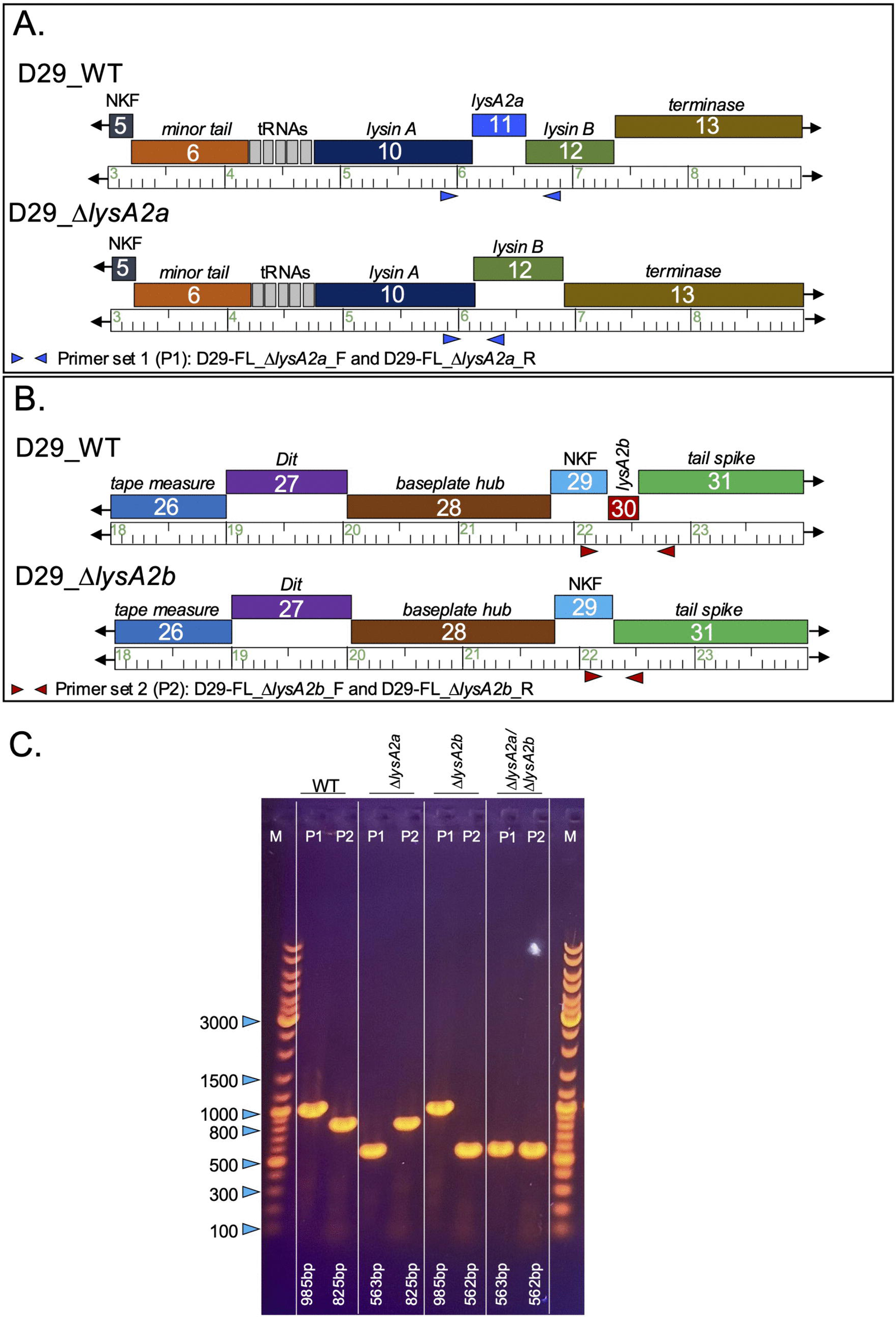
Deletion of D29 *lysA2a* and *lysA2b*. **A.** Genomic structure and PCR primer set locations for deletion of gene *11* (*lysA2a*). Primer set 1 (P1) uses primers D29-FL_*lysA2a* (F and R) and generates a 985bp fragment in D29_WT and 563bp with *lysA2a* deletion. Gene functions and location of tRNA genes are indicated. **B.** Genomic structure and PCR primer set locations for deletion of gene *30* (*lysA2b*). Primer set 2 (P2) uses primers D29-FL_*lysA2b* (F and R) and generates an 825bp fragment in D29_WT and 562bp with *lysA2b* deletion. Gene functions are indicated. NKF encodes a hypothetical protein with no known function. **C.** PCR verification of high titer lysates of D29_WT, D29_Δ*lysA2a*, D29_Δ*lysF1b* and D29_Δ*lysA2a*/Δ*lysF1b* using the indicated primer sets. PCR band sizes are indicated and show all phage have the expected deletions.

In contrast to the individual expression constructs, induction of the bicistronic lysA2a-lysA2b construct resulted in a pronounced cytotoxic phenotype, with substantial growth inhibition observed following aTc treatment (Figure 3B). To determine whether this effect was specific to LysA2b or reflected a more general interaction between LysA2a and 1TMD lysis regulators, LysA2a was co-expressed with Waterfoul gp32, a moderately toxic 1TMD LysF1b homolog that has been shown to functionally complement *lysF1b* deletions in both Girr and NormanBulbieJr [17]. Like the D29 bicistronic construct, co-expression of LysA2a and Waterfoul gp32 produced a highly toxic phenotype.

These findings indicate that neither LysA2a nor LysA2b is sufficient to disrupt host viability when expressed alone but suggest that the two proteins act synergistically to produce cytotoxicity. Moreover, the ability of Waterfoul gp32 to substitute for LysA2b suggests that the observed phenotype reflects a conserved functional interaction between the 2TMD and 1TMD classes of actinobacteriophage lysis regulators. Collectively, these data support the hypothesis that LysA2a and LysA2b function together during D29-mediated lysis and are consistent with the cooperative lysis-regulatory mechanism previously proposed for the LysF1a/LysF1b system of F1 cluster phages [17].

### Deletion of *lysA2a and lysA2b* in D29 is non-lethal but results in reduced plaque size

To directly evaluate the contributions of *lysA2a* and *lysA2b* to D29 lysis, individual and double deletion mutants were constructed using the BRED technique as described in Methods [34, 35]. Primary recombinant plaques containing each targeted deletion were identified by PCR screening, and clonal populations of D29_Δ*lysA2a*, D29_Δ*lysA2b*, and D29_Δ*lysA2a*/Δ*lysA2b* isolated through secondary plaque purification. The genomic organization of each mutant and PCR verification of the deletions are shown in Figure 4. The deletion junctions were confirmed by Sanger sequencing, and whole-genome sequencing verified the absence of secondary mutations elsewhere in the genome.

To determine the effects of these mutations on phage growth, equivalent titers of D29_WT, D29_Δ*lysA2a*, D29_Δ*lysA2b*, and D29_Δ*lysA2a*/Δ*lysA2b* were plated on *M. smegmatis*, and plaque formation efficiencies and plaque dimensions were measured as described in Methods. Plaque data us summarized in Table 2. All mutant phages formed plaques with efficiencies comparable to wild-type D29, indicating that deletion of either or both genes does not impair phage viability or adsorption under standard assay conditions.

**Table 2.** Plaque sizes in D29_WT and *lysF1a* and *lysF1b* deletion mutants.

| PHAGE | Plaque size (mm) | % of WT (diam.) | Plaque (vol, mm<sup>3</sup>) | % of WT (plaque vol) |
| --- | --- | --- | --- | --- |
| D29_WT | 4.96 + /- 0.46 | 100 | 19.32 + /- 3.46 | 100 |
| D29_ΔlysF1a | 3.45 + /- 0.49* | 69.6* | 9.34 + /- 2.65* | 48.34* |
| D29_ΔlysF1b | 1.58 + /- 0.59* | 31.85* | 1.96 + /- 1.46* | 10.14* |
| D29_ΔlysF1a/ΔlysF1b | 1.25 + /- 0.37*# | 25.20*# | 1.23 + /- 0.73*# | 6.31*# |

*M. smegmatis* was infected with identical numbers of phage and plaque size determined after 36 hrs. of growth at 37°C. Plaque diameter was measured using ImageJ. Data is presented as average + /- standard deviation for a minimum of 50 individual plaques. \* = statistically significant from D29\_WT phage $p < 0.0001$ . # = statistically significant from ΔlysF1b phage $p = 0.030$ .

Although plaque numbers were unaffected, all three mutants produced significantly smaller plaques than wild-type D29 (Figure 5; Table 2), indicating that the deleted genes contribute to efficient phage propagation. Deletion of *lysA2a* produced a relatively modest phenotype, reducing plaque diameter by approximately 30% and plaque volume by 52%. This phenotype closely resembles that reported previously following deletion of D29 gene *11* [53], providing independent validation of the mutant phenotype. In contrast, deletion of *lysA2b* resulted in a much more severe defect, reducing plaque diameter by approximately 70% and plaque volume by nearly 90%. The Δ*lysA2a/lysA2b* double mutant exhibited the strongest phenotype, with reductions of approximately 75% in plaque diameter and 94% in plaque volume compared to the D29_WT phage.

**Figure 5.**
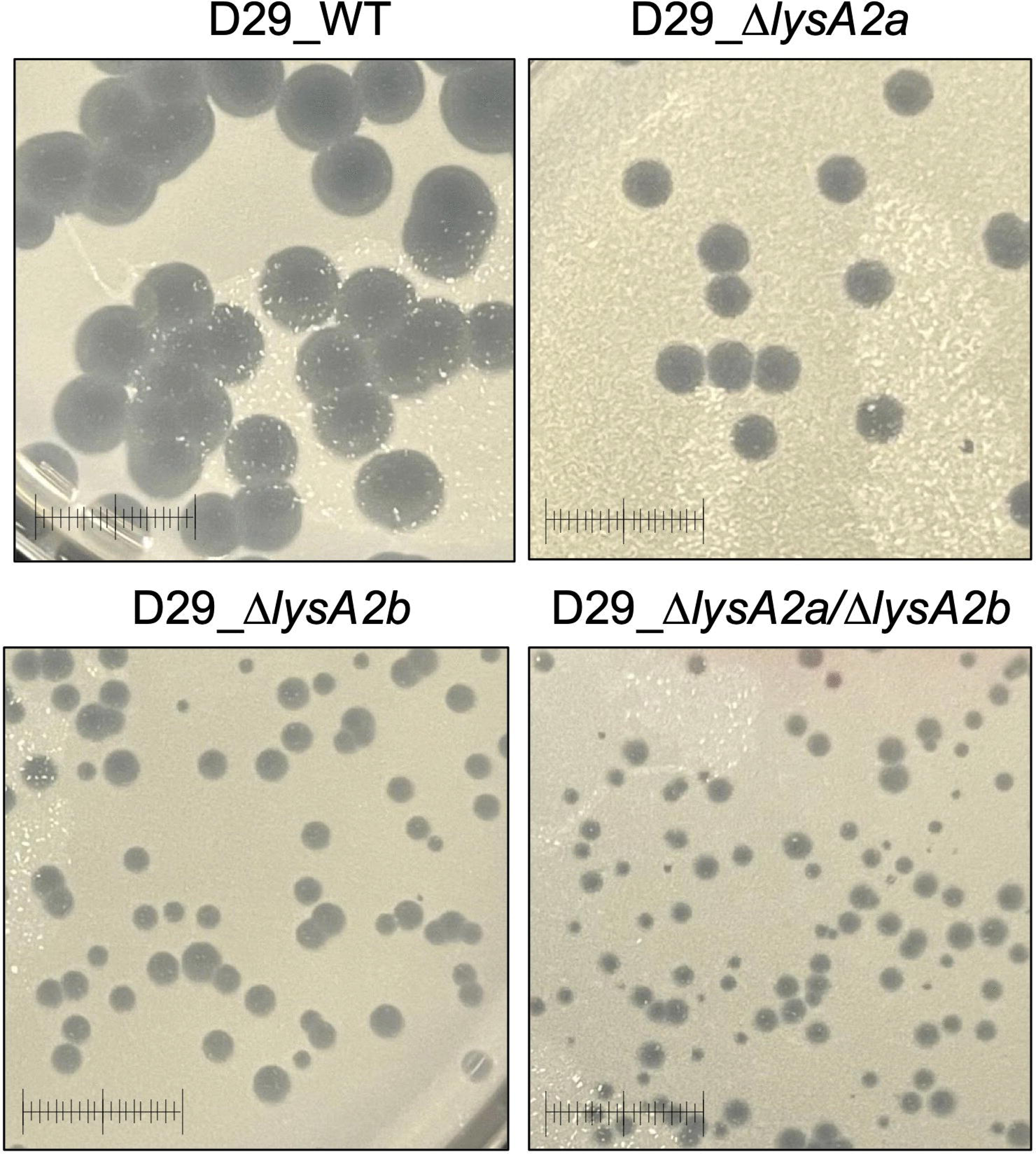
Plaque sizes in D29_WT, D29_ΔlysA2a, D29_ΔlysA2b and D29_ΔlysA2a/ΔlysF1b. *M. smegmatis* was infected with identical numbers of the indicated phage as detailed in Methods and plaque size determined after 36 hrs. of growth at 37°C. Scale = 1cm.

Although plaque diameter and volume of the double mutant were determined to be statistically different from the Δ*lysA2b* single mutant, the statistical difference was modest (*p = 0.03*). Thus, these results generally parallel those observed for the homologous LysF1a/LysF1b pair in F1 cluster phages, where deletion of the 1TMD regulator produces the dominant lysis phenotype [17]. The phenotype of the double mutant suggests that LysA2b is the primary determinant of the plaque-size defect, whereas LysA2a plays a supporting or regulatory role.

Together with the co-expression toxicity experiments, these findings strongly support the conclusion that LysA2a and LysA2b function cooperatively during D29 lysis, with LysA2b serving as the principal effector of the pair despite the non-canonical genomic organization of the D29 lysis genes.

### Deletion of *lysA2* and *lysA2b* genes from D29 result in distinct *M. smegmatis* liquid lysis phenotypes

To directly assess the impact of the *lysA2* gene deletions to host lysis by phage D29, liquid lysis assays were performed where the optical density (OD_600_) of infected log growth *M. smegmatis* cultures is measured over time as detailed in Methods. Figure 6A shows that cultures infected with D29_WT grow steadily until ∼85 minutes post infection. Beginning at 85-90 minutes there is a significant drop in the OD_600_ of the culture (triggering event), until it reaches a baseline of OD_600_ ∼0.05 by 200 minutes post infection. Cells infected with D29_Δ*lysA2a* also show a defined triggering event, but it is consistently delayed by ∼10 minutes compared to D29_WT. Figure 6B shows the results for D29_Δ*lysA2a* on an expanded time scale to better illustrate the delay in triggering. The reduction in OD_600_ after triggering minors that of the D29_WT and both reach a similar OD_600_ baseline. In sharp contrast, cells infected with D29_Δ*lysA2b* do not show a defined triggering event, but the OD_600_ of the culture begins to decline between 150-180 minutes after infection and ultimately reaches an OD_600_ baseline of <0.1 after 400 minutes (> 6 hrs) (Figure 6A). Similar results are observed for cells infected with D29_Δ*lysA2a*/Δ*lysA2b*, although the reduction in OD_600_ is more protracted reaching an OD_600_ baseline of <0.1 after ∼600 minutes (> 10 hrs) post infection. To assess that there were no surviving cells in the infected cultures, 5ul of each culture was removed at 120 minutes post infection, spotted onto 7H10 agar plates and incubated for 36hr at 37°C. Figure 6C shows that all cultures infected with phage have no growth compared to the uninfected control. This result indicates that all cells in the culture have been killed in the experiment and surviving cells do not account for the reduced lysis timelines.

**Figure 6.**
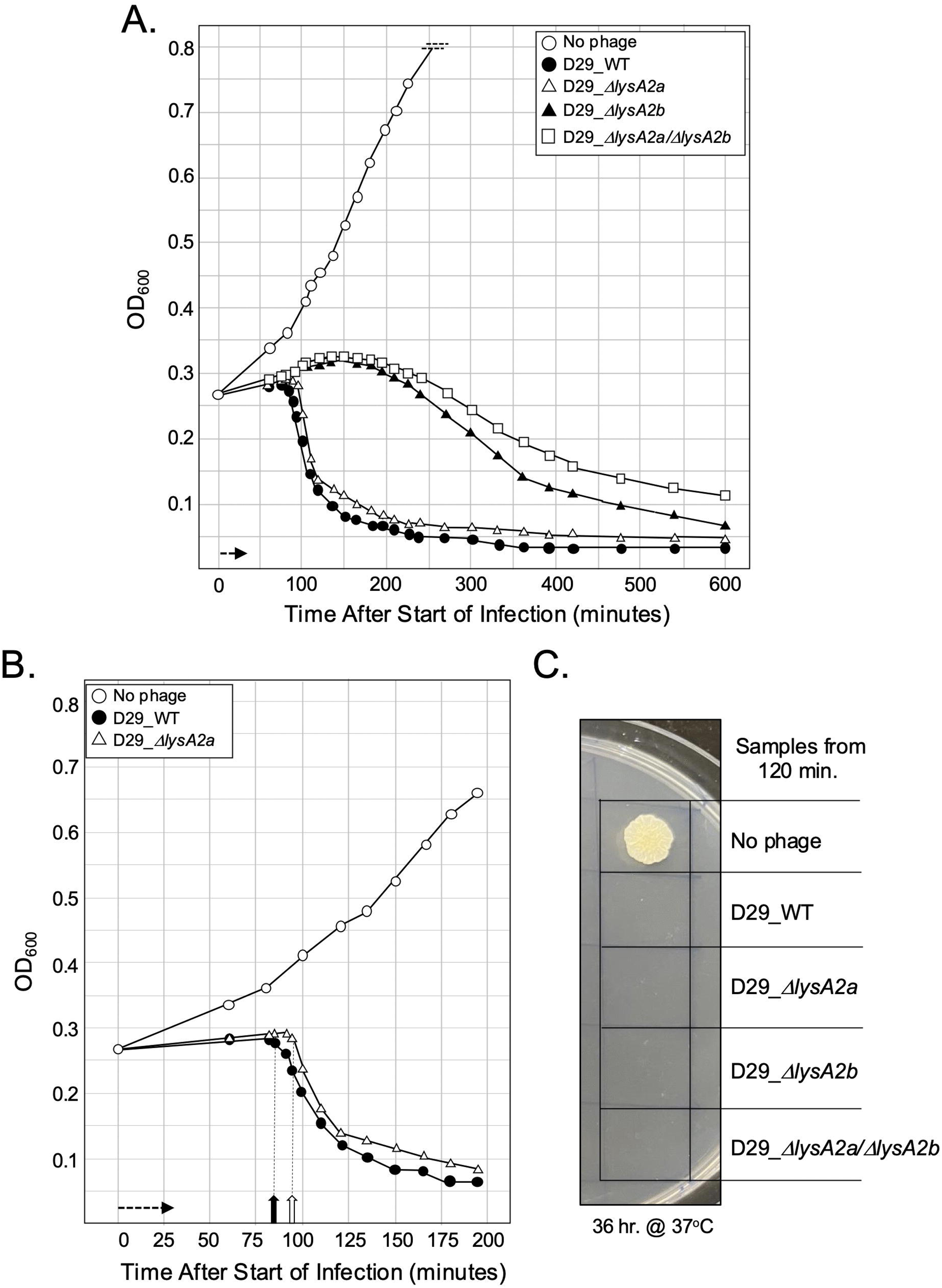
Liquid lysis assay of D29_WT and. Δ***lys* mutants.** At T_0_ a log growth culture of *M. smegmatis* at an OD_600_ of ∼0.25 was infected with the indicated phage at an MOI of 10 and incubated at 37°C for 30 minutes without shaking. Control cultures (no phage) did not receive phage. At T_30_ minutes the culture was constantly shaken a 225rpm, aliquots removed at the indicated times and bacterial growth evaluated at OD_600_. Horizonal arrow shows the 30-minute infection time. **A.** Representative results following infection with D29_WT, D29_Δ*lysA2a,* D29_Δ*lysA2b* and D29_Δ*lysA2a*/Δ*lysA2b*. **B**. Same data as presented in A but with extended x-axis to be show the difference in triggering of D29_WT and D29 Δ*lysA2a.* Solid vertical arrow shows the triggering of D29_WT and open vertical arrow ∼10-min delayed triggering time of D29 Δ*lysA2a.* **C.** Samples of each culture were removed at 120-minutes post infection and 5ul directly spotted onto 7H10 agar plates. Results show the level of bacterial growth after 36 hrs. at 37°C indicating no surviving cells in the infected cultures.

There are several important observations with this data set in comparison to the recent F1 cluster phage report [17]. First, the triggering time of 85 minutes for D29_WT is much earlier than the F1 phage Girr_WT (110 minutes) or NBJ_WT (130 minutes). This suggests that although both sets of phages utilize a similar set of *lys* proteins, there are clear phenotypic differences to how the proteins trigger the lysis pathway. Second, from the time of triggering, host lysis by D29_WT, Girr_WT and NBJ_WT D29_WT does not result in a precipitous 5–10-minute reduction in OD_600_ to baseline as observed for the lysis of *E. coli* [13–15]. Instead, host lysis occurs over an approximate 60-minute time frame indicating that the process is less efficient. Previous liquid lysis studies of D29_WT also show similar results [49, 51, 52]. Third, phages with deletions of the 2TMD *lysA2a* show a defined delay in triggering. However, following the triggering event, host lysis occurs without the 2TMD LysA2a protein along a similar 60-minute timeline as D29_WT phages. This result is identical to the F1 cluster data when the 2TMD *lysF1a* gene is deleted [17]. Finally, deletion of the 1TMD genes from either D29 or F1 cluster phages produces phages that do not initiate a defined triggering event but exhibit a slow, protracted reduction in OD_600_. For the mutant D29 phages, the reduction in OD_600_ is observable over 6 hrs from the time of infection while for the mutant F1 cluster phages, the decline in OD_600_ does not appear to begin until >8 hours post infection [17].

### Energy poisons cause early triggering by D29_Δ*lysA2b* and D29_Δ*lysA2a/*Δ*lysA2b* in liquid lysis assays

Phage-mediated host lysis is typically triggered by destruction of the host proton-motive force (PMF). Thus, one diagnostic assay that a protein is disrupting the PMF is the ability to cause early triggering of an infected culture by the addition of energy poisons (e.g., cyanide or the protonophore dinitrophenol) that destroy the PMF [14, 60]. For the F1 cluster phages, both wild type phages and those with deletion of the *lysF1a* gene could be triggered by KCN, defining the lysis regulator, LysF1b, as a “holin-like” protein [17]. In contrast, phages with a deletion of *lysF1b* or with deletion of both *lys* genes could not be triggered by KCN [17]. Thus, LysF1a did not exhibit “holin-like” function. To assess the D29 Lys proteins, *M. smegmatis* was infected with D29_WT, D29_Δ*lysA2b,* D29_Δ*lysA2a* and D29_Δ*lysA2a*/Δ*lysA2b* and the cultures treated with KCN as described in Methods. Figure 7 shows that KCN treatment of D29_WT at 70 minutes results in early triggering by 75 minutes. Like the F1 phages, KCN also triggered the D29_Δ*lysA2a* infected cells and the results are shown in S6 Figure 3 in the S2 File. However, in sharp contrast to the F1 phages with *lysF1b* deletions, KCN treatment resulted in a defined triggering event of cells infected with D29_Δ*lysF1b* (Figure 7). Similarly, KCN treatment of cells infected with D29_Δ*lysA2a*/Δ*lysA2b* resulted in a defined triggering event although there was a slight delay in its onset compared to the cells infected with the D29_Δ*lysA1b*.

**Figure 7.**
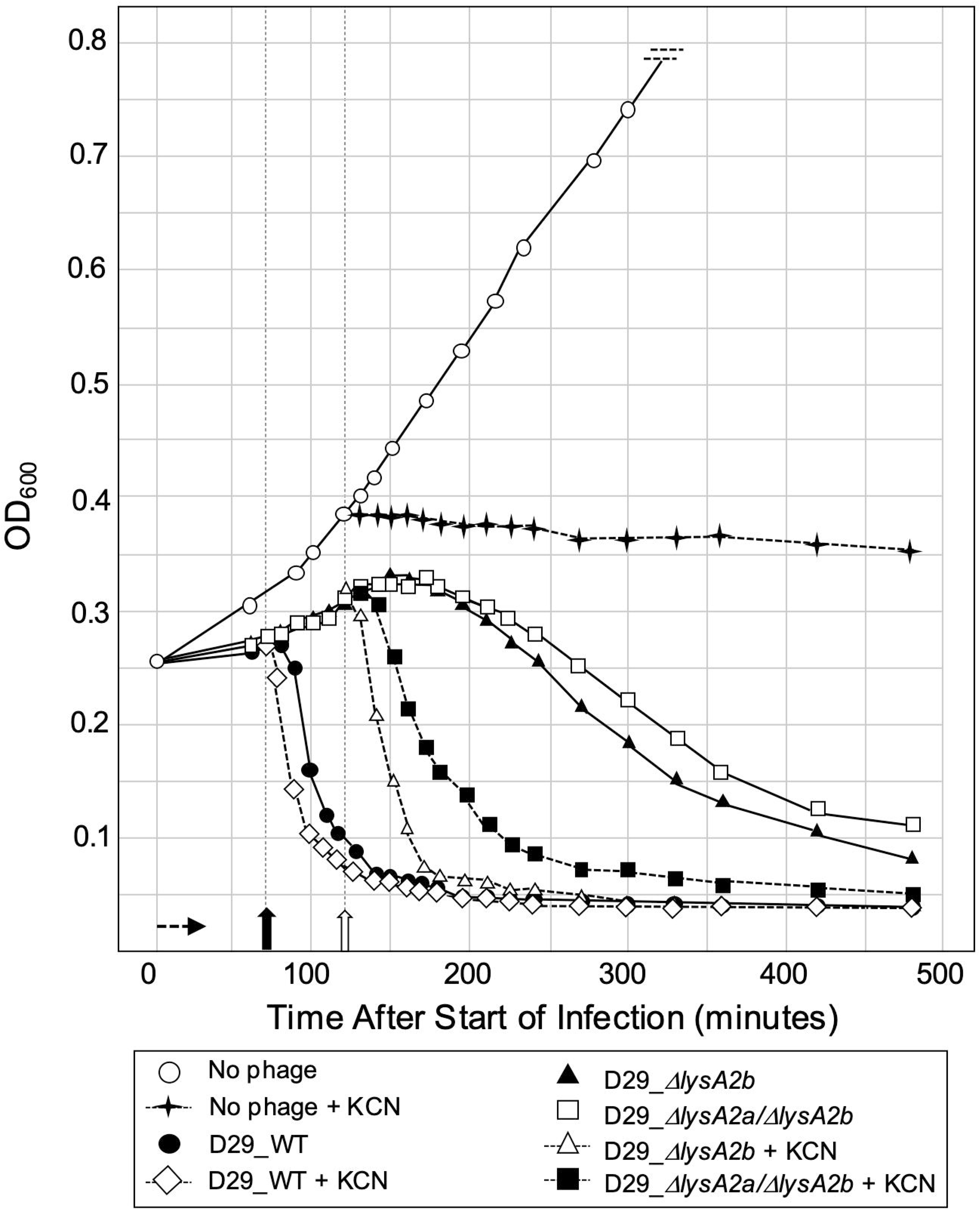
Energy poisons trigger early lysis in all D29 phages. At T_0_ a log growth culture of *M. smegmatis* at an OD_600_ of ∼0.25 was infected with the indicated phage at an MOI of 10 and incubated at 37°C for 30 minutes without shaking. Control cultures (no phage) did not receive phage. At T_30_ minutes the culture was constantly shaken a 225rpm, aliquots removed at the indicated times and bacterial growth evaluated at OD_600_. Horizonal arrow shows the 30-minute infection time. KCN (final concentration of 10mM) was added to a culture infected with D29_WT at 75 minutes (closed vertical arrow arrow). KCN (final concentration of 10mM) was added to a culture infected with D29_Δ*lysA2b*, D29_Δ*lysA2a*/Δ*lysA2b* and an uninfected culture at 120 minutes (open vertical arrow).

The F1 cluster phages Girr and NBJ are temperate and it was noted in our previous report that the liquid lysis assays of wild type phages did not show reductions in OD_600_ below 0.10 that was likely due to the presence of surviving lysogens after infection [17]. We also indicated a limitation of the work was that cultures of cells infected with phages deleted for *lysF1b* could not be evaluated past 480 minutes post infection due to the growth of the lysogens that would mask the OD_600_ reductions of infected cells. Since D29 is fully lytic and does not generate lysogens [19–21], it was of interest to assess if the phage lifecycle was impacting the KCN results of the F1 phages. Thus, the putative immunity repressor, gene *46,* was deleted from Girr_WT and from Girr_Δ*LysF1b* using BRED as described in Methods. Phages Girr_WT/Δ*46* and Girr_Δ*lysF1b*/Δ*46* were fully lytic, did not produce lysogens and showed OD_600_ reductions lower than the parent phages without the gene *46* deletions in liquid lysis assays [S7

Figure 4 in S2 File]. In addition, Girr_Δ*lysF1b*/Δ*46* shows a reduction in OD_600_ that begins ∼300-minutes after infection compared to Girr_Δ*lysF1b*. To directly compare the Girr and D29 phages and the response to KCN treatment, Girr_Δ*46*, Girr_Δ*lysF1b*/Δ*46,* D29_WT and D29_Δ*lysA2b* were evaluated in the liquid lysis assay. Figure 8A shows that D29_WT triggers lysis at 85 minutes and as expected, Girr_Δ*46* triggers later at ∼110 minutes, and both reach a similar OD_600_ baseline below 0.1. Cells infected with D29_Δ*lysA2b* show the expected protracted reduction in OD_600_ starting at 180 minutes, while cells infected with Girr_Δ*lysF1b*/Δ*46* show a much slower lysis profile, but have a defined decline in OD_600_ by ∼300 minutes that reaches an OD_600_ of 0.2 by 12 hours post infection (Figure 8A). Importantly, there are no surviving cells in any of the cultures as demonstrated by the spot test in Figure 8B. Figure 8A shows that KCN treatment at 150 minutes triggers D29_Δ*lysA2b*, but KCN does not trigger Girr_Δ*lysF1b*/Δ*46,* just as previously shown for Girr_Δ*lysF1b* [17].

**Figure 8.**
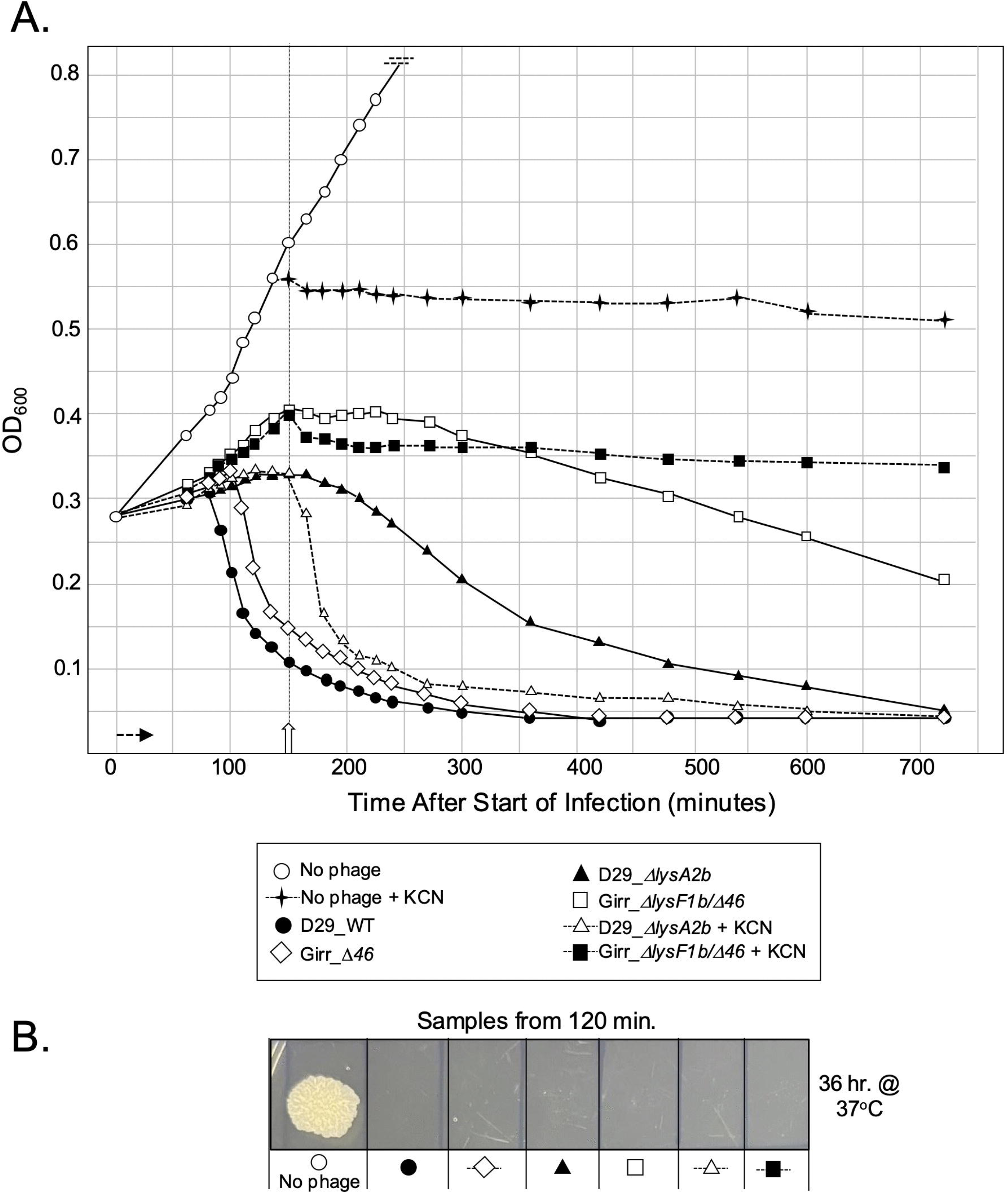
Comparison of liquid lysis profiles and KCN response of D29 and Girr phages with deletions of the immunity repressor (gene *46*). **A**. At T_0_ a log growth culture of *M. smegmatis* at an OD_600_ of ∼0.25 was infected with the indicated phage at an MOI of 10 and incubated at 37°C for 30 minutes without shaking. Control cultures (no phage) did not receive phage. At T_30_ minutes the culture was constantly shaken a 225rpm, aliquots removed at the indicated times and bacterial growth evaluated at OD_600_. Horizonal arrow shows the 30-minute infection time. KCN (final concentration of 10mM) was added to a culture infected with D29_Δ*lysA2b*, Girr_Δ*lysA2a*/Δ*46* and an uninfected culture at 150 minutes (open vertical arrow). **B.** Samples of each culture that did not receive KCN were removed at 120-minutes post infection and 5ul directly spotted onto 7H10 agar plates. Results show the level of bacterial growth after 36 hrs. at 37°C indicating no surviving cells in the infected cultures. .

Collectively, these experiments demonstrate that the D29 lysis pathway distal to triggering, differs fundamentally from that of the F1 cluster phages. Whereas deletion of *lysF1b* abolishes KCN-induced triggering in Girr and NBJ, KCN efficiently triggers lysis of D29 phages lacking *lysA2b* or both predicted lysis regulators. Elimination of lysogen formation in Girr by deletion of the immunity repressor did not alter this phenotype, demonstrating that the distinct KCN responses reflect intrinsic differences in the lysis mechanisms rather than differences in phage lifestyle. These findings indicate that D29 encodes an additional PMF-responsive component capable of promoting lysis independently of the LysA2 proteins.

### Genetic complementation of D29_**Δ***lysA2b* with LysF1b rescues the lysis defect phenotypes but changes the lysis triggering time

Based on the results in the previous section, it was pertinent to assess the ability of the Girr LysF1b protein to compliment the lysis defects of the D29_Δ*lysA2b* phage. Since exogenous expression of LysF1b is toxic (Figure 3) [31], BRED was utilized to insert the Girr *lysF1b* gene into D29_Δ*lysA2b.* The genomic structure, gene boundary sequences and PCR verification of the resulting D29_Δ*lysA2b*::Girr_*lysF1b* phage is shown in S8 Figure 5 in the S2 File. In the plaque assay, D29_Δ*lysA2b*::Girr_*lysF1b* rescues the small plaque phenotype observed in D29_Δ*lysA2b* and generates plaques that are 86% the size of D29_WT (Figure 9A). Liquid lysis assays show that D29_Δ*lysA2b*::Girr_*lysF1b* recovers the triggering defect of the D29_Δ*lysA2b* phage but effectively triggers lysis at approximately 110 minutes (Figure 9B).

**Figure 9.**
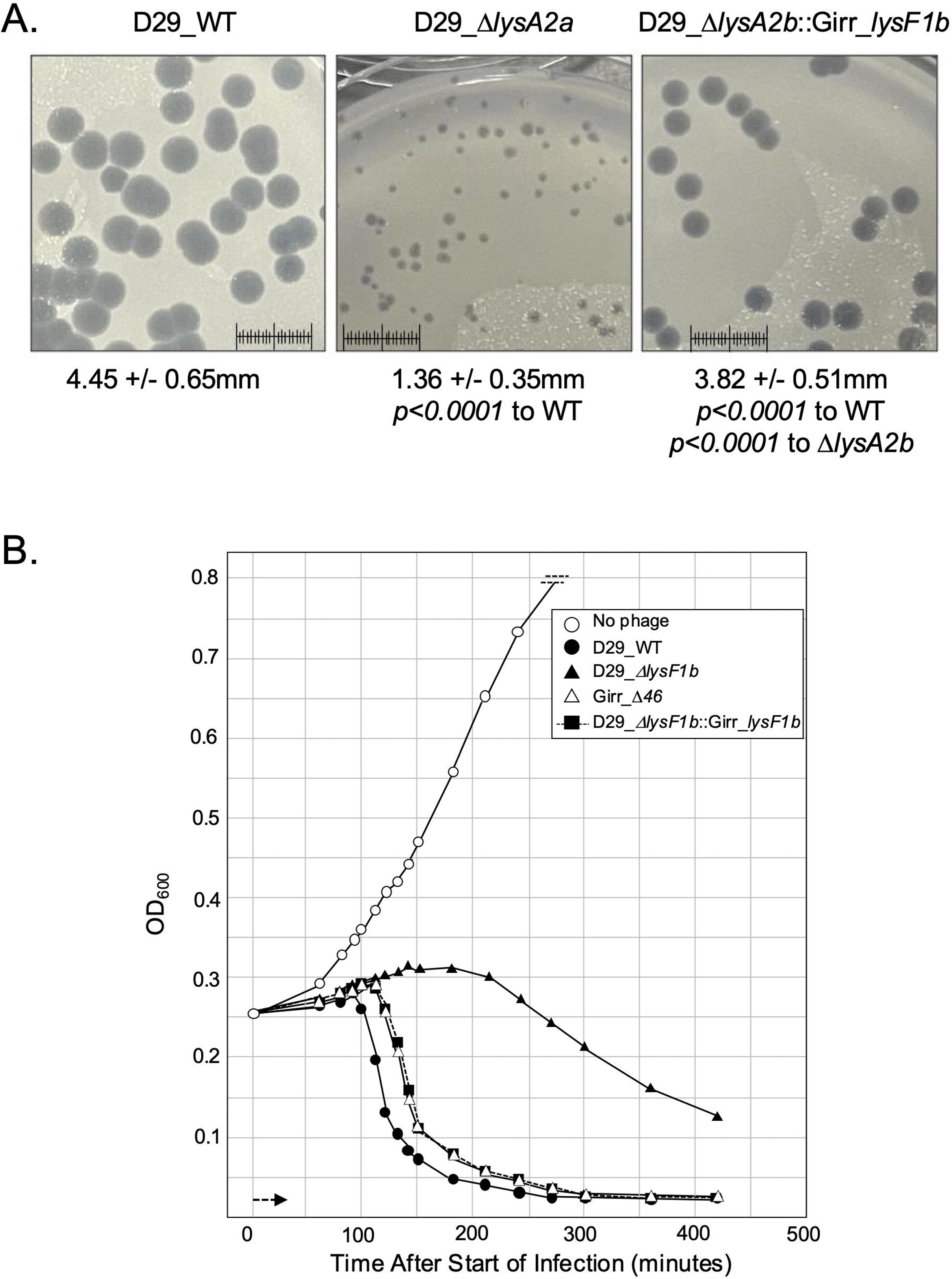
Plaque size and liquid lysis assay of D29 phage with *lysF1b* gene insertion. **A**. *M. smegmatis* was infected with identical numbers of the indicated phage as detailed in Methods and plaque size determined after 36 hrs. of growth at 37°C. Scale = 1cm. Plaque diameters and statistically significance is indicated. **B.** At T_0_ a log growth culture of *M. smegmatis* at an OD_600_ of ∼0.25 was infected with the indicated phage at an MOI of 10 and incubated at 37°C for 30 minutes without shaking. Control cultures (no phage) did not receive phage. At T_30_ minutes the culture was constantly shaken a 225rpm, aliquots removed at the indicated times and bacterial growth evaluated at OD_600_. Horizonal arrow shows the 30-minute infection time.

Strikingly, the triggering time, slope and overall lysis timeline of the D29_Δ*lysA2b*::Girr_*lysF1b* phage mirrors that of the Girr_Δ*46* (Figure 9B). Taken together, this data strongly supports the hypothesis that LysF1b and LysA2b are homologs and the expression of these 1TMD proteins determine the triggering time following phage infection.

### D29 deleted for *lysF1b* have greatly reduced burst timing and burst size

To assess phage release from the various D29 phages, one-step growth experiments were performed as described previously using infective centers to determine total phage production and burst size [17]. Figure 10 shows the one-step growth curves for D29_WT, D29_Δ*lysA2a* and D29_Δ*lysA2a*/Δ*lysA2b*, with Girr_WT included for comparison. D29_WT exhibited a lag of approximately 120 min., followed by gradual phage release over the next 90 min. before reaching a plateau at approximately 210 min. This timeline is consistent with the liquid lysis experiments, in which D29_WT showed a lysis-triggering event at approximately 85 min. post-infection. The gradual release of phage over approximately 90 min. is also consistent with the approximately 60-min. period of OD_600_ decline following the triggering event in liquid lysis assays.

**Figure 10.**
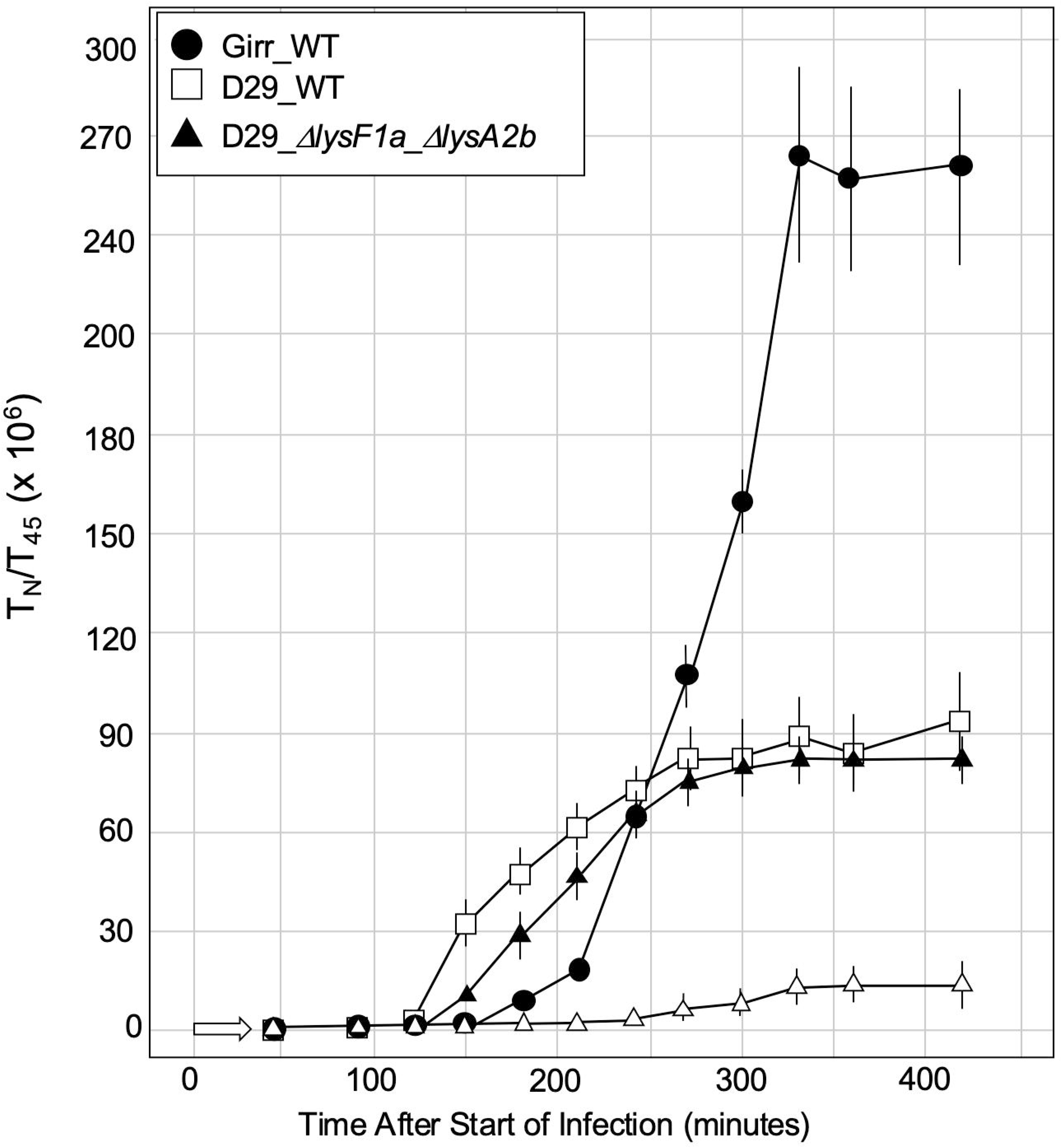
One-step growth curves of *M. smegmatis* infected with D29_WT, D29_Δ*lysA2b* and D29_Δ*lysA2a*/Δ*lysA2b*. **A**. One-step growth curves using the indicated phages were completed as described in Methods. Each time point represents the average and standard deviation from 3 independent experiments. The average bust size +/-standard deviation from this data set is presented in Table 3.

Table 3 summarizes the results, and the burst size of D29_WT was 105 ± 35 PFU/cell across three independent experiments, approximately one-third of the burst size measured for Girr_WT (310 PFU/cell). The substantially lower burst size of D29_WT may, at least in part, reflect its approximately 25-min earlier lysis-triggering time compared with Girr_WT, providing less time for phage genome replication and assembly before lysis. The D29_WT burst size is also consistent with recent studies reporting burst sizes of approximately 90 PFU/cell for wild type D29 [49, 51].

**Table 3.**
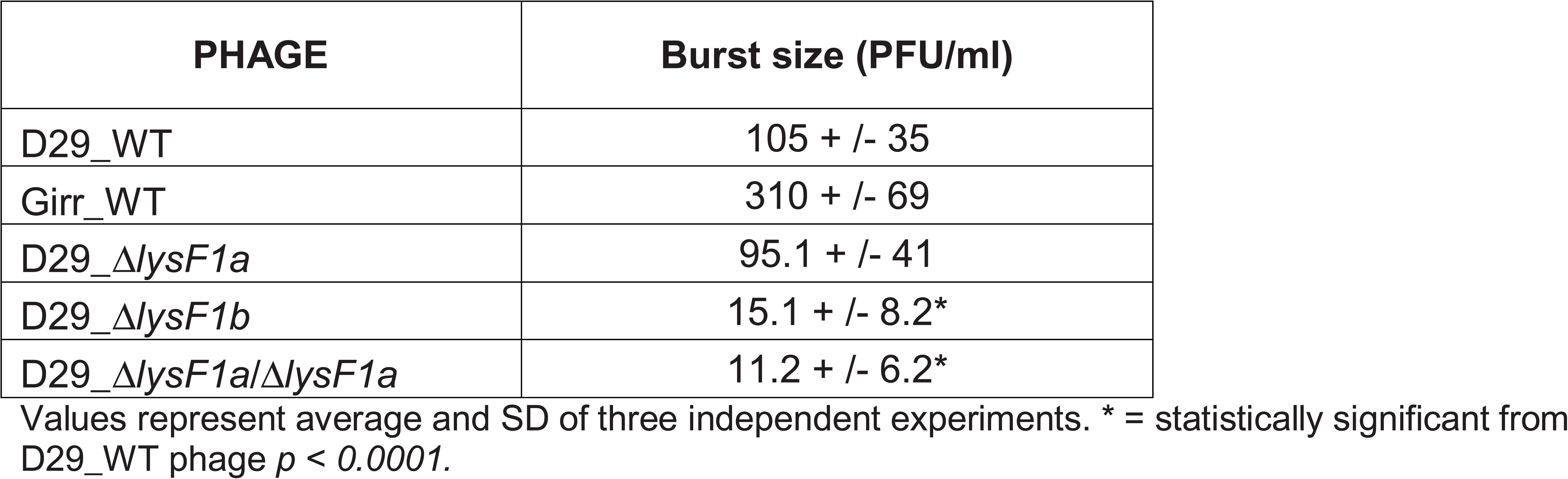
Burst size of Girr_WT, D29_WT and *lysA2* mutants.

Table 3 summarizes the results, and the burst size of D29_WT was 105 ± 35 PFU/cell across three independent experiments, approximately one-third of the burst size measured for Girr_WT (310 PFU/cell). The substantially lower burst size of D29_WT may, at least in part, reflect its approximately 25-min earlier lysis-triggering time compared with Girr_WT, providing less time for phage genome replication and assembly before lysis. The D29_WT burst size is also consistent with recent studies reporting burst sizes of approximately 90 PFU/cell for wild type D29 [49, 51].

Although deletion of *lysA2a* results in an approximately 30% reduction in plaque size, D29_Δ*lysA2a* efficiently killed *M. smegmatis* in liquid lysis assays (Figure 6). Consistent with this phenotype, D29_Δ*lysA2a* exhibited a modest delay in phage release and a slightly reduced, but not statistically significant, burst size of 95.1 PFU/cell compared with D29_WT (Figure 10). Thus, deletion of *lysA2a* has relatively little effect on the overall production of phage progeny possibly due to the 10-minute delay in triggering and production of more phage than D29_WT.

In striking contrast, D29_Δ*lysA2a*/Δ*lysA2b* exhibited a lag of more than 240 min before detectable phage release and produced a minimal burst of only 11.2 ± 6.2 PFU/cell across three independent experiments. This phenotype is similar to the burst size of 15.2 PFU/cell observed for D29_Δ*lysA2b* and is consistent with the greatly reduced burst sizes previously observed for F1 cluster phages lacking *lysF1b* [17]. Together, these results are consistent with the plaque-size and liquid-lysis phenotypes and indicate that although deletion of *lysA2b* does not completely prevent loss of *M. smegmatis* membrane integrity, it severely impairs the efficient release of progeny phage. Similar burst size phenotypes are observed in phages with deleted or impaired Lysin enzymes that would have reduced cell wall degradation [51, 52, 61]. Additionally, the near-wild-type burst size observed following infection with D29_Δ*lysA2a* further supports a primary role for LysA2b in regulating productive lysis and phage release as in F1 cluster phages. Thus, while LysA2a contributes to efficient lysis, LysA2b appears to be the principal lysis regulator required for efficient release of D29 progeny.

### Lysis recovery mutants of D29**Δ***lysA2b* restore wild-type plaque morphology through mutations in *lysA2a*

To identify genetic suppressors capable of restoring efficient lysis in the absence of LysA2b, spontaneous lysis recovery mutants (LRMs) were isolated by repeatedly propagating D29_Δ*lysA2b* and screening for plaques that had regained a wild-type morphology. Like our previous observations with F1 cluster phages [17], large-plaque variants were readily recovered from cultures infected with D29_Δ*lysA2b*. Phage isolated from individual large plaques retained the restored plaque phenotype following serial dilution and replating, demonstrating that the phenotype was genetically stable.

A total of 15 independent LRMs representing nine distinct mutations were identified. Remarkably, every suppressor mutation mapped within the *lysA2a* open reading frame despite the parental defect being deletion of *lysA2b* (Table 4). The positions of the recovered mutations are summarized in the Figure 11A schematic, and representative plaque morphologies are shown in Figure 11B. PCR analysis confirmed that all suppressor phages retained the original *lysA2b* deletion, while whole-genome sequencing verified the absence of additional mutations elsewhere in the genome.

**Figure 11.**
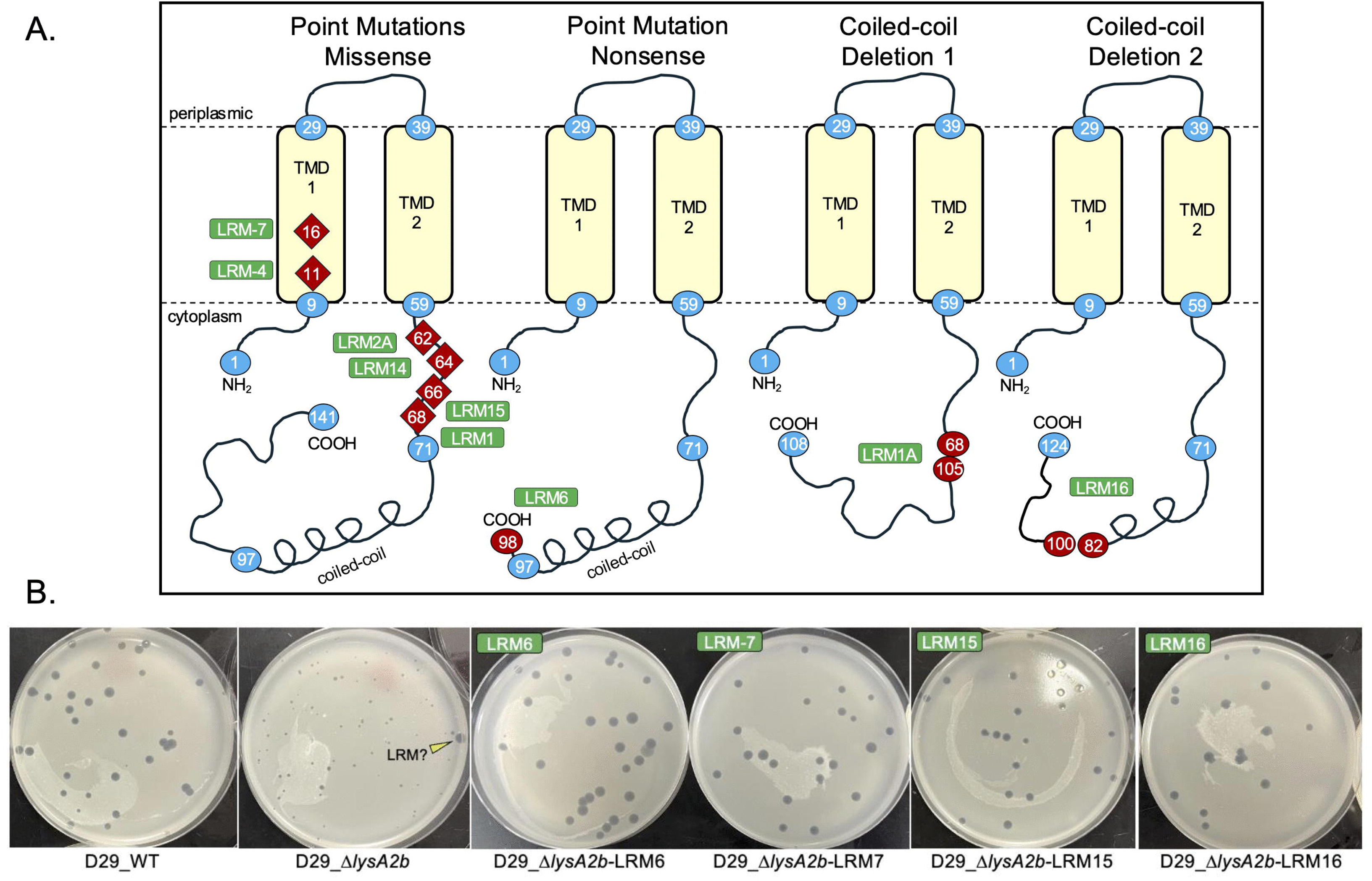
Location of mutations and plaque sizes of selected D29_Δ*lysA2b*-LRM mutants. **A**. The location of all D29_Δ*lysA2b*-LRM amino acid mutants in the LysA2a protein. **B.** *M. smegmatis* was infected with the indicated phages and plaque assays performed as described in Methods. Plates were incubated at 37°C for 36 hrs. A large plaque (LRM?) in the D29_Δ*lysA2b* background is indicated with the yellow arrowhead.

The recovered mutations cluster within functionally important regions of LysA2a. Two independent suppressors contained substitutions within the first transmembrane domain (Y11N and V16A). Similar transmembrane mutations were previously identified among suppressors isolated from F1 cluster phages, and none of the phages had any mutations within the second transmembrane domain [17]. Four additional D29 suppressors encoded substitutions immediately downstream of the second transmembrane domain (Q62L, R64S, D66G, and T68A), corresponding to the same general region in which multiple suppressor mutations were identified in the F1 phages [17]. In each case, the mutation replaced the native residue with an amino acid of markedly different chemical character, suggesting disruption of a conserved regulatory function rather than conservative sequence drift.

Unlike the approximately LysF1a proteins that only have a C-terminal cytoplasmic domain of 18-amino acids, D29 LysA2a contains an 80-amino acid C-terminal cytoplasmic domain that includes a predicted 27aa oiled-coil region (Figure 1). Consistent with this structural difference, additional suppressors unique to D29 disrupted this extended C-terminal domain. LRM1A and LRM16 contain deletion mutations that remove substantial portions of the predicted coiled-coil, whereas LRM6 introduces a premature stop codon at residue 99, truncating the distal 42 amino acids of the protein.

The genetic architecture of these suppressors provides important insight into the function of LysA2a. The recovery of wild-type plaque morphology through mutations exclusively within *lysA2a*, despite the continued absence of *lysA2b*, demonstrates that alteration of LysA2a alone is sufficient to bypass the requirement for the 1TMD regulator. These findings support the cytotoxicity results in Figure 3 and suggest that the wild-type LysA2a protein exists in a regulated, relatively inactive state. In the absence of LysA2b, mutations that alter the transmembrane domain, the conserved juxtamembrane cytoplasmic region, or the extended C-terminal domain appear to relieve this constraint, thereby restoring efficient lysis. Together with the analogous suppressor mutations previously identified in F1 cluster phages [17], these results support a conserved model in which the 2TMD protein is maintained in an inactive state until activated by its cognate 1TMD lysis regulator.

**Table 4.** Lysis Recovery Mutants (LRM) from D29_Δ*lysA2b* Phages.

| Phage | # Found | Region of Mutation to LysA2a | Amino Acid Number of LysA2a | Amino Acid Change to LysA2a | Characterization of the Amino Acid Mutation |
| --- | --- | --- | --- | --- | --- |
| D29_ΔlysA2b-LRM4A | 1 | TMD 1 | 11 | Y11N | nonpolar aromatic to polar |
| D29_ΔlysA2b-LRM7 | 2 | TMD 1 | 16 | V16A | nonpolar to nonpolar |
| D29_ΔlysA2b-LRM2A | 2 | C-terminal cytoplasmic | 62 | Q62L | polar to nonpolar |
| D29_ΔlysA2b-LRM14 | 2 | C-terminal cytoplasmic | 64 | R16S | polar charged to polar |
| D29_ΔlysA2b-LRM15 | 3 | C-terminal cytoplasmic | 66 | D66G | polar charged to nonpolar |
| D29_ΔlysA2b-LRM1 | 2 | C-terminal cytoplasmic | 68 | T68A | polar to nonpolar |
| D29_ΔlysA2b-LRM6 | 1 | C-terminal cytoplasmic | 100 | Q100stop | deletion of amino acids 100-141 |
| D29_ΔlysA2b-LRM1A | 1 | C-terminal coiled-coil | 69-104 | amino acid deletion | deletion of amino acids 71-103 |
| D29_ΔlysA2b-LRM16 | 1 | C-terminal coiled-coil | 83-99 | amino acid deletion | deletion of amino acids 83-99 |

### D29_Δ*lysA2b* lysis recovery mutants restore efficient lysis but exhibit distinct triggering times

Previous studies of the F1 cluster phages demonstrated that lysis recovery mutants (LRMs) isolated from Δ*lysF1b* phages trigger lysis substantially earlier than their wild-type parents, indicating that mutations in the 2TMD protein LysF1a bypass the requirement for the 1TMD LysF1b regulator while simultaneously accelerating lysis triggering [17]. These findings suggested that LysF1a is intrinsically capable of promoting lysis and is not simply an antiholin that antagonizes LysF1b activity.

To determine whether the D29 suppressor mutations behaved similarly, D29_Δ*lysA2b* LRMs were analyzed using liquid lysis assays as described in Methods. In contrast to the F1 phages, the D29 suppressors exhibited distinct lysis triggering phenotypes that correlated with the location of the mutation within LysA2a. The data for representative LRMs are shown in Figure 12. Two suppressors (LRM1A and LRM16) contain deletions within the extended C-terminal cytoplasmic domain that disrupt the predicted coiled-coil region (Figure 11A). As demonstrated for LRM16, these mutants consistently initiated lysis earlier than wild-type D29, closely resembling the phenotype previously observed for the F1 suppressor mutants.

**Figure 12.**
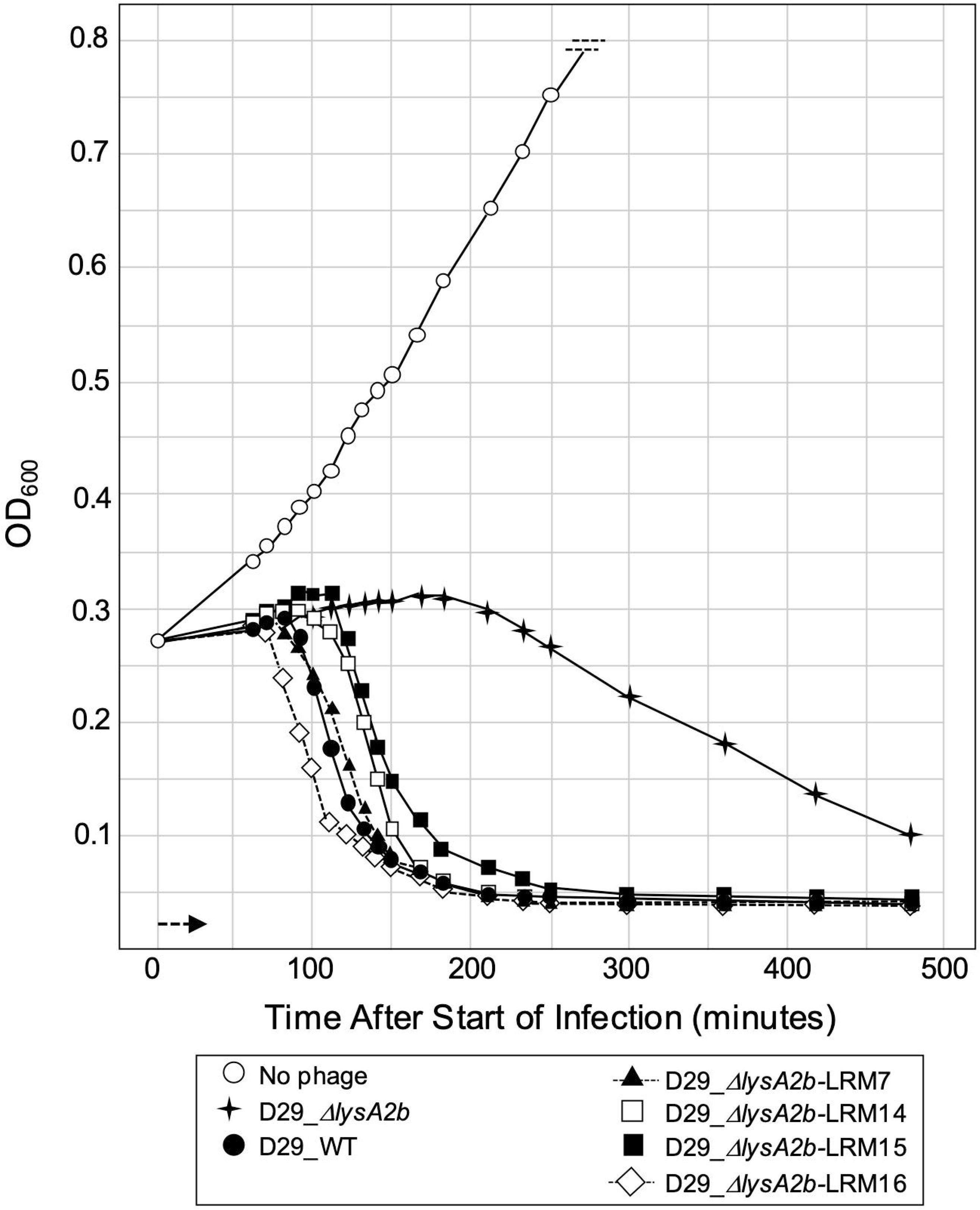
Liquid lysis assay of representative D29_Δ*lysA2b-*LRMs. At T_0_ a log growth culture of *M. smegmatis* at an OD_600_ of ∼0.25 was infected with the indicated phage at MOI 10 for 30 minutes at 37°C without shaking. Control cultures (no phage) did not receive phage. At T_30_ minutes the culture was constantly shaken a 225rpm and aliquots removed over time. *M. smegmatis* growth was evaluated at OD_600_.

A second class of suppressors (LRM1, LRM2A, LRM14, and LRM15) contains point mutations immediately downstream of the second transmembrane domain. In contrast to the coiled-coil mutants, these phages exhibited a reproducible delay in triggering, initiating lysis at approximately 100 min post-infection compared with approximately 85 min for wild-type D29 (Figure 12). Finally, suppressors containing substitutions within the first transmembrane domain (LRM4 and LRM7) displayed triggering kinetics that were nearly indistinguishable from wild-type D29. Despite their different triggering times, all D29 suppressors restored efficient plaque formation and eliminated the prolonged liquid lysis phenotype characteristic of D29_Δ*lysA2b*. Thus, restoration of efficient lysis is not strictly coupled to early triggering. Instead, these data demonstrate that distinct alterations within LysA2a can bypass the requirement for LysA2b while establishing different lysis timing programs.

These findings support a model in which the suppressor mutations promote a conformational state of LysA2a that is functionally active in the absence of LysA2b. However, unlike the F1 suppressors, which uniformly accelerate lysis, the D29 suppressors reveal that different structural regions of LysA2a contribute differently to regulation of lysis timing. Disruption of the extended C-terminal coiled-coil domain favors premature triggering, whereas mutations adjacent to the second transmembrane domain preserve LysA2a function but delay lysis initiation. The ability to genetically separate efficient lysis from trigger timing suggests that the extended cytoplasmic domain unique to D29 LysA2a provides an additional level of regulatory control that is absent from the shorter LysF1a proteins of the F1 cluster phages.

### Isolation of Lysis Recovery Mutants from D29_**Δ***lysA2a*/**Δ***lysA2b* identifies a novel lipoprotein that is dispensable but can modulate lysis when its localization is altered

Genetic suppressors capable of restoring efficient lysis in the absence of both LysF1a and LysF1b could not be identified in the F1 cluster phages. However, spontaneous lysis recovery mutants (LRMs) were isolated by repeatedly propagating D29_Δ*lysA2a/*Δ*lysA2b* and screening for plaques that had regained a wild-type morphology. Figure 13A shows the isolation of D29_Δ*lysA2a/*Δ*lysA2b-*LRM20 and the resulting large plaque phenotype that was genetically stable. In all, five different LRMs were isolated in D29_Δ*lysA2a/*Δ*lysA2b* phages and all showed the same single nucleotide mutation in codon 31 of gene *64* that changed cysteine 31 to a tyrosine. Gene *64* encodes a 137aa protein that is predicted to be a lipoprotein by Signal P 6.0, and it has the characteristic LVGC lipobox motif [62–64] and conserved cysteine residue necessary for lipidation (Figures 13B). HHpred analysis of gp64 returned many high probability hits (>98%) to putative lipoproteins with functions as sugar binding proteins and ABC transporters. Most relevant were hits to two *M. tuberculosis* proteins: PDB 8JA8_A: trehalose-binding lipoprotein LpqY [65] and PDB 4MFI_A: Sn-glycerol-3-phosphate ABC transporter substrate-binding protein UspB [66]. However, these proteins are 468aa and 438aa, respectively and gp64 aligned to 30% of these targets including the N-terminal secretory signal and conserved cysteine. AlphaFold3 models gp64 well with a pTM of 0.75 and shows that it has a single α-helix and a predominant β-sheet structure (Figure 13C). Importantly, the five identified D29_Δ*lysA2a/*Δ*lysA2b*-LRMs all have the C31Y mutation that is predicted to ablate the ability of gp64 to be tethered to the membrane and would cause it to be secreted into the periplasmic space.

**Figure 13.**
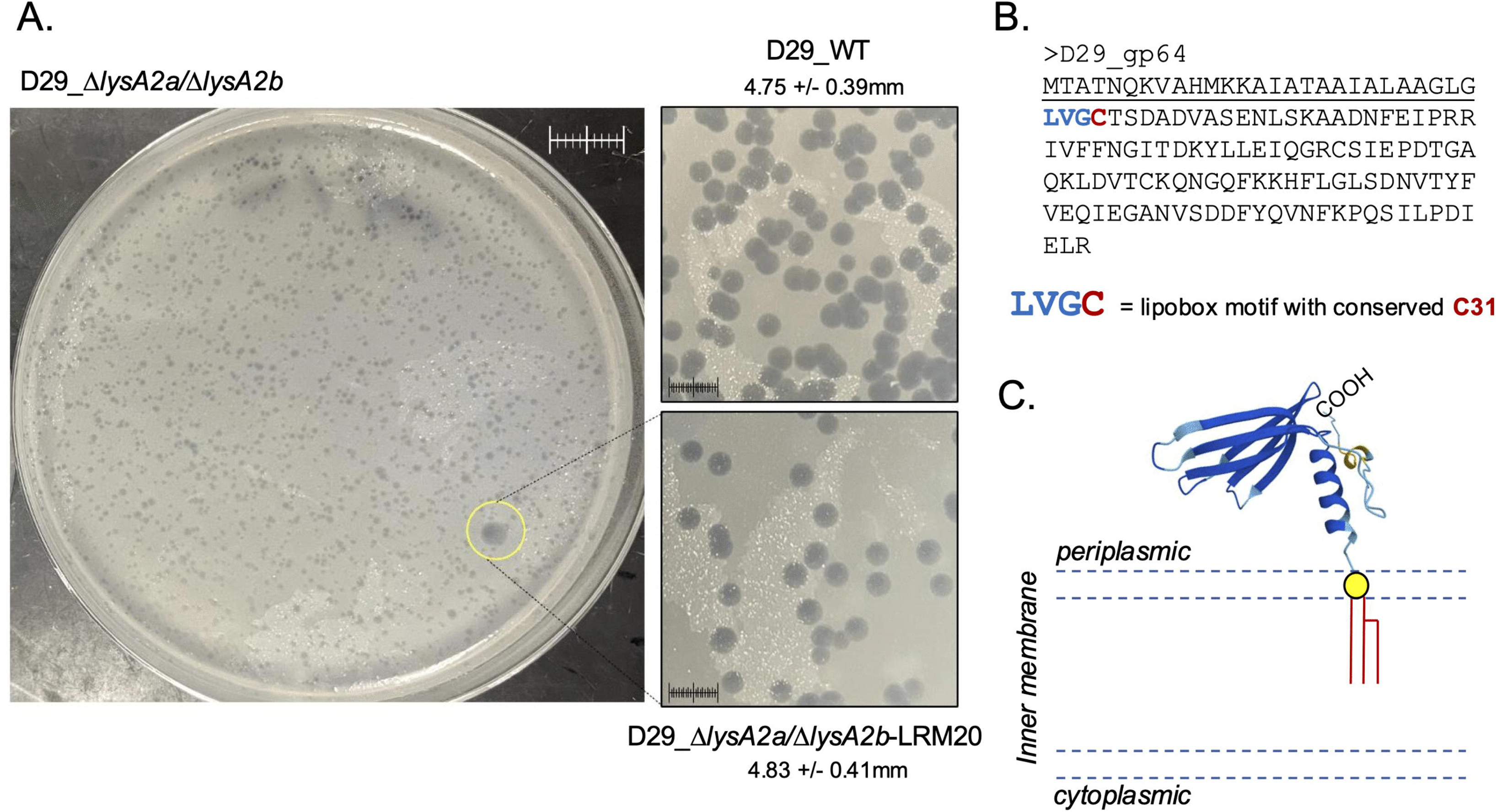
Identification of LRM mutants in D29_Δ*lysA2a/ΔlysA2b*. **A**. Plaque assay of D29_Δ*lysA2a/ΔlysA2b* showing a large plaque and isolation of D29_Δ*lysA2a/ΔlysA2b-*LRM20. *M. smegmatis* was infected with identical numbers of the indicated phage as detailed in Methods and plaque size determined after 36 hrs. of growth at 37°C. Average diameter +/-SD for a minimum of 20 plaques is shown. Scale = 1cm. **B**. Amino acid sequence of gp64 showing the cleaved signal sequence (underline), lipobox motif (LVGC) and conserved cysteine 31 residue (red). **C.** gp64 without the cleaved signal sequence was modeled using AlpahFold3 as described in Methods. The conserved cysteine 31 residue is shown as the yellow circle with the predicted lipidation and insertion in the outer leaflet of the inner membrane. pTM = 0.73. NH = N-terminus.

To begin to assess the function of gp64, it was of interest to determine if the expression of gp64 or the gp64C31Y protein was cytotoxic to *M. smegmatis*. PCR was utilized to amplify WT_*64* or *64*_C31Y and the resulting genes were ligated into pExTra01 and used for cytotoxicity analysis as described in Methods. Figure 14A shows that expression of either WT gp64 or gp64C31Y is fully toxic to *M. smegmatis*. Figure 14B also shows that D29_Δ*lysA2a/*Δ*lysA2b-*LRM20 triggers early lysis by 70 minutes post infection in the liquid lysis assay, approximately 15-20 minutes prior to D29_WT. The OD_600_ decline after triggering is rapid, reaching OD_600_ of 0.15 by 100 minutes, but then gently declines to below OD_600_ 0.1 by 420 minutes.

**Figure 14.**
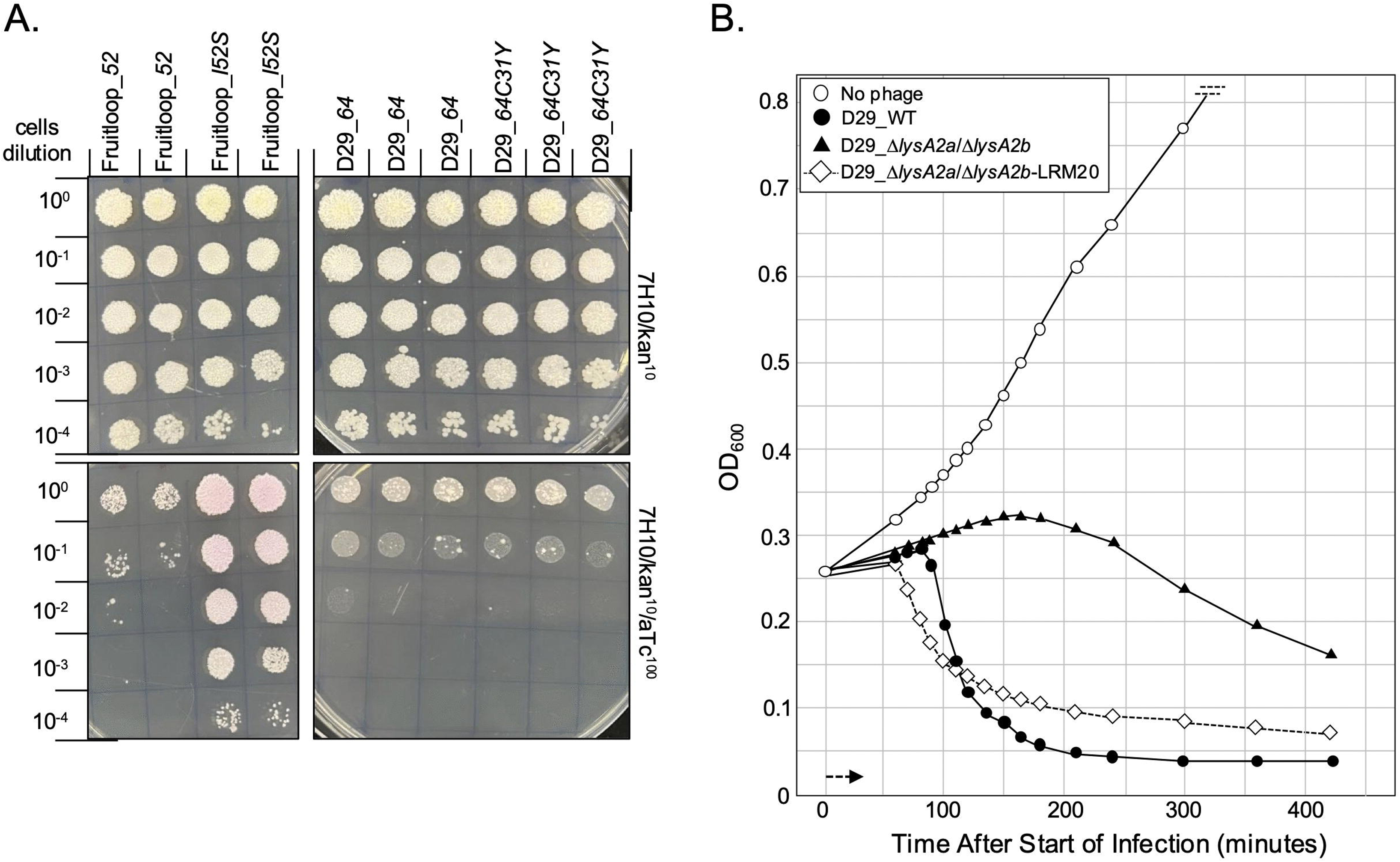
Cytotoxicity assay of gp64 and gp64C31Y and liquid lysis assay of D29_Δ*lysA2a/ΔlysA2b*-LRM20. **A**. The indicated genes were cloned into pExTra01 as described in Methods. The resulting plasmids were transfected into *M. smegmatis* and individual colonies with the specified pExTra plasmid was resuspended in 7H9, serially diluted, and spotted on 7H10 Kan agar containing 0 or 100 ng/mL aTc and incubated at 37°C as detailed in Methods. Fruitloop_*52* serves as the positive cytotoxicity control and the Fruitloop_*52I/S* serves as the negative control [31, 32]. **B**. At T_0_ a log growth culture of *M. smegmatis* at an OD_600_ of ∼0.25 was infected with the indicated phage at MOI 10 for 30 minutes at 37°C without shaking. Control cultures (no phage) did not receive phage. At T_30_ minutes the culture was constantly shaken a 225rpm and aliquots removed over time. *M. smegmatis* growth was evaluated at OD_600_.

Since the C31Y mutation in gp64 suppressed the plaque size and liquid lysis phenotypes of the D29_Δ*lysA2a/*Δ*lysA2b* phage, it was of interest to determine the impact of deleting gene *64* from the various D29 phages. BRED was utilized to delete gene *64* from D29_WT, D29Δ*lysA2b* and D29_Δ*lysA2a/*Δ*lysA2b.* The genomic context of gene *64* and PCR validation of the deletions is show in S9 Figure 6 in the S2 File. Deletion of *64* did not impact the viability of any of the phages and did not impact plaque size or morphology (S10 Figure 7 in S2 File). Figure 15A shows the liquid lysis curves for the phages with the Δ*64* mutations. D29_Δ*64* and D29_WT both trigger at ∼85 minutes and show identical timing of the OD_600_ decline. The curves for D29_Δ*lysA2b* and D29_Δ*lysA2b/*Δ*64* also show the same protracted timing and reach the same OD_600_ baseline by ∼400 minutes post infection. The curves for D29_Δ*lysA2a/*Δ*lysA2b* and D29_Δ*lysA2a/*Δ*lysA2b/Δ64* are also similar and show the expected protracted decline of OD_600_. Thus, deletion of gene *64* does not rescue or exacerbate any of the lysis phenotypes. Importantly, Figure 15B shows that both D29_Δ*lysA2a/*Δ*64* and D29_Δ*lysA2a/*Δ*lysA2b/Δ64* can be effectively triggered with KCN treatment. This is a key result as it indicates that gp64 is not required for KCN to trigger D29_Δ*lysA2a* and D29_Δ*lysA2a/*Δ*lysA2b* (Figures 7 and 8).

**Figure 15.**
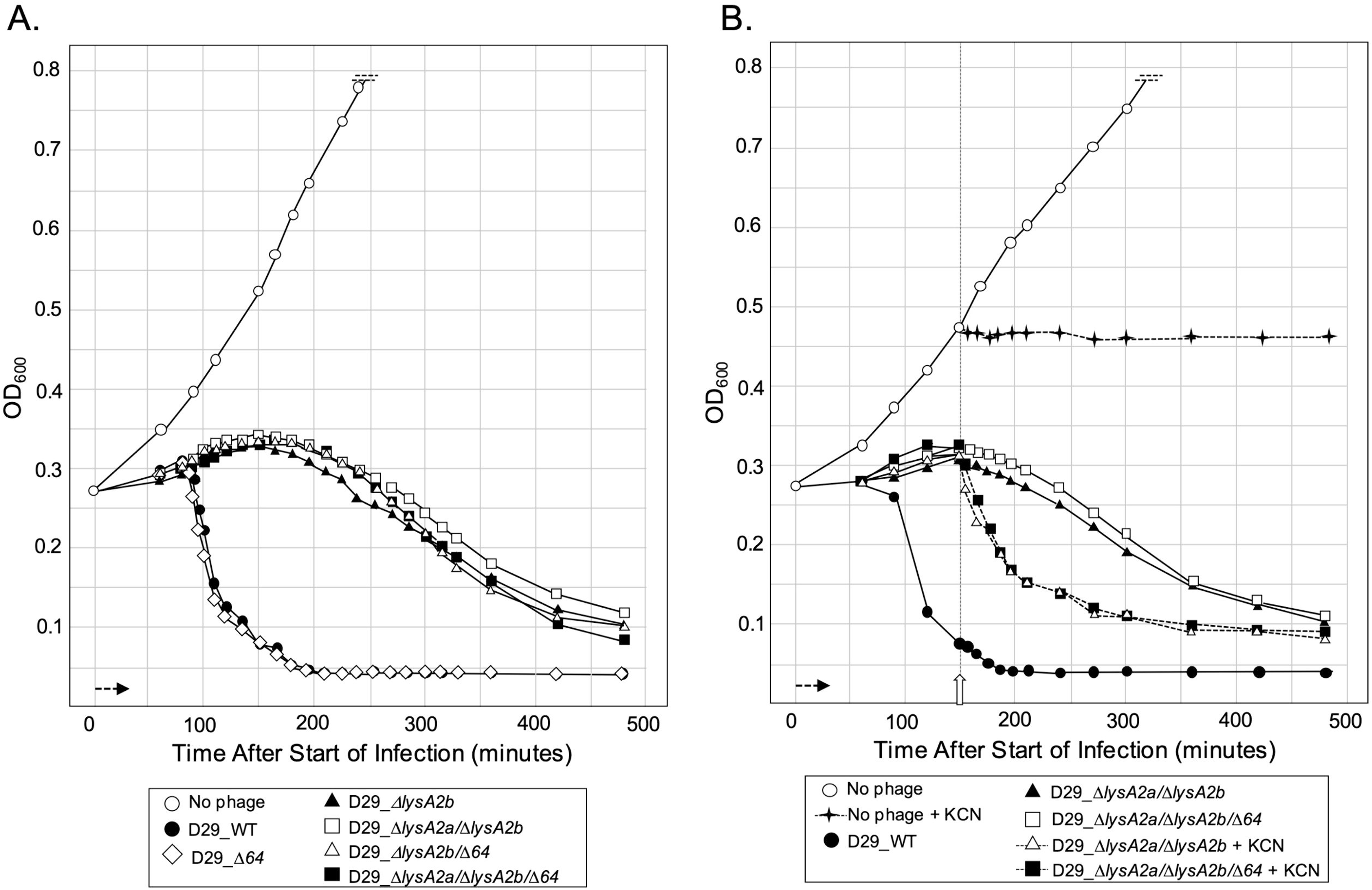
Liquid lysis assay and KCN responsiveness of phages with gene *64* deletions. **A.** At T_0_ a log growth culture of *M. smegmatis* at an OD_600_ of ∼0.25 was infected with the indicated phage at an MOI of 10 and incubated at 37°C for 30 minutes without shaking. Control cultures (no phage) did not receive phage. At T_30_ minutes the culture was constantly shaken a 225rpm, aliquots removed at the indicated times and bacterial growth evaluated at OD_600_. Horizonal arrow shows the 30-minute infection time. B. *M. smegmatis* was infected and propagated as described in A. KCN (final concentration of 10mM) was added to a culture infected with D29_Δ*lysA2b*Δ*64*, D29_Δ*lysA2a*/Δ*lysA2b*/Δ*64* and an uninfected culture at 150 minutes (open vertical arrow).

Since the D29_Δ*lysA2a/*Δ*lysA2b/Δ64* phage was viable and still could be triggered by KCN, it is possible that D29 encodes additional proteins involved in the lysis pathway. Bioinformatic analysis of all D29 gene products revealed three additional TMD proteins: gp38 is a small 50aa lipoprotein, gp59.1 is a small 47aa 1TMD protein and gp34.1 is a 145aa 4TMD protein. S11 Figure 8 in the S2 File shows the genomic location of these genes and predicted AlphaFold models. No LRM mutants have mapped to any of these proteins and exogenous expression of gp34.1, gp38 or gp59.1 does not result in cytotoxicity to *M. smegmatis* in comparison to gp64 (S12 Figure 9 in the S2 File). Extensive analysis with AlpahFold3 also failed to identify any significant interactions of these proteins with LysA2a, LysA2b or gp64. However, we have identified genes encoding 4TMD proteins adjacent to the *lysA2b* gene in cluster A2, A4, A7, A10 and A12 phages (S3 Figure 2A, S4 Figure 2B and S5 Figure 2C in S2 File) that have similar topologies to gp34.1. We have also reported that *Gordonia* phages encode 4TMD proteins within putative lysis cassettes [22]. Therefore, to assess the impact of gp34.1 on the D29 lysis pathway, BRED was utilized to delete *34.1* from D29_Δ*lysA2a*/Δ*lysA2b*. S13 Figure 10 and S14 Figure 11 in the S2 File shows that like the deletion of *64*, loss of *34.1* has no observable impact on plaque size, liquid lysis kinetics or KCN responsiveness compared to D29_Δ*lysA2a*/Δ*lysA2b*.

In summary, deletion of either *64* and *34.1* produce no detectable effects on plaque morphology, lysis kinetics, or KCN responsiveness. However, these experiments identify gp64 as a previously unrecognized lipoprotein capable of genetically bypassing the requirement for both LysA2 proteins. A single C31Y substitution that abolishes the conserved lipobox cysteine and is predicted to convert gp64 from a membrane-anchored lipoprotein into a secreted protein consistently restored efficient lysis of the D29_Δ*lysA2a/*Δ*lysA2b* mutant. Because wild type gp64 and gp64C31Y are both cytotoxic when expressed exogenously, suppression cannot be explained simply by acquisition of toxicity. Instead, the data suggest that the cellular localization of gp64 is critical for its activity. We therefore propose that gp64 is not a core component of the LysA2 regulatory system but rather an accessory envelope protein whose activity is normally constrained by membrane tethering. Disruption of lipidation through the C31Y substitution may release this constraint, allowing gp64 to promote envelope disruption independently of the LysA2 proteins. The pham for gp64 contains genes from ∼400 phages that infect *M. smegmatis*, *Arthrobacter sp*., *Streptomyces sp*., *Rhodococcus sp*., *Gordonia sp*., *Microbacterium sp*. and *Corynbacterium sp*. However, there are no clusters where all the phages in the cluster encode gene gp64 homologs and even in cluster A2 only 40 of the 123 phages have the gene *64* equivalent. Thus, although the molecular target of gp64 remains unknown, its restricted phylogenetic distribution among actinobacteriophages suggests that it represents an accessory function that may enhance lysis efficiency rather than an essential component of the conserved lysis LysA2a/LyA2b regulators. Whether gp34.1, gp38 and gp59.1 also contribute as accessory proteins to lysis pathway will require the generation of a D29 phage with all TMD genes deleted to uncover possible subtle lysis phenotypes.

### The Lys proteins confer a strong competitive fitness advantage during phage propagation

Since the deletion of *lysA2a*, *lysA2b* and gene *64* either individually or in the same phage was not lethal to D29 propagation, it was of interest to assess the fitness of the D29_Δ*lysA2a/ΔlysA2b*/Δ*64* compared to D29_WT. A sample of purified D29_WT and D29_Δ*lysA2a/ΔlysA2b*/Δ*64* was prepared where the concentration of each phage was set to ∼5 x 10^9^ PFU/ml (1:1 stock). *M. smegmatis* was infected with the 1:1 stock at an MOI of ∼2.8 x 10^-6^. The infection was performed at a very low MOI to ensure that nearly all infected cells received only a single phage particle, thereby preventing complementation between wild type and mutant phages during mixed infection. S15 Figure 12 in the S2 File shows the plaque assay and PCR validation of the plaques from the plating of the 1:1 stock. There are nearly equivalent numbers of D29_WT (24) and D29_Δ*lysA2a/ΔlysA2b*/Δ*64* (21) plaques on the plate validating the initial phage concentrations in the 1:1 stock and the ability of the wild type and mutant phages to infect the host at the same efficiency. Phage were then harvested from the plate, filtrates prepared, serial diluted and evaluated by plaque assays as detailed in Methods. In all plaque assays, *M. smegmatis* was infected at low MOI as described for the 1:1 stock. This process was repeated four times as shown in the S16 Figure 13 in the S2 File schematic. After each infection, plates with ∼200-350 plaques were evaluated for the presence of D29_Δ*lysA2a/ΔlysA2b*/Δ*64* by PCR of picked plaques (S17 Figure 14, S18 Figure 15 and S19 Figure 16 in S2 File). In addition, the filtrates were evaluated by DADA PCR to assess the relative ratio of D29_WT to D29_Δ*lysA2a/ΔlysA2b*/Δ*64.* The DADA PCR primer, genomic context, PCR procedure and sensitivity to detect the mutant phage in a wild type background is presented in S2 Figure 1 in S2 File. Importantly, phage with the l*ysA2b* deletion could be detected when the ratio was 1:15,000 to D29_WT. All fitness data is summarized in Table 5 and the DADA analysis of the various filtrates is presented in Figure 16.

**Figure 16.**
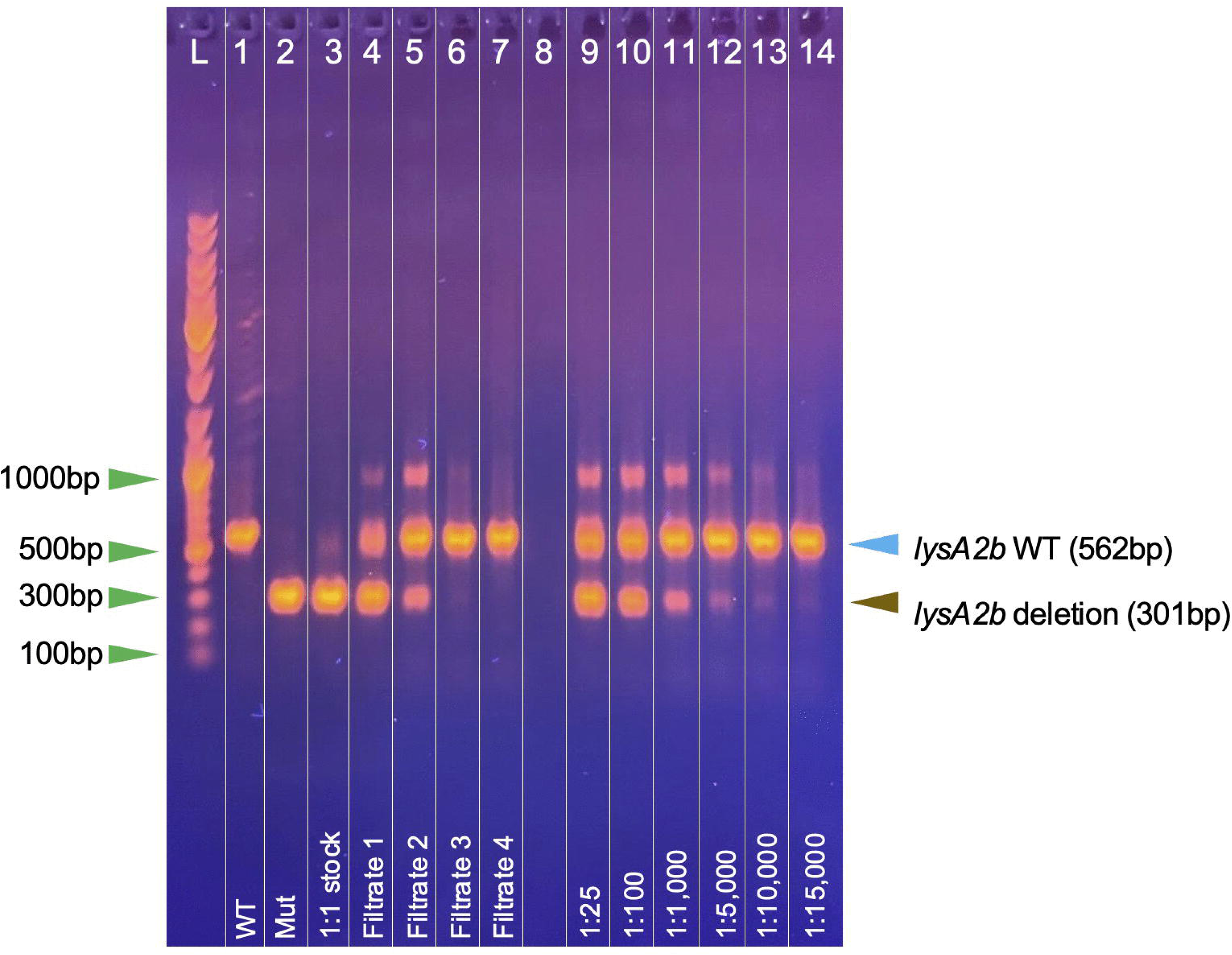
DADA PCR analysis of filtrates from phage fitness assay. The indicated samples were evaluated by DADA PCR using the D29_DADA4Δ*lysA2b* primer as described in Methods. The 1:1 stock represents the sample with an equal ratio of D29_D*lysA2a*/D*lysA2b*/Δ*64:*D29_WT and the generation of the filtrates is detailed in the text and Methods. The 1:25, 1:100, 1:1,000, 1:5,000, 1:10,000 and 1:15,000 samples indicate the ratio of D29_Δ*lysA2a*/Δ*lysA2b*/Δ*64* to D29_WT. WT = control sample of WT_D29. Mut = control sample of D29_Δ*lysA2a*/Δ*lysA2b*/Δ*64*.

It is expected that phages that have the same level of fitness should maintain a 1:1 ratio after multiple rounds of plating and harvesting, while phages that have a fitness disadvantage should become progressively lost from the population. Following a single round of propagation, the ratio of D29_WT to D29_Δ*lysA2a/ΔlysA2b*/Δ*64* increased from 1:1 to approximately 28:1 (S17 Figure 14 in S2 File). After two rounds, the ratio increased to approximately 320:1 (S18 Figure 15 in S2 File), and after three rounds no mutant plaques were identified among the screened isolates (S19 Figure 16 in S2 File). Consistent with these results, DADA PCR demonstrated a progressive decline in the mutant genotype in the harvested filtrates until it was no longer detectable if the filtrate isolated after the fourth round of propagation (Figure 16). These results demonstrate that although deletion of the *lysA2* genes is not lethal, it imposes a substantial competitive fitness cost that results in rapid elimination of the mutant from mixed phage populations, most likely due to restricted phage release from the plaques after lysis compared to D29_WT (Figure 10). Thus, the LysA2 proteins function as major fitness determinants during phage propagation, providing a plausible evolutionary explanation for their widespread conservation among actinobacteriophages despite not being individually essential for viability.

**Table 5.** Summary of fitness data.

| Sample | WT<br>plaques<br>detected<br>by PCR | Mut<br>plaques<br>detected<br>by PCR | WT/Mut Plaque<br>Ratio | WT phage<br>detected by<br>DADA PCR | Mut phage<br>detected by<br>DADA PCR |
| --- | --- | --- | --- | --- | --- |
| 1:1 stock | 22 | 20 | 1.1/1 | detected | detected |
| Filtrate 1 | 142 | 5 | 28.4/1 | detected | detected |
| Filtrate 2 | 320 | 1 | 320/1 | detected | detected |
| Filtrate 3 | 250 | ND | ----- | detected | detected |
| Filtrate 4 | 335 | ND | ----- | detected | not detected |

Phage were picked from plaques and evaluated by PCR as described in Methods. Filtrates were prepared from plates and evaluated by DADA PCR as described in Methods. WT = D29\_WT. Mut = D29\_Δ*lysA2a*/Δ*lysA2b*/Δ64. ND = none detected

## CONCLUSIONS AND IMPLICATIONS

### D29 utilizes two distinct Lys proteins like F1 cluster phages to regulate and trigger host lysis that are localized to different regions of the genome

Although the two Lys proteins encoded by D29 are not in a defined lysis cassette, the findings of the current study replicate many of the key results from our previous report on host lysis by F1 cluster phages and support their role as critical components to phage fitness. Most pertinent is the finding that A2 cluster phage D29 utilizes two distinct lysis regulators to trigger efficient host lysis: the 2TMD LysA2a and the 1TMD LysA2b. These proteins appear to be homologs to the F1 cluster phage lysis regulators, LysF1a and LysF1b, and this is supported by the ability to genetically complement D29_Δ*lysA2b* phage by insertion of the *lysF1b* gene. Importantly, like the Lys proteins from the F1 cluster phages, the 1TMD LysA2b protein can support triggering and liquid lysis independent of the 2TMD LysA2a. Thus, like LysF1a, the LysA2a appears to exist in a non-functional conformation that is activated by the LysA2b. However, based on the isolation and activity of the various LysaA2a LRM mutants, LysA2a is not simply an on/off switch. Due to its extended C-terminal domain, it appears to be a multi-domain regulatory protein in which different structural elements independently influence activation, timing, and overall lytic efficiency that may provide D29 with a fitness advantage.

### All D29 phages with deletions of lysA2a or lysA2b can be triggered by addition on energy poisons possibly due to activation/release of Lysin A

A key finding from the work on F1 cluster phages Girr and NBJ was that only phages with a deletion of the 2TMD *lysF1a* gene could be triggered by the addition of KCN. The inability to trigger lysis in the NBJ_Δ*lysF1a*/Δ*lysF1b* double mutant confirmed that disruption of the PMF by KCN in the NBJ_Δ*lysF1a* phage was due to the presence of the 1TMD LysF1b protein [17]. This data was consistent with the proposed holin-like function of LysF1b especially since the Girr and NBJ encoded Lysin A enzymes lacked any defined secretion signals. The current data shows that KCN triggers D29_Δ*lysA2a*, D29_Δ*lysA2b*, D29_Δ*lysA2a*/Δ*lysA2b* and even D29_Δ*lysA2a*/Δ*lysA2b*/Δ*64.* Thus, disruption of the PMF in D29 triggers lysis independently of the two Lys proteins and gp64. Since the LysA2a and LysA2b generally mimic the functions of the LysF1a and LysA2b proteins on host lysis in the genetic studies, the KCN triggering in the absence of the Lys proteins is suggestive that an additional phage encoded factor is being activated.

One attractive candidate is the Lysin A enzyme. It has long been established that exogenous expression of Lysin A from a select number of actinobacteriophage (that include D29), is highly cytotoxic when expressed exogenously even though there is no evidence for secretion signals or SAR-like domains in these enzymes [48, 67]. We have confirmed this result using the pExTra expression system and show that D29 Lysin A is highly cytotoxic and Girr Lysin A is not (S20 Figure 17 in S2 File). The cytotoxicity of D29 Lysin A is also supported by recent work from the Jain laboratory who also report that exogenously expressed GFP-tagged D29 Lysin A is exported from the cell independently of the predicted lysis regulators and localized to the cell periphery [51, 52]. Together, these observations support the hypothesis that D29 Lysin A is exported prior to lysis and remains in an inactive or constrained state until membrane depolarization [51]. Under this model, KCN would bypass the requirement for the LysA2 proteins by directly promoting PMF-dependent activation of the pre-exported enzyme. In the absence of KCN, spontaneous membrane depolarization during late infection would gradually activate Lysin A, accounting for the slow, protracted lysis observed in the *ΔlysA2b* mutants and the cytotoxicity of the exogenously expressed Lysin A. Furthermore, deletion of the N-terminal catalytic domain of D29 Lysin A (D29^ΔNTD^) results in a protein with inefficient activity toward the peptidoglycan and phage carrying the mutated gene show smaller plaques, prolonged liquid lysis timing and reduced burst size due to compromised phage release [52]. These phenotypes mirror the D29 phage with *lysA2b* deletions that do not effectively trigger lysis and thus, activate the Lysin A. Although D29 Lysin A lacks a canonical SAR domain, this proposed mechanism shares important conceptual features with the *E. coli* phages SAR-endolysin systems, in which membrane depolarization activates an already exported endolysin [68]. In contrast, Girr Lysin A does not produce a strong cytotoxic phenotype when expressed exogenously and also has an extended linker region between the two catalytic domains that may account for this phenotype (S20 Figure 17 in S2 File) [48, 51]. This is consistent with the continued requirement for the Lys proteins to achieve efficient lysis in the F1 cluster phages.

### D29 utilizes a modular lysis network for effective host lysis

Collectively, the findings from the current report expand upon the previous F1 cluster phage data and are supportive of a **modular lysis network** model that is outlined in Table 6.

**Table 6:** Modular Lysis Network Model.

| Module | Evidence | Likely role |
| --- | --- | --- |
| **LysA2a/LysA2b** | genetics, complementation, suppressors | Conserved PMF-responsive lysis regulators that determine lysis timing and disrupt the PMF. May also participate in lesion formation |
| **Lysin A/Lysin B** | KCN triggering, secretion studies, toxicity | Core enzymes that degrade the cell wall components. In some cases, may be exported independent of the Lys Regulators |
| **gp64 and other proteins** | suppressor genetics, localization mutation, toxicity | Accessory envelope factors that modulate lysis efficiency when membrane localization is altered. May interact with other proteins and/or Lys Regulators |

The key aspects of this model are a conserved **regulatory module** (LysA2a/LysA2b) to disrupt the PMF that interfaces with a **non-canonical endolysin pathway** (exported Lysin A, Lysin B) and **accessory proteins** (gp64 and others?) that can genetically bypass the regulators under specific conditions. Since the observed lysis timing and release of phage particles by *M. smegmatis* infected with F1 and A2 cluster phage is generally in excess of 60-90 minutes, it may be the concerted effort and interaction of this suite of proteins that disrupt membrane physiology, generate membrane lesions and ultimately promote the release of phage once the PMF is destroyed. Since nearly all actinobacteriophages annotated to date have versions of the two Lys regulators (Ref 17 and S3 Figure 2A, S4 Figure 2B and S5 Figure 2C in the S2 File), it may be the Lysin A homolog and the accumulation of additional TMD/secreted accessory proteins that provide higher levels of fitness to a given phage. How the Lysin A enzyme is exported from D29 or other actinobacteriophages remains to be determined as does the nature of the membrane lesions. In addition, it will be necessary to generically evaluate additional phages that have diverse lysis gene organizations, Lysin A homologs and versions of gp64 or other TMD proteins to expand and refine this model.

## ACKNOWLEDGMENTS

We are grateful to the members of the Science Education Alliance for their invaluable research support and for supplying phage and plasmid reagents, particularly Dainelle Heller, Viknesh Sivanathan, Graham Hatfull, Deborah Jacobs-Sera, Becky Garlena and Dan Russell. We also thank the Pittsburgh Bacteriophage Institute for genome sequencing several of the phages described in this study. Loc Nguyen is recognized for help with some experiments. Dr. V. Jain (Microbiology and Molecular Biology Laboratory, Department of Biological Sciences, Indian Institute of Science Education and Research) is also acknowledged for kindly supplying D29^Δ11^ phage from his 2020 report [53], although this reagent was not used in any of the figures but for plaque size validation.

