## Supplemental Figures for "Genetic dissection of Mycobacteriophage D29 host lysis reveals two lysis regulators and a novel lipoprotein that regulate the lysis event and are localized to distinct regions of the genome"

**S2 Figure 1. Genomic context of *lysA2b* deletion, DADA PCR procedure and DADA PCR sensitivity to detect *lysA2b* deletion**

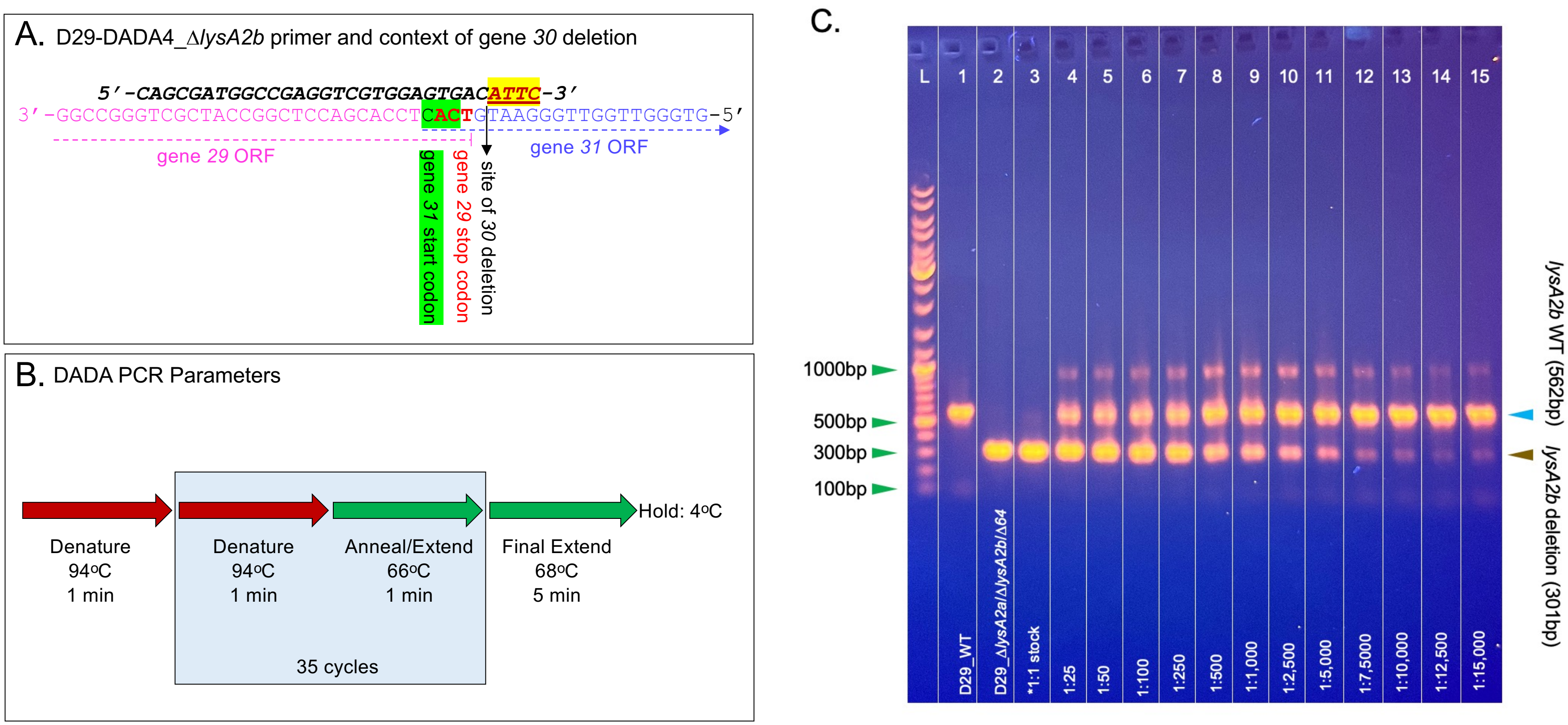

**S2 Figure 1. Genomic context of *lysA2b* deletion, DADA PCR procedure and DADA PCR sensitivity to detect *lysA2b* deletion. A.** The coding strand of the Δ*lysA2b phage* with gene *30* deletion showing the context of the deletion and the GTGA 4bp overlap of gene *29* and gene *31*. The D29-DADA4_Δ*lysA2b* primer with the ATTC overlap to the deleted region highlighted in yellow with red font. **B.** The DADA PCR parameters. **C.** The sensitivity of the DADA PCR procedure to detect the D29_Δ*lysA2a*/Δ*lysA2b*/Δ*64* in the D29_WT background. Samples were prepared with the indicated ratios of D29_Δ*lysA2a*/Δ*lysA2b*/Δ*64*:D29_WT. The *1:1 stock represents the same stock used throughout the fitness assay. The data show that as the ratio of D29_Δ*lysA2a*/Δ*lysA2b*/Δ*64*:D29_WT decreases, the D29_Δ*lysA2a*/Δ*lysA2b*/Δ*64* phage can still be detected even at a ratio of 1:15,000.

**S3 Figure 2A. Identification of genes that encode LysF1a and LysF1b homologs in A cluster phages**

| Cluster | Ref Phage | | | gene | gene | gene | gap | gene | gene | gene | gene | gene | gene | | |
|---|---|---|---|---|---|---|---|---|---|---|---|---|---|---|---|
| | | | | 10 | 11 | 12 | | | | | | 30 | | | |
| **A2** | D29 | | 5' end | LYS A | 2TM | LYS B | ~15kb | tape measure | minor tails | minor tails | NKF | 1TM | minor tails | 3' end | |
| | | | | 296446 | 296091 | 296060 | | | | | | 299040 | | | |
| | | | | 9 | 10 | 11 | | | | | | 30 | | | |
| **A2** | Fameo | | 5' end | LYS A | 2TM | LYS B | ~15kb | tape measure | minor tails | minor tails | NKF | 1TM | minor tails | 3' end | |
| | | | | 296446 | 235559 | 296060 | | | | | | 297967 | | | |
| | | | | 12 | 13 | 14 | | | | | | 33 | | | |
| **A17** | 40AC | | 5' end | LYS A | 2TM | LYS B | ~15kb | tape measure | minor tails | minor tails | NKF | 1TM | minor tails | 3' end | |
| | | | | 296446 | 298590 | 296060 | | | | | | 297967 | | | |
| | | | | 12 | 14 | 10 | | | | | | 30 | | | |
| **A21** | Kinmap | | 5' end | LYS A | 2TM | LYS B | ~15kb | tape measure | minor tails | minor tails | NKF | 1TM | minor tails | 3' end | |
| | | | | 298320 | 296091 | 296060 | | | | | | 297967 | | | |
| | | | | 12 | 13 | 14 | | | | | | 32 | 33 | | |
| **A2** | ArcherNM | | 5' end | LYS A | 2TM | LYS B | ~15kb | tape measure | minor tails | minor tails | NKF | 4TM | 1TM | minor tails | 3' end |
| | | | | 296446 | 298590 | 296060 | | | | | | 235123 | 235138 | | |
| | | | | 9 | 10 | 11 | | | | | | 30 | 31 | | |
| **A7** | DroogsArmy | | 5' end | LYS A | 2TM | LYS B | ~15kb | tape measure | minor tails | minor tails | NKF | 4TM | 1TM | minor tails | 3' end |
| | | | | 296446 | 298590 | 296060 | | | | | | 235123 | 235138 | | |
| | | | | 11 | 12 | 13 | | | | | | 31 | 32 | | |
| **A12** | Steamy | | 5' end | LYS A | 2TM | LYS B | ~15kb | tape measure | minor tails | minor tails | NKF | 4TM | 1TM | minor tails | 3' end |
| | | | | 296094 | 298590 | 296060 | | | | | | 235123 | 235138 | | |
| | | | | 8 | 9 | 10 | | | | | | 29 | 30 | | |
| **A4** | Cici | | 5' end | LYS A | 2TM | LYS B | ~15kb | tape measure | minor tails | minor tails | NKF | 4TM | 1TM | minor tails | 3' end |
| | | | | 296094 | 296091 | 296060 | | | | | | 235123 | 235138 | | |

| | |
|---|---|
| 10 | gene number |
| LYS A | gene function |
| 296446 | pham number |
| LYS A | Lysin A |
| LYS B | Lysin B |
| 1TM | protein w/1TMD |
| 2TM | protein w/2TMD |
| 4TM | protein w/4TMD |
| NKF | unknown function |
| | terminase gene |

**S3 Figure 2A. Identification of genes that encode LysF1a and LysF1b homologs in A cluster phages**. Reference phages (Ref Phage) in the clusters listed in the table were analyzed for the presence of genes for *tape measure protein, minor tails, lysin A, lysin B* and associated genes encoding TMD proteins. All TMD proteins were evaluated for TMD domains and topology as described in Methods. The schematic is not to scale for size and simply shows the relative location of the annotated lysis genes in relation to the *tape measure protein, minor tails* and genes encoding TMD proteins. The order of the genes in relation to the lysis A are exact, with genes of unknown function labeled as NKF. The order of any TMD genes downstream of the *tape measure* and *minor tails* is exact. All genes encoding proteins predicted to have 1TMD or 2TMD that are homologs to the LysF1a and LysF1b proteins are identified. All genes with the same functional call or number of TMDs that are in the same color are grouped to the same pham and the pham number is shown below the gene. Gene number in the Ref phage is shown above the gene. The intent of the schematic is to show that the genes encoding the Lysin enzymes are in the 3' arm of the A cluster phages while all A cluster phages contain 1-3 TMD genes downstream of the *tape measure* and minor tail genes ~15,000bp from the Lysins. A key to the gene designations is shown.

**S4 Figure 2B. Identification of genes that encode LysF1a and LysF1b homologs in A cluster phages**

| Cluster | Ref Phage | | | *gene* | *gene* | *gene* | *gap* | *gene* | *gene* | *gene* | *gene* | *gene* | *gene* | | |
|---|---|---|---|---|---|---|---|---|---|---|---|---|---|---|---|
| | | | | *10* | *11* | | | | | | | *31* | *32* | | |
| **A1** | Acme | | *5' end* | **LYS A** | **LYS B** | *terminase* | ~15kb | *tape measure* | *minor tails* | *minor tails* | **NKF** | **2TM** | **1TM** | *minor tails* | *3' end* |
| | | | | *296094* | *296060* | | | | | | | *84700* | *217564* | | |
| | | | | *10* | *11* | | | | | | | *30* | *31* | | |
| **A3** | Caviar | | *5' end* | **LYS A** | **LYS B** | *terminase* | ~15kb | *tape measure* | *minor tails* | *minor tails* | **NKF** | **2TM** | **1TM** | *minor tails* | *3' end* |
| | | | | *294363* | *296060* | | | | | | | *84700* | *297967* | | |
| | | | | *9* | *10* | | | | | | | *26* | *27* | | |
| **A19** | Kimona | | *5' end* | **LYS A** | **LYS B** | *terminase* | ~15kb | *tape measure* | *minor tails* | *minor tails* | **NKF** | **2TM** | **1TM** | *minor tails* | *3' end* |
| | | | | *294363* | *296060* | | | | | | | *84700* | *297967* | | |
| | | | | *7* | *8* | *9* | | | | | | *27* | *28* | | |
| **A5** | Chadwick | | *5' end* | **LYS A** | **NKF** | **LYS B** | ~15kb | *tape measure* | *minor tails* | *minor tails* | **NKF** | **2TM** | **1TM** | *minor tails* | *3' end* |
| | | | | *296219* | *125376* | *296060* | | | | | | *84700* | *297967* | | |
| | | | | *7* | *8* | | | | | | | *26* | *27* | | |
| **A8** | Dixon | | *5' end* | **LYS A** | **LYS B** | *terminase* | ~15kb | *tape measure* | *minor tails* | *minor tails* | **NKF** | **2TM** | **1TM** | *minor tails* | *3' end* |
| | | | | *296219* | *296060* | | | | | | | *84700* | *297967* | | |
| | | | | *6* | *7* | | | | | | | *25* | *26* | | |
| **A5** | Zolita | | *5' end* | **LYS A** | **LYS B** | *terminase* | ~15kb | *tape measure* | *minor tails* | *minor tails* | **NKF** | **2TM** | **1TM** | *minor tails* | *3' end* |
| | | | | *298242* | *296060* | | | | | | | *84700* | *297967* | | |
| | | | | *9* | *10* | | | | | | 29 | *30* | *31* | | |
| **A10** | Chupacabra | | *5' end* | **LYS A** | **LYS B** | *terminase* | ~15kb | *tape measure* | *minor tails* | **NKF** | **2TM** | **4TM** | **1TM** | *minor tails* | *3' end* |
| | | | | *294363* | *296060* | | | | | | *106665* | *235123* | *235138* | | |
| | | | | *11* | *12* | | | | | | | | *31* | | |
| **A12** | DarthPhader | | *5' end* | **LYS A** | **2TM** | *terminase* | ~15kb | *tape measure* | *minor tails* | *minor tails* | *minor tails* | NKF | **1TM** | *minor tails* | *3' end* |
| | | | | *296094* | *298590* | | | | | | | | *297967* | | |

| | |
|---|---|
| *10* | gene number |
| **LYS A** | gene function |
| *296446* | pham number |
| **LYS A** | Lysin A |
| **LYS B** | Lysin B |
| **1TM** | protein w/1TMD |
| **2TM** | protein w/2TMD |
| **4TM** | protein w/4TMD |
| **NKF** | unknown function |
| | *terminase gene* |

**S4 Figure 2B. Identification of genes that encode LysF1a and LysF1b homologs in A cluster phages**. Reference phages (Ref Phage) in the clusters listed in the table were analyzed for the presence of genes for *tape measure protein, minor tails, lysin A, lysin B* and associated genes encoding TMD proteins. All TMD proteins were evaluated for TMD domains and topology as described in Methods. The schematic is not to scale for size and simply shows the relative location of the annotated lysis genes in relation to the *tape measure protein, minor tails* and genes encoding TMD proteins. The order of the genes in relation to the lysis A are exact, with genes of unknown function labeled as NKF. The order of any TMD genes downstream of the *tape measure* and *minor tails* is exact. All genes encoding proteins predicted to have 1TMD or 2TMD that are homologs to the LysF1a and LysF1b proteins are identified. All genes with the same functional call or number of TMDs that are in the same color are grouped to the same pham and the pham number is shown below the gene. Gene number in the Ref phage is shown above the gene. The intent of the schematic is to show that the genes encoding the Lysin enzymes are in the 3' arm of the A cluster phages while all A cluster phages contain 1-3 TMD genes downstream of the *tape measure* and minor tail genes ~15,000bp from the Lysins. A key to the gene designations is shown.

### S5 Figure 2C. Identification of genes that encode LysF1a and LysF1b homologs in A cluster phages

| Cluster | Ref Phage | | | gene | gene | gene | gap | gene | gene | | gene | gene | gene | |
|---|---|---|---|---|---|---|---|---|---|---|---|---|---|---|
| | | | | 11 | 12 | | | | | | | 32 | | |
| **A16** | EagleEye | | 5' end | LYS A | 2TM | terminase | ~15kb | tape measure | minor tails | minor tails | NKF | 1TM | minor tails | 3' end |
| | | | | 296446 | 298590 | | | | | | | 297967 | | |
| | | | | 13 | 14 | | | | | | | 33 | | |
| **A6** | CookieDough | | 5' end | LYS A | 2TM | terminase | ~15kb | tape measure | minor tails | minor tails | NKF | 1TM | minor tails | 3' end |
| | | | | 296094 | 296091 | | | | | | | 297967 | | |
| | | | | 7 | 8 | | | | | | | 28 | | |
| **A18** | MyraDee | | 5' end | LYS A | 2TM | terminase | ~15kb | tape measure | minor tails | minor tails | NKF | 1TM | minor tails | 3' end |
| | | | | 296094 | 296091 | | | | | | | 297967 | | |
| | | | | 10 | 11 | | | | | | | 30 | | |
| **A11** | Aneem | | 5' end | LYS A | 2TM | terminase | ~15kb | tape measure | minor tails | minor tails | NKF | 1TM | minor tails | 3' end |
| | | | | 298320 | 296091 | | | | | | | 297967 | | |
| | | | | 9 | 10 | | | | | | | 30 | | |
| **A14** | Jeeves | | 5' end | LYS A | 2TM | terminase | ~15kb | tape measure | minor tails | minor tails | NKF | 1TM | minor tails | 3' end |
| | | | | 298320 | 296091 | | | | | | | 297967 | | |
| | | | | 10 | 11 | | | | | | | 31 | | |
| **A22** | Chargerpower | | 5' end | LYS A | 2TM | terminase | ~15kb | tape measure | minor tails | minor tails | NKF | 1TM | minor tails | 3' end |
| | | | | 298320 | 296091 | | | | | | | 297967 | | |
| | | | | 10 | 11 | | | | | | | 30 | | |
| **A9** | Charm | | 5' end | LYS A | 2TM | terminase | ~15kb | tape measure | minor tails | minor tails | NKF | 1TM | minor tails | 3' end |
| | | | | 298320 | 296091 | | | | | | | 297967 | | |
| | | | | 6 | 7 | | | | | | | 25 | | |
| **A13** | Phlei | | 5' end | LYS A | 2TM | terminase | ~15kb | tape measure | minor tails | minor tails | NKF | 1TM | minor tails | 3' end |
| | | | | 294363 | 296091 | | | | | | | 297967 | | |
| | | | | 10 | 11 | | | | | | | 29 | | |
| **A20** | Anthony | | 5' end | LYS A | 2TM | terminase | ~15kb | tape measure | minor tails | minor tails | NKF | 1TM | minor tails | 3' end |
| | | | | 296446 | 299631 | | | | | | | 297967 | | |

| | |
|---|---|
| 10 | gene number |
| LYS A | gene function |
| 296446 | pham number |
| LYS A | Lysin A |
| LYS B | Lysin B |
| 1TM | protein w/1TMD |
| 2TM | protein w/2TMD |
| 4TM | protein w/4TMD |
| NKF | unknown function |
| | terminase gene |

**S5 Figure 2C. Identification of genes that encode LysF1a and LysF1b homologs in A cluster phages**. Reference phages (Ref Phage) in the clusters listed in the table were analyzed for the presence of genes for *tape measure protein, minor tails, lysin A, lysin B* and associated genes encoding TMD proteins. All TMD proteins were evaluated for TMD domains and topology as described in Methods. The schematic is not to scale for size and simply shows the relative location of the annotated lysis genes in relation to the *tape measure protein, minor tails* and genes encoding TMD proteins. The order of the genes in relation to the lysis A are exact, with genes of unknown function labeled as NKF. The order of any TMD genes downstream of the *tape measure* and *minor tails* is exact. All genes encoding proteins predicted to have 1TMD or 2TMD that are homologs to the LysF1a and LysF1b proteins are identified. All genes with the same functional call or number of TMDs that are in the same color are grouped to the same pham and the pham number is shown below the gene. Gene number in the Ref phage is shown above the gene. The intent of the schematic is to show that the genes encoding the Lysin enzymes are in the 3' arm of the A cluster phages while all A cluster phages contain 1-3 TMD genes downstream of the *tape measure* and minor tail genes ~15,000bp from the Lysins. A key to the gene designations is shown.

**S6 Figure3. Liquid lysis kinetics and KCN triggering of D29_Δ*lysA2a***

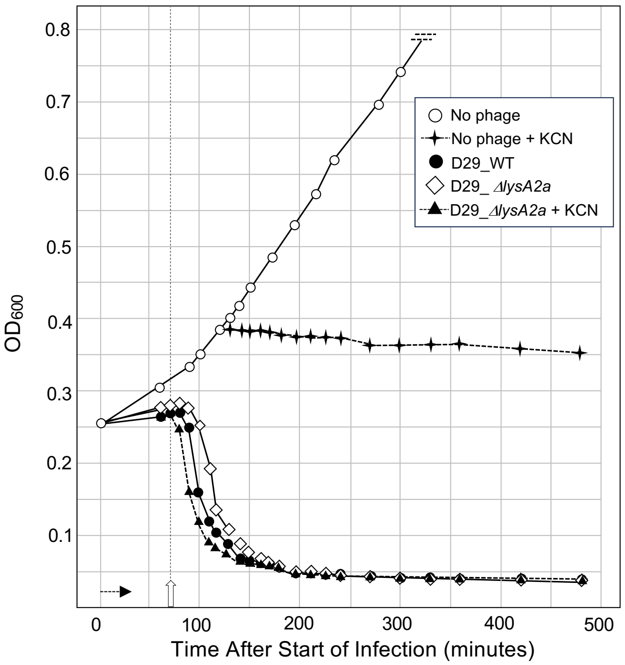

**S6 Figure 3. Liquid lysis kinetics and KCN triggering of D29_Δ*lysA2a*.** At $T_0$ a log growth culture of *M. smegmatis* at an $OD_{600}$ of ~0.25 was infected with the indicated phage at an MOI of 10 and incubated at 37°C for 30 minutes without shaking. Control cultures (no phage) did not receive phage. At $T_{30}$ minutes the culture was constantly shaken a 225rpm, aliquots removed at the indicated times and bacterial growth evaluated at $OD_{600}$. Horizonal arrow shows the 30-minute infection time. KCN (final concentration of 10mM) was added to a culture infected with D29_Δ*lysA2b* and an uninfected culture at 150 minutes (open vertical arrow).

**S7 Figure 4. Generation and validation of Girr_Δ46 phage**

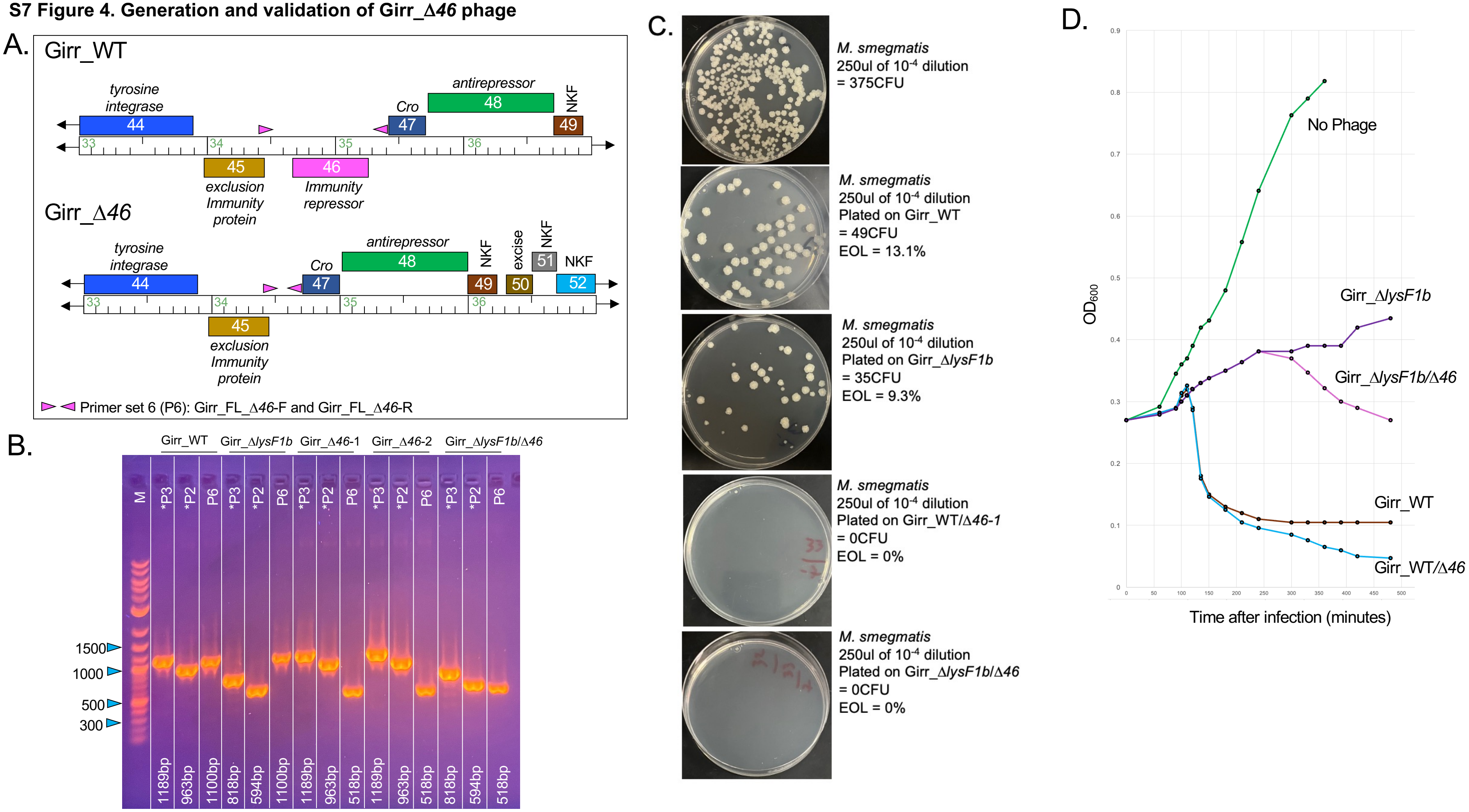

**S7 Figure 4. Generation and validation of Girr_Δ*46* and Girr_Δ*lysF1b*/Δ*46* phage.**

**A.** Genomic structure and PCR primer set location for deletion of gene *46* (*immunity repressor*). Gene functions of genes flanking *46* are indicated. NKF = hypothetical protein with no known function.

**B.** PCR verification of Δ*46* phages. The indicated phages were evaluated using primer primer set 6 (S1 Table 1 in the S1 File) and uses primers Girr_FL_Δ*46* (F and R) and generates a 1100bp fragment in Girr_WT and 518bp with *46* deletion. Primers sets *P2 and *P3 are from the F1 cluster phage report [17] and allow detection of the *lysF1b* gene. All fragments correspond to the expected sizes and validate the deletion of gene *46* and *lysF1b* in the Girr_Δ*lysF1b*/Δ*46* phage.

**C.** Efficiency of lysogeny (EOL) of Girr_WT, Girr_Δ*lysF1b*, Girr_Δ*46* and Girr_Δ*lysF1b*/Δ*46*. 100ul of 1 x $10^9$ stocks of the indicated phages were spread on 7H10 agar and covered with 250ul of a 1 x $10^{-4}$ dilution of saturated *M. smegmatis*. One plate did not receive phage and served as the control for colony growth. Plates were propagated at 37$^o$C for 4 days and colonies counted. EOL = number of colonies on phage plates/number of colonies on no phage plates. The data show that Girr phages with an intact immunity repressor (gene *46)* have an EOL of 9-13% in this experiment consistent with data reported previously [17] and for F1 phage NBJ [36]. Girr_Δ*46* and Girr_Δ*lysF1b*/Δ*46* have an EOL of 0% and are fully lytic.

**D.** Liquid lysis of Girr_WT, Girr_Δ*lysF1b*, Girr_Δ*46* and Girr_Δ*lysF1b*/Δ*46.* At $T_0$ a log growth culture of *M. smegmatis* at an $OD_{600}$ of ~0.25 was infected with the indicated phage at an MOI of 10 and incubated at 37$^o$C for 30 minutes without shaking. Control cultures (no phage) did not receive phage. At $T_{30}$ minutes the culture was constantly shaken a 225rpm, aliquots removed at the indicated times and bacterial growth evaluated at $OD_{600}$. Horizonal arrow shows the 30-minute infection time. Girr_Δ*46* and Girr_Δ*lysF1b*/Δ*46* show greater reductions in $OD_{600}$ compared to the phages that express gene *46*.

**S8 Figure 5. Generation and validation of D29_Δ*lysA2b*::Girr_*lysF1b***

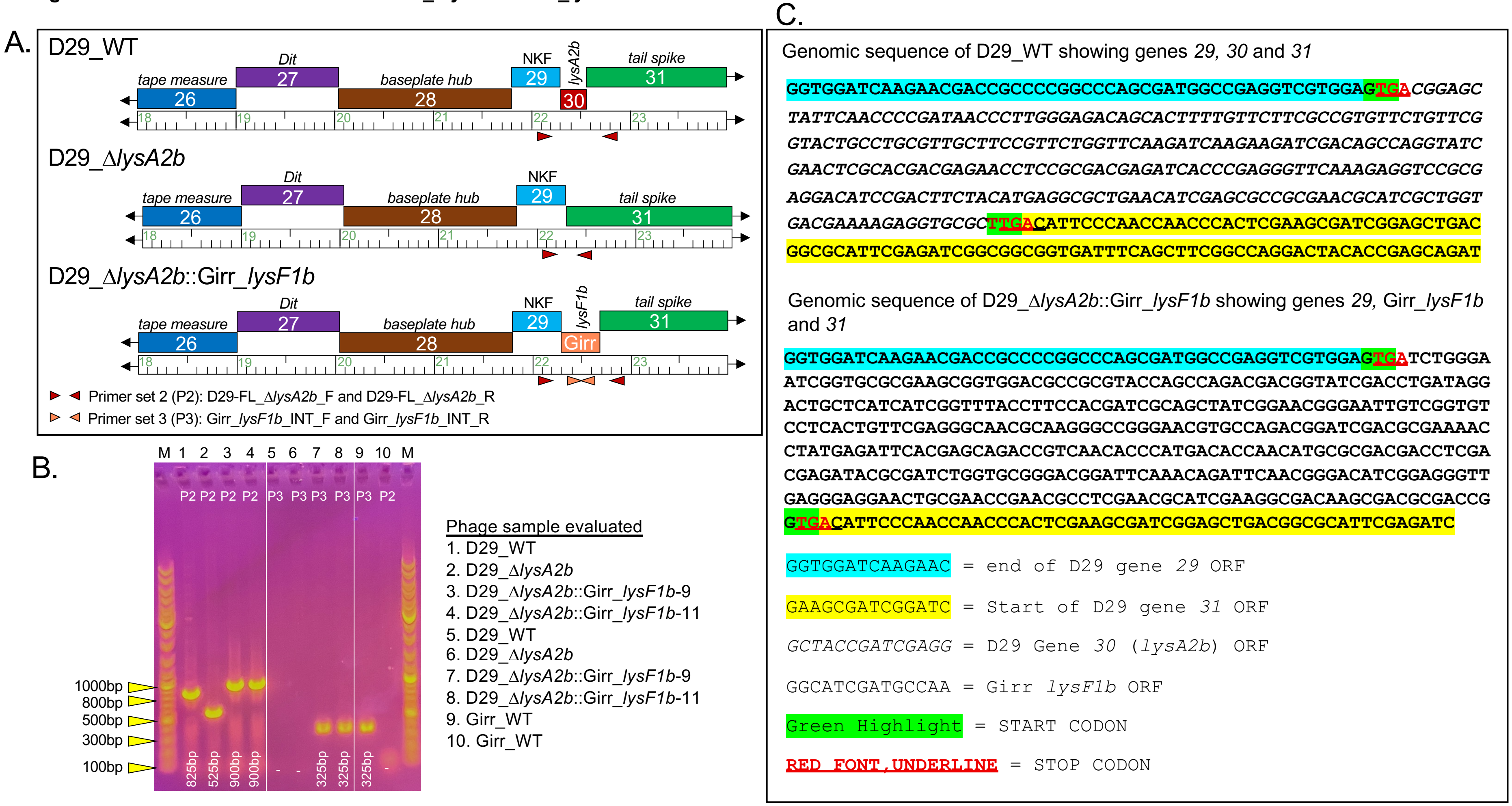

**S8 Figure 5. Generation and validation of D29_Δ*lysA2b::*Girr_*lysF1b***

**A.** Genomic structure and PCR primer set location for insertion of Girr *lysF1b into D29_ΔlysA2b*. Gene functions of genes flanking *lysA2b* are indicated. NKF = hypothetical protein with no known function.

**B.** PCR verification of phages. The indicated phages were evaluated using primer sets 2 and 3 that are listed in the S1 Table 1 in the S1 File. Primer set 2 uses primers D29-FL_Δ*lysA2b* (F and R) and generates a 825bp fragment in D29_WT, a 525bp fragment in D29_Δ*lysA2b* and a 900bp fragment with the Girr *lysF1b* insertion. Primer set 3 uses internal primers to Girr *lysF1b* and generates a 325bp fragment if *lysF1b* is present. All PCR fragments correspond to the expected sizes and validate the insertion of the *lysF1b* gene.

**C.** Genomic context of the *lysF1b* insertion. **Top sequence:** the *lysA2b* open reading frame in D29 has a 4bp GTGA overlap to gene *29* and TTGA overlap to gene *31*. The Girr *lysF1b* open reading frame has 4bp GTGA overlap to gene *34* and GTGA overlap to gene *36* ([31]; PhagesDB: https://phagesdb.org/phages/Girr ). **Bottom sequence:** the Girr *lysF1b* gene was inserted between D29 *29* and *31* and maintains its original GTGA overlaps as shown.

### S9 Figure 6. Generation and validation of D29 phages with gene *64* deletion

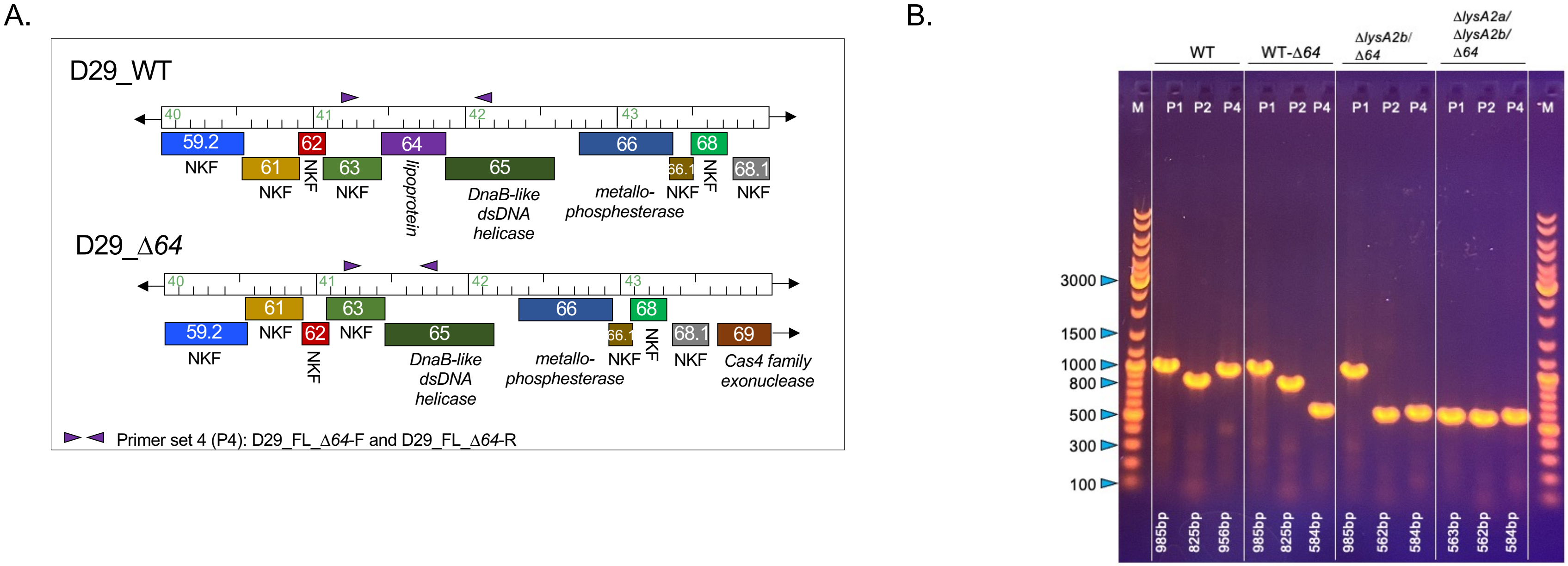

**S9 Figure 6. Generation and validation of D29 phages with gene *64* deletion. A.** Genomic structure and PCR primer set location for deletion of gene *64*. Functions of genes flanking *64* are indicated. NKF = hypothetical protein with no known function. **B.** PCR verification of phages. The indicated phages were evaluated using primer sets 1, 2 and 4 that are listed in the S1 Table 1 in the S1 File. Primer sets 1 and 2 are detailed in Figure 4 and primer set 6 uses primers D29-FL_Δ*64* (F and R) and generates a 956bp fragment in D29_WT and a 584bp fragment in when gene *64* is deleted. All PCR fragments correspond to the expected sizes and validate the deletion of gene *64*.

**S10 Figure 7. Plaque sizes in phages with gene *64* deletion.**

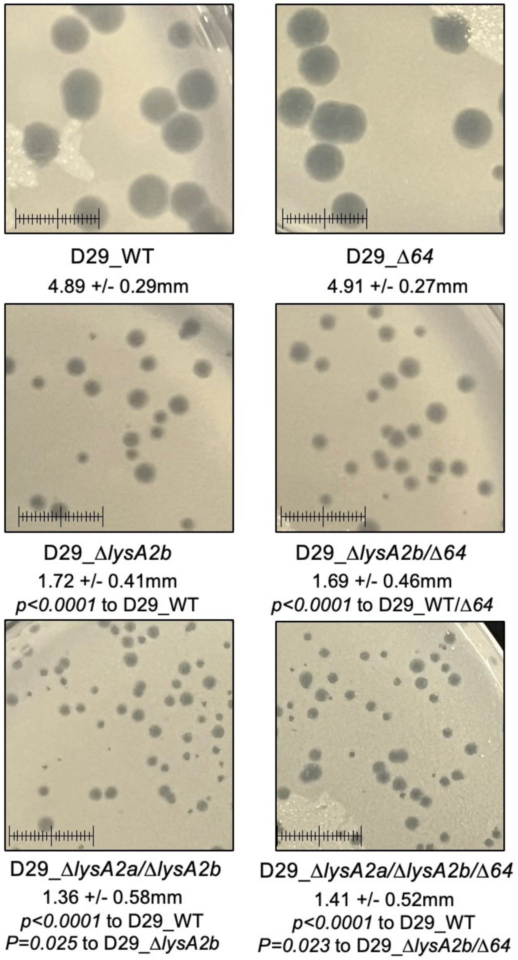

**S10 Figure 7. Plaque sizes in phages with gene *64* deletion.** *M. smegmatis* was infected with identical numbers of the indicated phage as detailed in Methods and plaque size determined after 36 hrs. of growth at 37$^{o}$C. Average diameter +/- SD for a minimum of 20 plaques is shown. Statistical significance was determined as detailed in Methods. Scale = 1cm.

**S11 Figure 8. Identification of D29 genes encoding TMD proteins and AlphaFold modeling.**

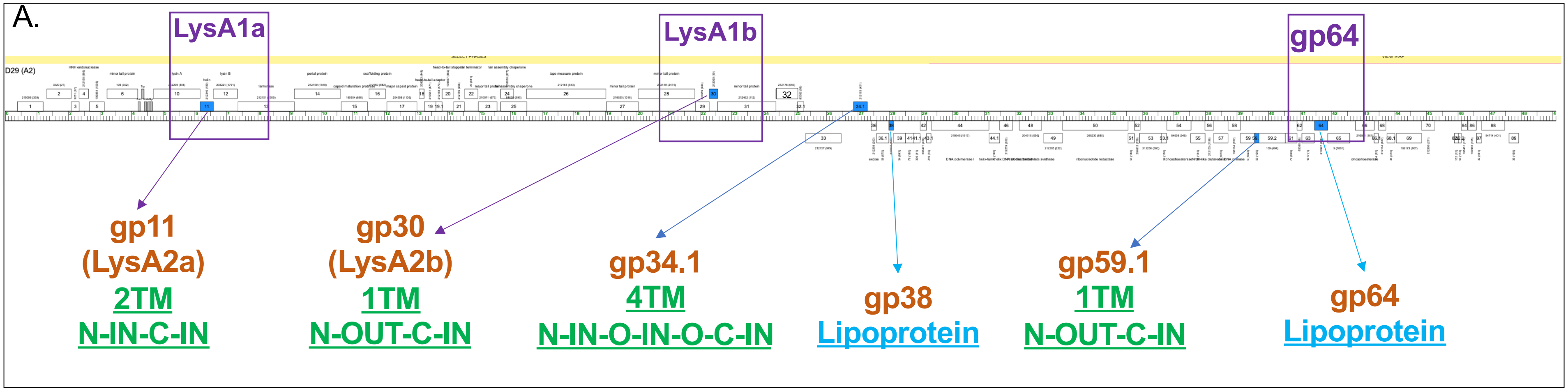

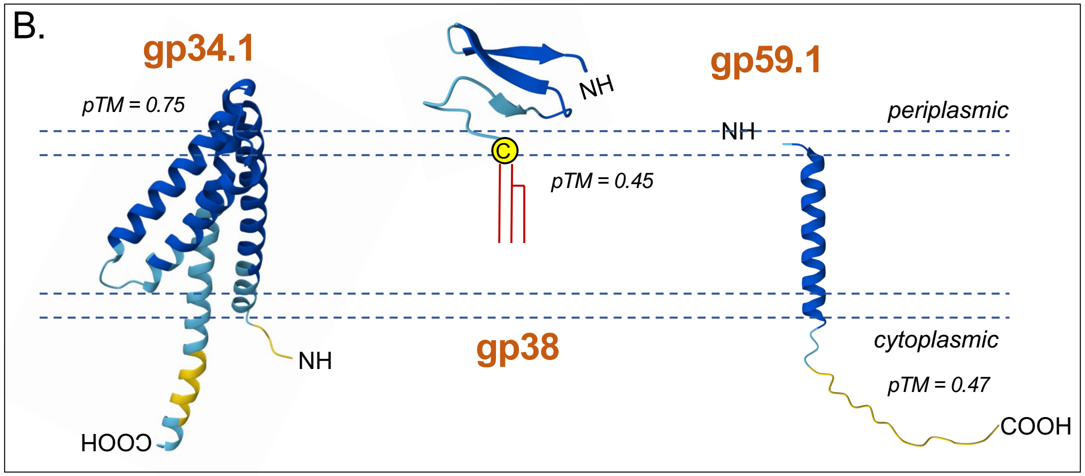

**S11 Figure 8. Identification of D29 genes encoding TMD proteins and AlphaFold modeling. A.** The genomic location of all encoded TMD proteins in D29 are shown in relation to the Lysis regulators and gp64. **B.** The indicated proteins were modeled using AlpahFold3 as described in Methods. The predicted topology of each protein was determined as described in Methods. The predicted orientation of each protein in the lipid bilayer is shown. NH = N-terminus. COOH = C-terminus. The conserved cysteine residue, lipidation and periplasmic location of gp38 is shown. pTM scores for each model are shown.

**S12 Figure 9. Cytotoxicity analysis of D29 TMD proteins**

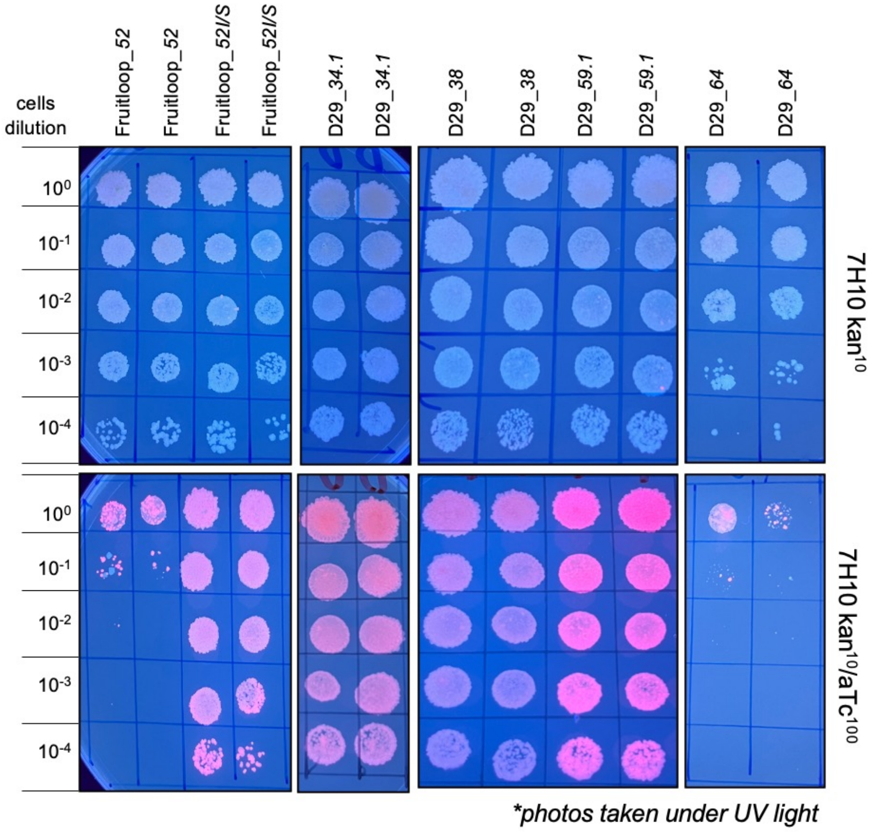

**S12 Figure 9. Cytotoxicity analysis of D29 TMD proteins.** The indicated genes were cloned into pExTra01 as described in Methods. The resulting plasmids were transfected into *M. smegmatis* and individual colonies with the specified pExTra plasmid was resuspended in 7H9, serially diluted, and spotted on 7H10 Kan agar containing 0 or 100 ng/mL aTc and incubated at 37ºC as detailed in Methods. Plates were inverted and photographed under UV light to visualize the mCherry expression. Fruitloop_*52* serves as the positive cytotoxicity control and the Fruitloop_*52I/S* serves as the negative control [31, 32].

**S13 Figure 10. Generation and validation of D29_Δ*lysA2a*/Δ*lysA2a*/Δ*34.1*.**

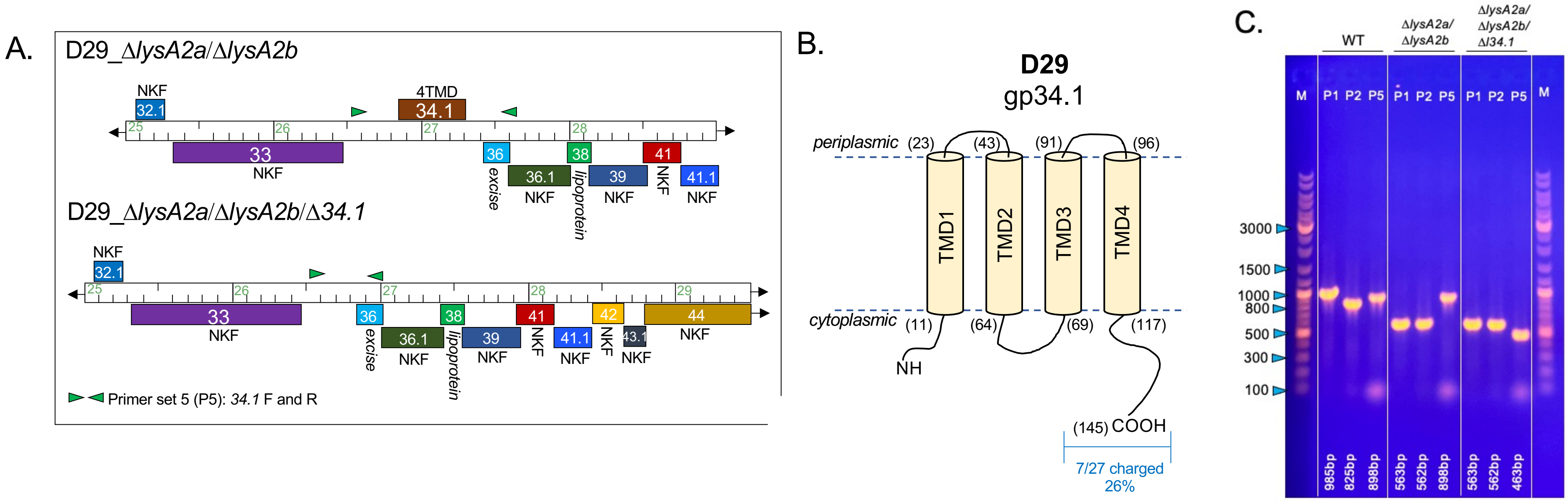

**S13 Figure 10. Generation and validation of D29_Δ*lysA2a*/Δ*lysA2b*/Δ*34.1*.**

**A.** Genomic structure and PCR primer set location for deletion of D29 *34.1*. Functions of genes flanking *34.1* are indicated. NKF = hypothetical protein with no known function.

**B.** Predicted topology of gp34.1. The amino acid sequence of 34.1 was evaluated for TMDs and topology as described in Methods. The gp34.1 protein was predicted to have 4 TMDs. Numbers in parenthesis indicate the amino acid number. NH = N-terminus. COOH = C- terminus.

**C.** PCR verification of phages. The indicated phages were evaluated using primer sets 1, 2 and 5 that are listed in the S1 Table 1 in the S1 File. Primer sets 1 and 2 are detailed in Figure 4 and primer set 5 uses primers D29-FL_Δ*34.1* (F and R) and generates an 898bp fragment in D29_WT and a 463bp fragment when gene *34.1* is deleted. All PCR fragments correspond to the expected sizes and validate the deletion of gene *34.1*.

**S14 Figure 11. Analysis of D29_Δ*lysA2a*/Δ*lysA2b*/Δ*34.1* plaques and liquid lysis kinetics**

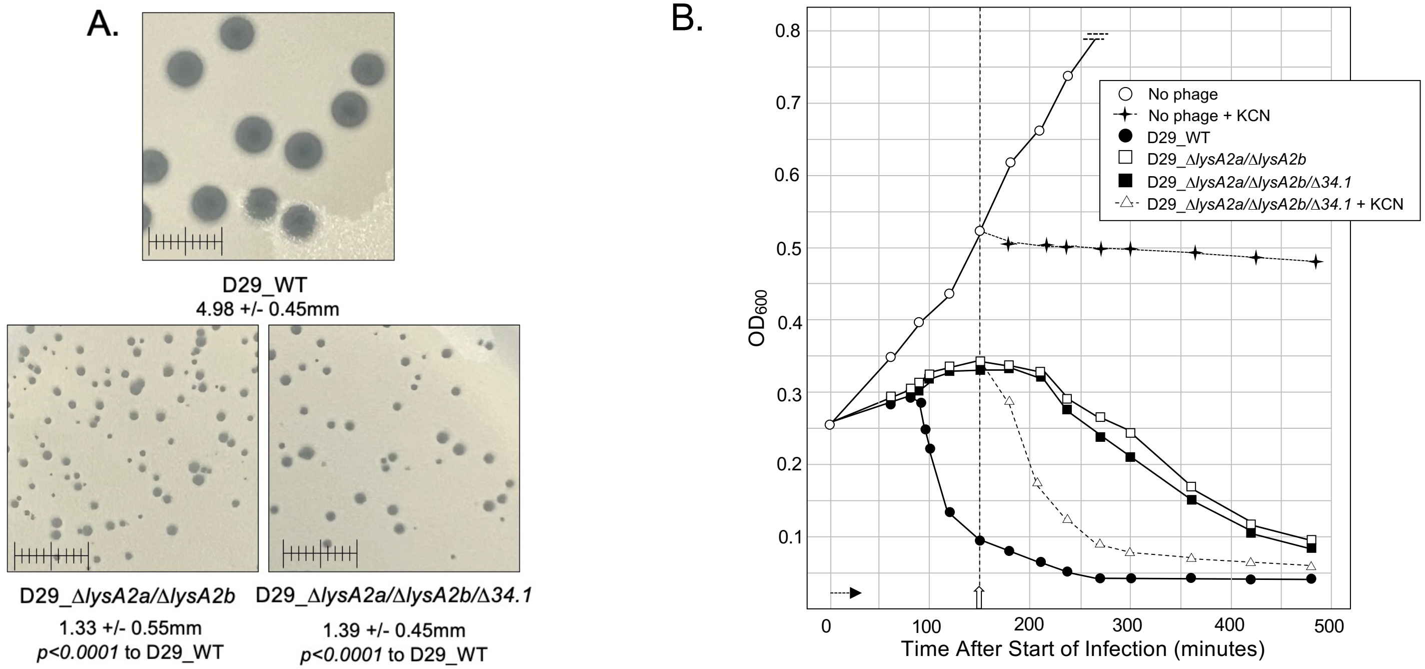

**S14 Figure 11. Analysis of D29_Δ*lysA2a*/Δ*lysA2b*/Δ*34.1* plaques and liquid lysis kinetics. A.** *M. smegmatis* was infected with identical numbers of the indicated phage as detailed in Methods and plaque size determined after 36 hrs. of growth at 37ºC. Average diameter +/- SD for a minimum of 20 plaques is shown. Statistical significance was determined as detailed in Methods. Scale = 1cm. B. At $T_0$ a log growth culture of *M. smegmatis* at an $OD_{600}$ of ~0.25 was infected with the indicated phage at an MOI of 10 and incubated at 37ºC for 30 minutes without shaking. Control cultures (no phage) did not receive phage. At $T_{30}$ minutes the culture was constantly shaken a 225rpm, aliquots removed at the indicated times and bacterial growth evaluated at $OD_{600}$. Horizonal arrow shows the 30-minute infection time. KCN (final concentration of 10mM) was added to a culture infected with D29_Δ*lysA2a*/Δ*lysA2b*/Δ*34.1* and an uninfected culture at 150 minutes (open vertical arrow).

**S15 Figure 12. Analysis of 1:1 stock plaque assay**

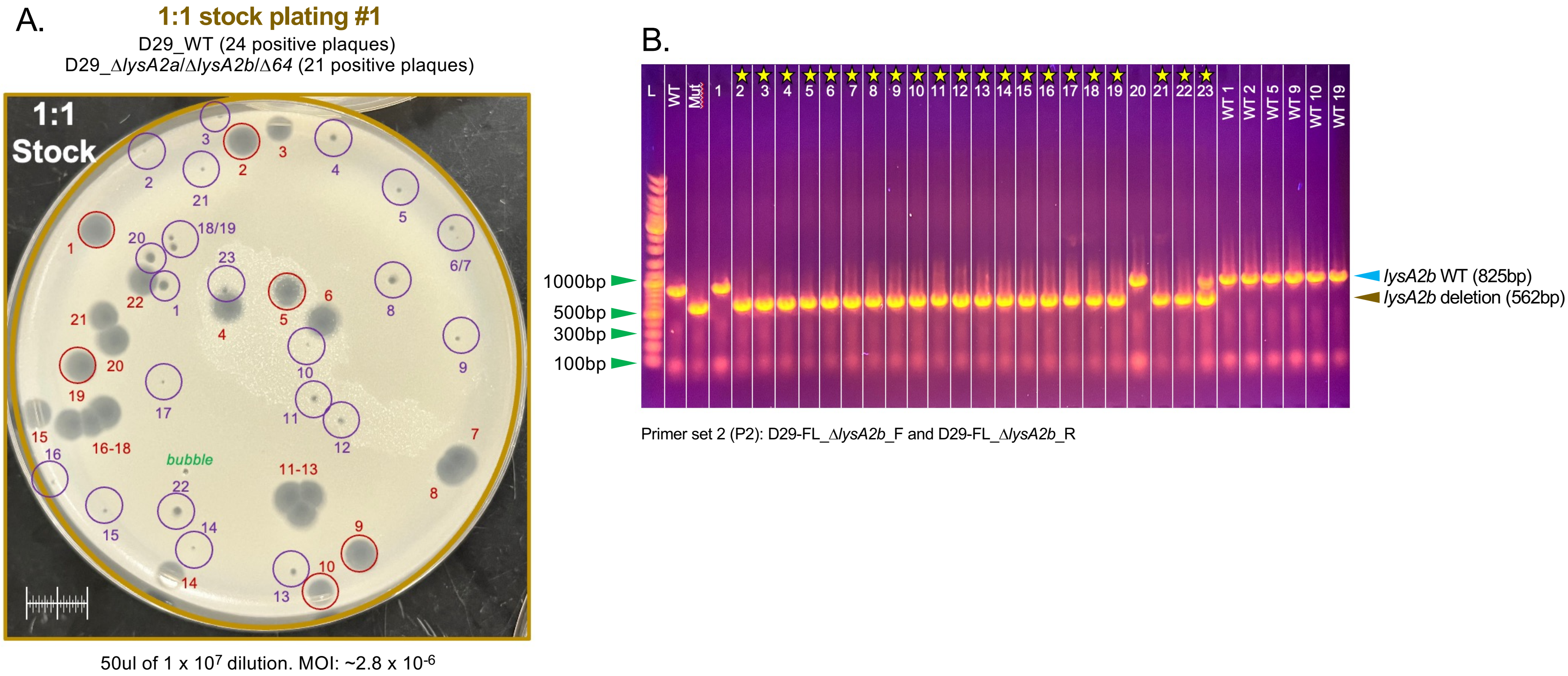

**S14 Figure 11. Analysis of 1:1 stock plaque assay. A.** 50ul of the 1 x $10^7$ dilution from the 1:1 stock was mixed with *M. smegmatis* and plaque assays performed as detailed in Methods. MOI = ~2.8 x $10^{-6}$. All plaques on the plate were picked into 120ul PB and then evaluated by PCR using primer set 2 to amplify the *lysA2b* gene region. **B.** Results of the PCR analysis of the indicated circled and numbered plaques in panel A. Although all plaques were screened, six of the large plaques positive for the D29_WT phage are shown for comparison (WT1, WT2, WT5, WT9, WT10, WT19). Stars identify the plaques generated by the D29_Δ*lysA2a*/Δ*lysA2b*/Δ*64* phage. Due to the proximity of plaque 23 to plaque 4, both D29_WT and D29_Δ*lysA2a*/Δ*lysA2b*/Δ*64* were detected. WT = control PCR using D29_WT. Mut = control PCR using D29_Δ*lysA2a*/Δ*lysA2b*/Δ*64.*

**S16 Figure 13. Workflow of the Fitness Assay showing the plates flooded to prepare the filtrates**

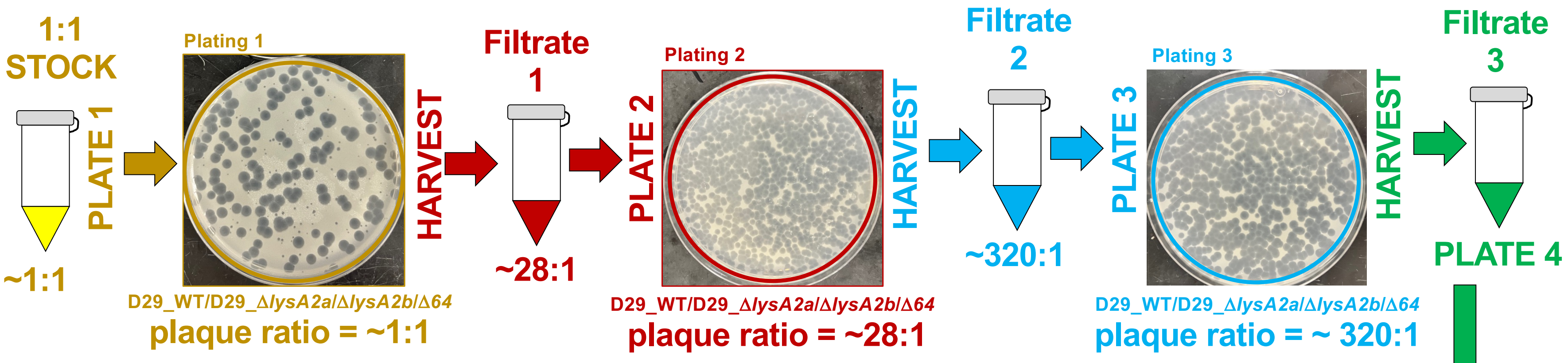

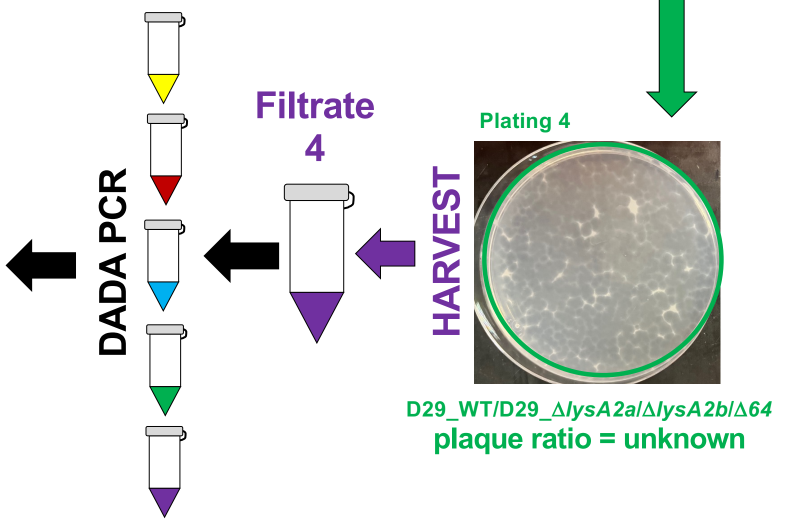

**S16 Figure 13. Workflow of the Fitness Assay showing the plates flooded to prepare the filtrates.** The 1:1 stock was serial diluted and plaque assays performed as described in Methods. Plates with well separated plaques were evaluated by PCR to verify the ratio of D29_WT to D29_*ΔlysA2a/ΔlysA2b/Δ64* and the plate shown was flooded with PB and Filtrate #1 prepared as described in Methods. Filtrate #1 was serial diluted and plaque assays performed. Plates with well separated plaques were evaluated by PCR to verify the ratio of D29_WT to D29_*ΔlysA2a/ΔlysA2b/Δ64* and the plate shown was flooded with PB and Filtrate #2 prepared. Filtrate #2 was serial diluted and plaque assays performed. Plates with well separated plaques were evaluated by PCR to verify the ratio of D29_WT to D29_*ΔlysA2a/ΔlysA2b/Δ64* and the plate shown was flooded with PB and Filtrate #3 prepared. Filtrate #3 was serial diluted and plaque assays performed. Plates with well separated plaques were evaluated by PCR to verify the ratio of D29_WT to D29_*ΔlysA2a/ΔlysA2b/Δ64* and the plate shown was flooded with PB and Filtrate #4 prepared. The ratio of D29_WT: D29_*ΔlysA2a/ΔlysA2b/Δ64* was determined by PCR analysis of the plates presented in S16 Figure 13, S17 Figure 14 and S18 Figure 15 in the S2 File . The 1:1 stock and Filtrates were evaluated by DADA PCR as described in Methods.

**S17 Figure 14. Analysis of Filtrate #1 plaque assay**

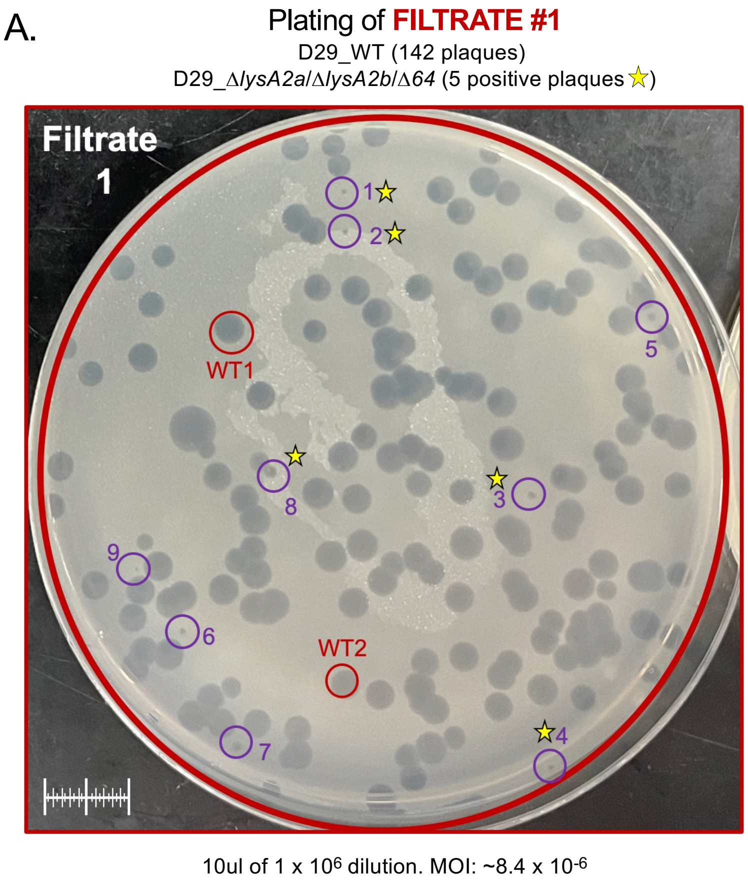

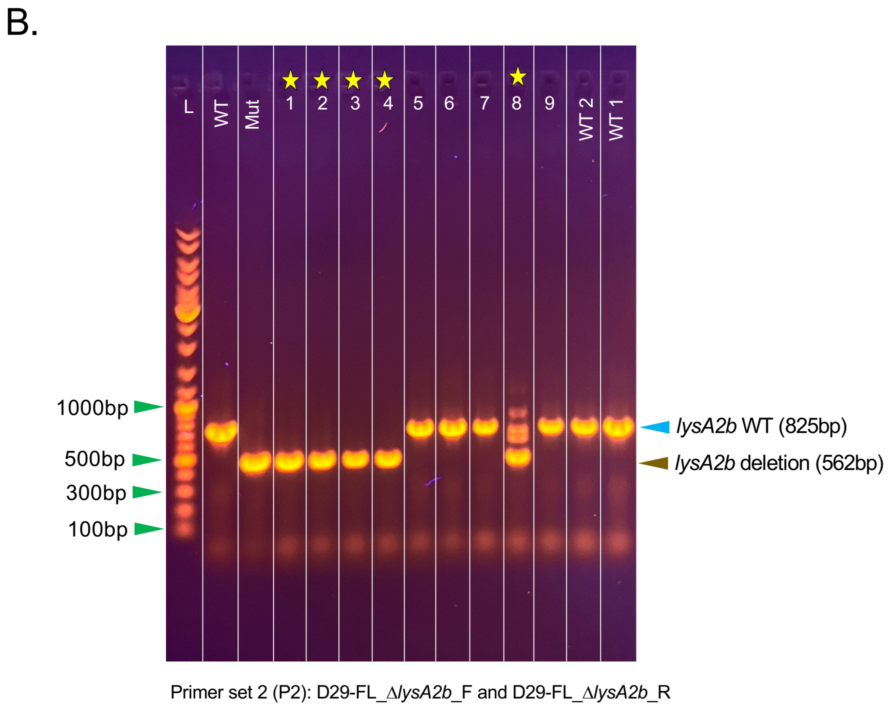

**S17 Figure 14. Analysis of Filtrate #1 plaque assay. A.** 10ul of the 1 x $10^6$ dilution from Filtrate #1 was mixed with *M. smegmatis* and plaque assays performed as detailed in Methods. MOI = ~8.4 x $10^{-6}$. All plaques on the plate were picked into 120ul PB and then evaluated by PCR using primer set 2 to amplify the *lysA2b* gene region. Stars identify plaques positive for D29_*ΔlysA2a/ΔlysA2b/Δ64.* **B.** Results of the PCR analysis of the indicated circled and numbered plaques in panel A. Although all plaques were screened, only two of the large plaques positive for the D29_WT phage are shown for comparison (WT1, WT2). Stars idenify plaques generated by the D29_*ΔlysA2a/ΔlysA2b/Δ64* phage. WT = control PCR using D29_WT. Mut = control PCR using D29_*ΔlysA2a/ΔlysA2b/Δ64.*

**S18 Figure 15. Analysis of Filtrate #2 after plaque assay**

A.

Plating of FILTRATE #2

D29_WT (320 plaques)

D29 Δ*lysA2a*/Δ*lysA2b*/Δ*64* (1 positive plaque ☆)

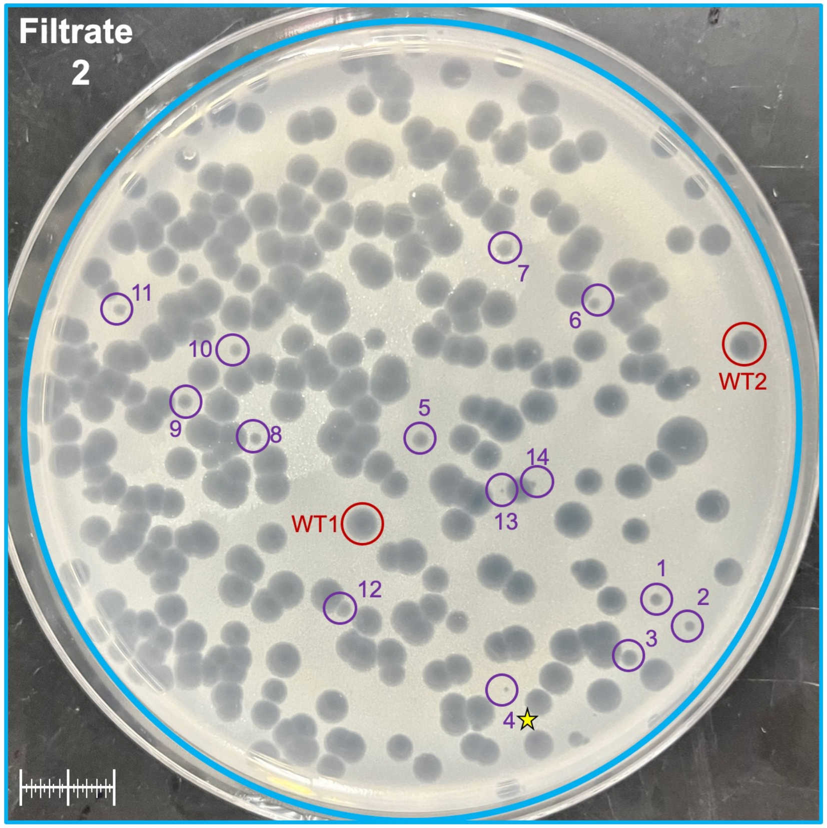

25ul of 1 x $10^6$ dilution. MOI: ~1.8 x $10^{-5}$

B.

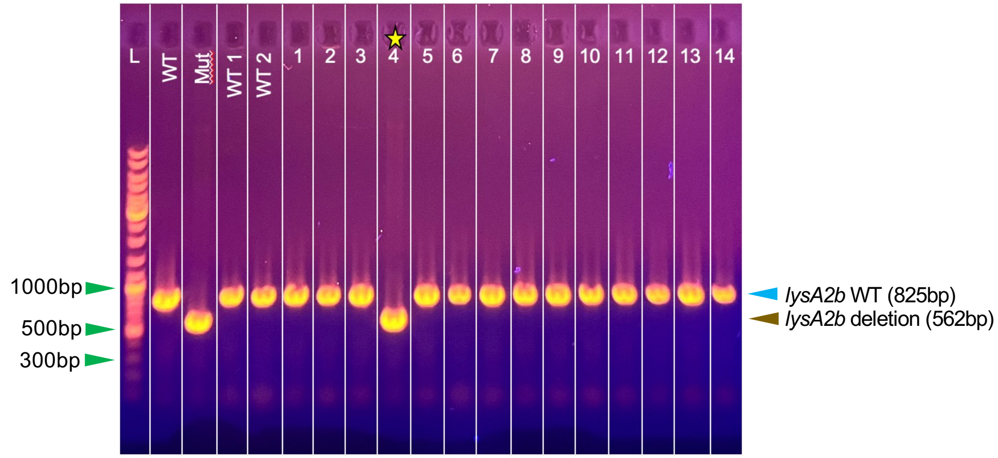

Primer set 2 (P2): D29-FL_Δ*lysA2b*_F and D29-FL_Δ*lysA2b*_R

**S18 Figure 15. Analysis of Filtrate #2 after plaque assay. A.** 25ul of the 1 x $10^6$ dilution from Filtrate #1 was mixed with *M. smegmatis* and plaque assays performed as detailed in Methods. MOI = ~1.8 x $10^{-5}$. All plaques on the plate were picked into 120ul PB and then evaluated by PCR using primer set 2 to amplify the *lysA2b* gene region. The star identifies the one plaque positive for D29_Δ*lysA2a*/Δ*lysA2b*/Δ*64.* **B.** Results of the PCR analysis of the indicated circled and numbered plaques in panel A. Although all plaques were screened, only two of the large plaques positive for the D29_WT phage are shown for comparison (WT1, WT2). Stars indicate plaques generated by the D29_Δ*lysA2a*/Δ*lysA2b*/Δ*64* phage. WT = control PCR using D29_WT. Mut = control PCR using D29_Δ*lysA2a*/Δ*lysA2b*/Δ*64.*

**S19 Figure 16. Analysis of Filtrate #3 after plaque assay**

Plating of FILTRATE #3

A.

D29_WT (250 plaques)

D29_Δ*lysA2a*/Δ*lysA2b*/Δ*64* (0 positive plaques)

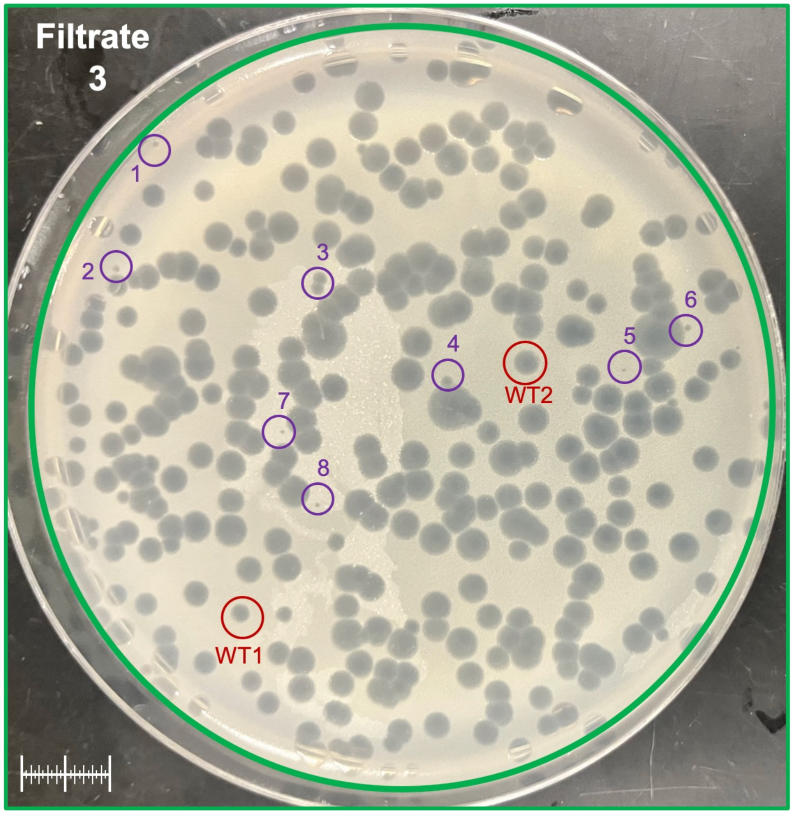

25ul of 1 x $10^6$ dilution. MOI: ~1.8 x $10^{-5}$

B.

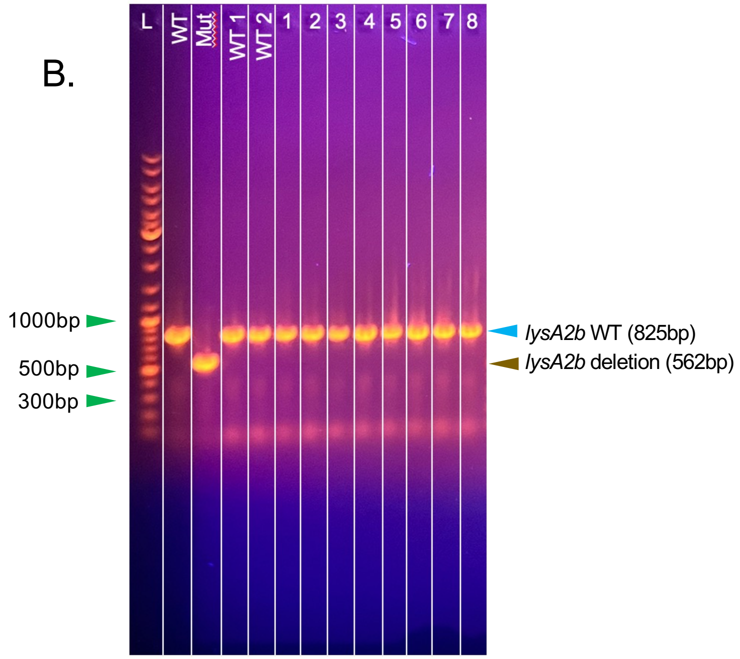

Primer set 2 (P2): D29-FL_Δ*lysA2b*_F and D29-FL_Δ*lysA2b*_R

**S19 Figure 16. Analysis of Filtrate #3 after plaque assay. A.** 25ul of the 1 x $10^6$ dilution from Filtrate #1 was mixed with *M. smegmatis* and plaque assays performed as detailed in Methods. MOI = ~1.8 x $10^{-5}$. All plaques on the plate were picked into 120ul PB and then evaluated by PCR using primer set 2 to amplify the *lysA2b* gene region. **B.** Results of the PCR analysis of the indicated circled and numbered plaques in panel A. Although all plaques were screened, only two of the large plaques positive for the D29_WT phage are shown for comparison (WT1, WT2). There were no plaques that were positive for the D29_Δ*lysA2a*/Δ*lysA2b*/Δ*64* phage. WT = control PCR using D29_WT. Mut = control PCR using D29_Δ*lysA2a*/Δ*lysA2b*/Δ*64.*

**S20 Figure 19. AlphaFold modeling of Lysin A and cytotoxicity assay**

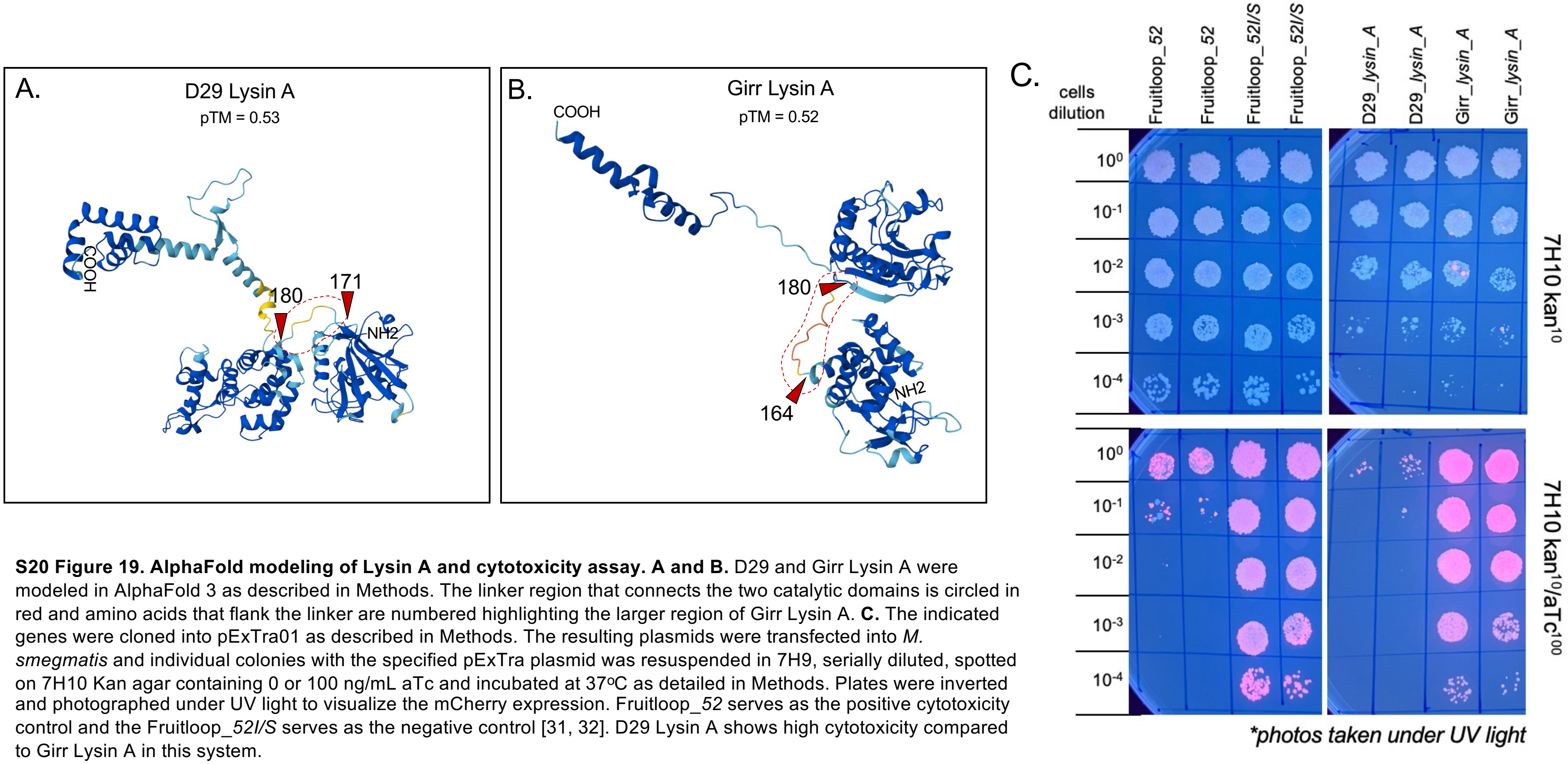

**S20 Figure 19. AlphaFold modeling of Lysin A and cytotoxicity assay. A and B.** D29 and Girr Lysin A were modeled in AlphaFold 3 as described in Methods. The linker region that connects the two catalytic domains is circled in red and amino acids that flank the linker are numbered highlighting the larger region of Girr Lysin A. **C.** The indicated genes were cloned into pExTra01 as described in Methods. The resulting plasmids were transfected into *M. smegmatis* and individual colonies with the specified pExTra plasmid was resuspended in 7H9, serially diluted, spotted on 7H10 Kan agar containing 0 or 100 ng/mL aTc and incubated at 37ºC as detailed in Methods. Plates were inverted and photographed under UV light to visualize the mCherry expression. Fruitloop_*52* serves as the positive cytotoxicity control and the Fruitloop_*52I/S* serves as the negative control [31, 32]. D29 Lysin A shows high cytotoxicity compared to Girr Lysin A in this system.
